# The adapt-to-nutrient NRPS-like secondary metabolite gene cluster facilitates *Verticillium dahliae* adaptation to different nutrient environments

**DOI:** 10.1101/2025.10.20.683378

**Authors:** Ying-Yu Chen, Lea S. Steglich, Nadja Spasovski, Marcel H. W. Franzius, Merle Aden, Isabel Maurus, Rebekka Harting, Gerhard H. Braus

## Abstract

Filamentous fungi produce a wide range of secondary metabolites to adapt to changing environments. RNA sequencing revealed that nine biosynthetic gene clusters (BGCs) of the phytopathogenic *Verticillium dahliae* react to different nutrient environments. The <u>a</u>dapt-to-<u>n</u>utrient <u>N</u>RPS-like (*ANN*) cluster contributes to antibacterial activity and developmental processes important for the early biotrophic life cycle, but is dispensable for virulence on tomato (*Solanum lycopersicum*). Transcription of the core biosynthetic enzyme-encoding *ANN3* is highly induced in nutrient-poor environment. *ANN3* is transcriptionally controlled by global and in-cluster transcription factors. *ANN3* is activated by early colonisation transcription factors Som1 and Vta2, but repressed by Mtf1, which governs late stages of disease progression. The in-cluster transcription factor Ann1, which represses *ANN3*, is less stable in nutrient-poor environment or when *V. dahliae* encounters antagonists. Ann1 promotes resting structure formation but suppresses conidiation and antibacterial activity. Possible products of the *ANN* cluster were revealed by comparing metabolites extracted from *ANN3* regulator mutants and from the bacterial-fungal interaction zone. Our findings revealed that *V. dahliae* perceives different nutrient environments and changes its survival strategy by differential expression of the *ANN* secondary metabolite gene cluster.

**Author summary:** *Verticillium dahliae* is an economically significant phytopathogen that is widely distributed. The fungus adjusts and adapts its survival strategy according to the surrounding environment. Transcriptome data revealed that the core biosynthetic gene *ANN3* of the <u>a</u>dapt-to-<u>n</u>utrient <u>N</u>RPS-like (*ANN*) cluster is most expressed in nutrient-poor environments. The expression of *ANN3* is governed by in-cluster repressor Ann1 and global transcriptional regulators that regulate other metabolic processes. Transcription factors Som1 and Vta2 are involved in the early plant-root infection process, whereas Mtf1 regulates late stage of disease development. *ANN3* is activated by Som1 and Vta2, but repressed by Mtf1. The repressor Ann1 is less stable in nutrient-poor environments or when bacterial competitors are present. Ann1 promotes dormancy and represses spreading by conidiation. Vegetative growth is reduced but antibacterial activity is promoted when *ANN1* is deleted. Possible chemical products of the *ANN* cluster were identified by comparing the metabolites extracted from the regulator mutant strains and the bacterial-fungal interaction zone. In summary, our findings show how *V. dahliae* react to environmental signals to balance growth, survival, and competition through the *ANN* cluster.

## Introduction

*Verticillium dahliae* is a haploid soil-born plant pathogen that infects a wide range of plant hosts (1). Verticillium wilt, the monocyclic plant disease caused by *V. dahliae* infection, results in significant losses in agriculture worldwide (1). The control of the fungus is difficult due to its resistant resting structures known as microsclerotia, and the rapid spread within the plant vasculature system by asexual conidiospores (1). Heavily melanised microsclerotia are typically released into the environment from decaying plant material, and the dormancy of *Verticillium* can persist for years (2). Environmental conditions such as favourable temperature, moisture levels, and chemical signals produced by soil bacteria as well as plant root exudates induce germination from microsclerotia (3–5). The *ex planta* life cycle following dormancy is relatively short. Upon arrival at its plant host, the fungus adheres, colonises, and penetrates the root surface (6, 7). As *V. dahliae* crosses through the cortex and enters the xylem vessels, it produces conidiospores to cause systemic infection (1). Along with the senescence of the plant host, *V. dahliae* transitions to saprophytic growth, and forms microsclerotia, which are subsequently released into the environment along with decaying plant materials (1).

*V. dahliae* encounters antagonists including different soil bacteria in the rhizosphere during the short saprophytic *ex planta* phase (5, 8). Filamentous fungi protect themself and outcompete other microbes mainly by chemical defence strategies (9). Secondary metabolites (SMs) are molecules that are derived from primary metabolic pathways. These natural products do not directly contribute to survival, but are rather produced under specific conditions to increase fitness. For example, DHN-melanin is produced during *V. dahliae* microsclerotia formation, and lagopodin B is synthesised by *Coprinopsis cinerea* when it’s confronted with bacteria (10, 11). For the ease of regulation, genes involved in the synthesis of a specific SM are often located in close proximity within the genome, thus forming biosynthetic gene clusters (BGCs) (12). The increased availability of whole genome sequences and bioinformatic algorithms made studies on cryptic or uncharacterised BGCs possible, leading to the discovery of previously unknown natural products (13). Transcriptional regulation of BGCs takes place at several levels. Whereas global transcriptional regulators or protein complexes may act on multiple metabolic processes, some BGCs also contain cluster-specific transcription factors (14).

Unlike many other filamentous fungi, previous observations indicate that *V. dahliae* prefers to physically escape antagonists by changing growth directions and reducing metabolic activities (8). Whereas more studies focused on the inhibition of *Verticillium* spp. growth by soil bacteria, recent studies suggest that the fungus can also chemically defend itself from competitors (8, 15–17). To date, understanding towards the metabolite producing potential of *V. dahliae* remains elusive. Only three BGCs in the *V. dahliae* genome have been studied. The melanin biosynthesis *PKS2* (also published as *PKS1*) cluster is regulated by in-cluster transcription factors Vta1 and Cmr1 (11, 18–20), the local transcription factor Nag1-controlled *HYB1* cluster is involved in developmental processes and melanin biosynthesis (19, 21), whereas the nonribosomal peptide synthetases in the *NRPS1* cluster contributes to the regulation of developmental processes and pathogenicity towards plants (22).

Transition from the fungal *ex planta* to the *in planta* life cycle is controlled by a series of sequentially acting transcription factors. The *V. dahliae* Som1 regulator is involved in the control of adherence and penetration of root surfaces and corresponds to *S. cerevisiae FLO8* that activates adherence and flocculation. The *Verticillium* transcription activator of adhesion 3 (Vta3) controls the colonisation of root surfaces, and subsequently Vta2 is involved in the fungal proliferation within roots (6, 7). The Som1-Vta3-Vta2 regulatory network coordinates physiological processes to adapt to different phases of the *V. dahliae* life cycle and its corresponding nutrient environment (23, 24). Transcription factors active during early phases of infection can promote regulatory subnetworks that also drive the onset of plant infection, such as the activation of *VTA3* by Som1, and the increased expression of *VTA2* by Som1 and Vta3 (6, 7). On the contrary, Vta3 and Vta2 decrease the expression of the Mtf1-regulated subnetwork that is involved in later phases of disease development (23).

In this study, we describe that the presence of a nutrient-poor environment activates the <u>a</u>dapt-to-<u>n</u>utrient <u>N</u>RPS-like cluster (*ANN* cluster, formerly termed *OTHER3* cluster (19)) in *V. dahliae*. The in-cluster transcription factor Ann1 is a transcriptional repressor of the core biosynthetic enzyme-encoding gene *ANN3*. Protein stability of Ann1 is affected by the surrounding nutrient environment. *ANN3* expression can be controlled by global transcription factors that are involved in different phases of disease development. Our analysis disclosed the importance of *ANN1* in the *ex planta* life cycle of *V. dahliae*, such as the promotion of microsclerotia formation, the control of antibacterial activities, and the resistance towards osmotic stress.

## Results

### *V. dahliae* reacts to nutrient poor environments by increasing the transcript level of *ANN3* in the *ANN* secondary metabolite gene cluster

25 secondary metabolite gene clusters were identified by *in silico* prediction to be present in the *V. dahliae* JR2 genome (19). Transcriptional profiles of *V. dahliae* in nutrient-poor and nutrient-rich growth conditions were compared. RNA-seq experiments were carried out on RNA samples harvested from cultures grown only in the minimal medium CDM or in cultures initially grown in CDM and subsequently shifted to pectin-rich simulated xylem medium (SXM). Transcriptome analyses revealed that expression of the core biosynthetic enzyme-encoding genes in eight BGCs were decreased in pectin-rich SXM in comparison to the nutrient-poor CDM, whereas the biosynthetic gene in one BGC was more expressed in SXM than in CDM (Table 1).

**Table 1.**
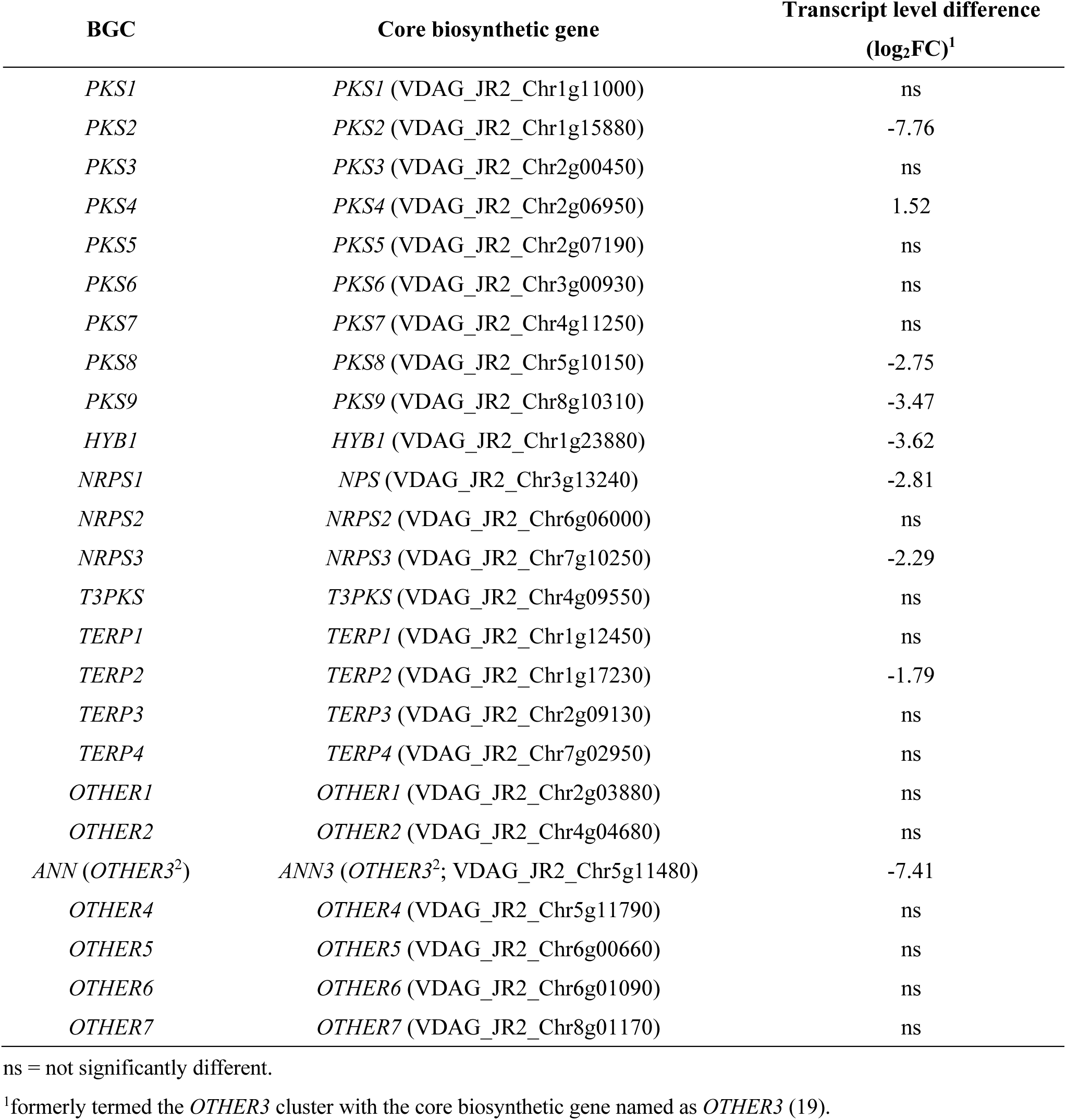
Differences in transcript level of each core biosynthetic gene of *V. dahliae* BGCs in samples shifted to SXM after initial CDM cultivation compared to samples cultured only in CDM.

*ANN3* in the *ANN* cluster (formerly termed *OTHER3* and the *OTHER3* cluster (19)) along with the *PKS2* (also known as *PKS1* in publications (11, 18, 20)) in the melanin biosynthesis cluster were two of the most differentially expressed core biosynthetic genes in nutrient-poor environments. The *ANN* cluster is located on the 5^th^ chromosome of the *V. dahliae* JR2 genome, and consists of 14 ORFs, including two genes for transcription factors as well as three biosynthetic and four transport-related genes, respectively. The function of the five remaining encoded proteins could not yet be predicted (Fig. 1a). *ANN3* encodes the core biosynthetic enzyme, which is predicted to contain an AMP-dependent synthetase/ligase domain, a phosphopantetheine binding ACP domain, and five transmembrane domains (S1 Fig.). Transcript levels of *ANN3* were analysed by qRT-PCR to compare the gene expression in different nutrient environments. The results confirmed that *ANN3* is highly induced in nutrient-poor growth condition compared to nutrient-rich media (Fig. 1b).

**Figure 1.**
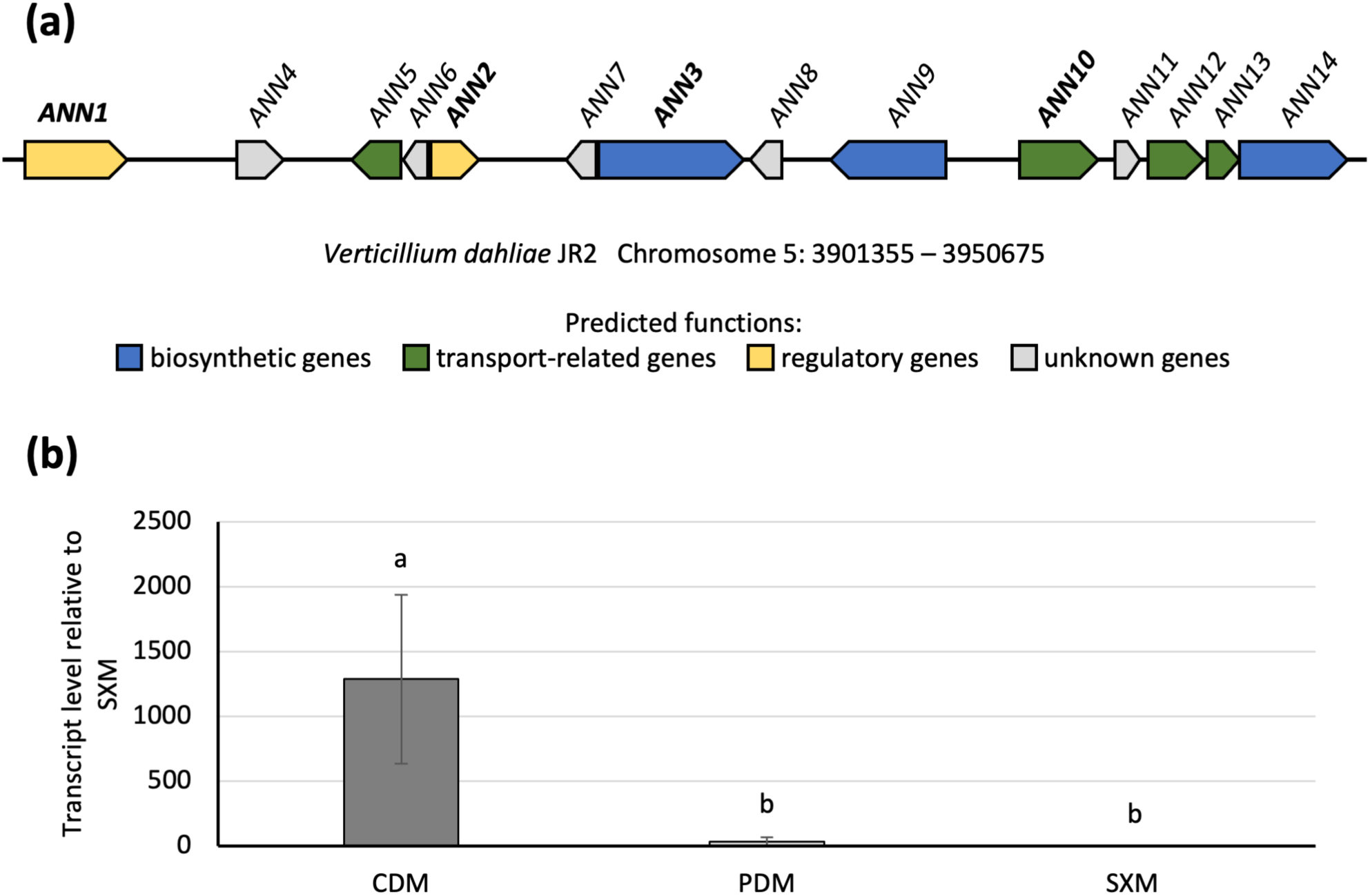
The *V. dahliae ANN* secondary metabolite cluster differentially expresses its core biosynthetic gene *ANN3* in different nutrient environments. (a) The *ANN* cluster contains two predicted regulatory genes (yellow), three predicted biosynthetic genes (blue), four predicted transport-related genes (green), and five genes with no predicted functions yet. (b) Transcript level of the core biosynthetic gene *ANN3* is significantly higher when cultured in nutrient-poor CDM compared to nutrient-rich PDM or pectin-rich SXM. qRT-PCR were performed with three biological replicates. One-way ANOVA with post-hoc Tukey HSD test was performed to compare transcript levels of *ANN3* in different culture media. A difference of the lower-case letter on top of each bar indicates significant difference (P < 0.05).

### Transcription factor Ann1 represses expression of the biosynthetic gene *ANN3* and is less stable in nutrient-poor culturing conditions

Fungal secondary metabolism is usually tightly regulated on several levels, allowing the fungus to react to specific environmental cues. Such regulation can be executed by global regulators that govern several cellular processes, or by cluster-specific transcription factors that are usually pathway-specific (12). Transcription factor-encoding genes within the *ANN* cluster, *ANN1* (VDAG_JR2_Chr5g11420a) and *ANN2* (VDAG_JR2_Chr5g11460a) were studied in detail to elucidate the regulatory mechanism of the nutrient-dependent expression of *ANN3* (VDAG_JR2_Chr5g11480a). *ANN1* is the first gene in the *ANN* cluster and encodes an ORF of 4751 base pairs (bp), which consists of three exons and two introns. The predicted Ann1 protein is 1172 amino acids (aa) in length and contains a C2H2 type zinc finger domain (PF00096) as well as a fungal specific transcription factor domain (PF04082; Fig. 2a).

**Figure 2.**
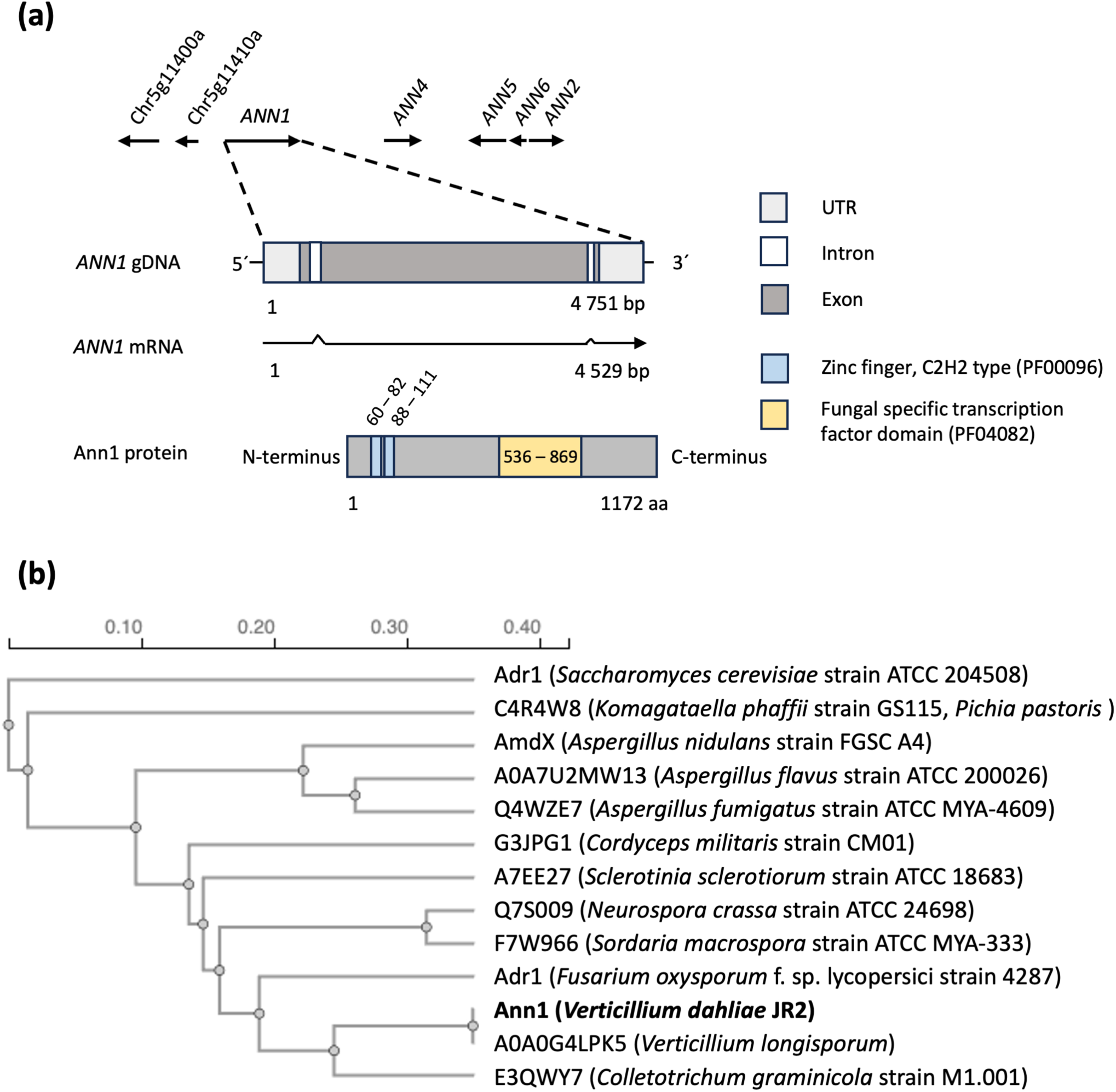
The genomic structure of the transcription factor-encoding gene *ANN1* and its orthologs in other fungal species. (a) The 4751 bp regulatory gene *ANN1* contains 2 introns (white), 2 exons (dark grey), and predicted 5’- and 3’ UTRs (light grey). The 1172 aa Ann1 protein contains a C2H2-type zinc finger domain (PF00096; 60 – 111 aa; light blue) and a fungal specific transcription factor domain (PF04082; 536 – 869 aa; yellow). (b) Phylogenic tree generated from the protein sequences of Ann1 and its orthologs in other fungal species. The names of each ortholog or the UniProt identifier are listed when a name is not given. The species and strain from which each protein originates are listed in brackets. Horizontal distance of each branch represents the number of substitutions per site.

A BLAST search for proteins in other species revealed that Ann1 orthologs are widespread among the phylum Ascomycota, with orthologs from other *Verticillium* species sharing sequence similarities above 90%, and 167 orthologs from various species that share more than 50% similarity. Among the species that contain an Ann1 ortholog, we discovered other pathogenic and non-pathogenic filamentous fungi or yeasts with wide-spread applications. Such as crop pathogen *Colletotrichum graminicola*, caterpillar fungus *Cordyceps militaris*, human pathogen *Aspergillus fumigatus*, methylotrophic yeast *Komagataella phaffii* (formerly known as *Pichia pastoris*), and baker’s yeast *Saccharomyces cerevisiae* (Fig. 2b). AmdX from *Aspergillus nidulans* was previously studied to be involved in the nitrogen utilisation regulatory network (25), and Adr1 from *S. cerevisiae* controls the utilisation of non-fermentable carbon sources (26, 27).

The deletion, the Gfp-fused complementation, as well as the over expression strains of *ANN1* or *ANN2* were generated and confirmed by Southern hybridisation to examine how the *ANN* cluster is being governed by transcription factors Ann1 and Ann2. All strains were cultured in PDM, SXM or CDM for 5 days before mycelia were harvested for RNA extraction. qRT-PCR experiments were performed to examine if Ann1 or Ann2 regulate the core biosynthetic enzyme-encoding gene *ANN3*, and whether Ann1 and Ann2 mutually influence each other’s expression. Our results verified the transcript levels of the mutant strains and revealed that the absence of *ANN1* resulted in a two-fold increase in *ANN3* transcript level when cultured in nutrient-rich PDM and SXM, but it doesn’t further increase in CDM. In contrast, ectopically over expressing *ANN1* seven-fold resulted in a five-fold decrease in *ANN3* transcripts when cultured in minimal medium CDM, but the expression of *ANN3* doesn’t further reduce in PDM and SXM (Fig. 3a and 3b). Ann1 therefore acts as a transcriptional repressor of the core biosynthetic enzyme-encoding gene *ANN3*. Nutrient availability in each tested media affect *ANN3* expression levels more than the transcript levels of transcription factor *ANN1*. We also discovered that deletion of *ANN1* resulted in a decrease in expression levels of the second transcription factor-encoding gene *ANN2* in all culture media (Fig. 3c). On the contrary, neither *ANN1* nor *ANN3* expression levels changed more than two-fold upon the deletion or over expression of *ANN2* (S2 Fig.). Our results suggest that Ann1 acts upstream of Ann2 as a transcriptional activator, and expression of *ANN3* is mainly controlled by the transcriptional repressor Ann1 in the tested conditions and not by the functionally elusive Ann2.

**Figure 3.**
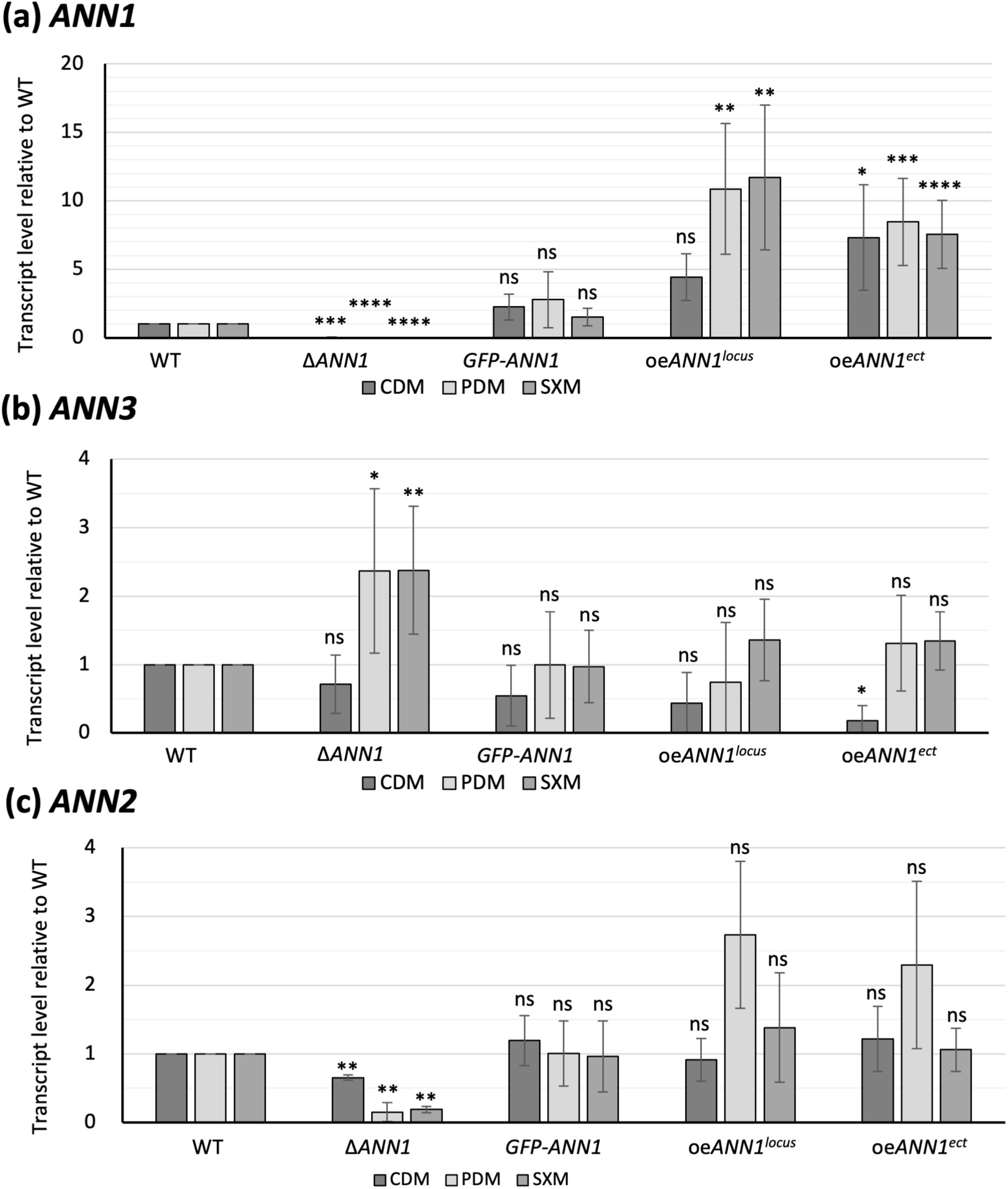
*V. dahliae* Ann1 is a transcriptional repressor of the core biosynthetic enzyme-encoding gene *ANN3*, and an activator of transcription factor-encoding *ANN2*. Transcript levels of (a) *ANN1*, (b) *ANN3*, and (c) *ANN2* in the *ANN1* mutant strains were analysed by qRT-PCR. *ANN1* transcript levels were verified to be elevated in both in locus (locus) and ectopic (ect.) over expression strains. Compared to the WT gene expression levels, the transcript of *ANN3* is increased more than 2-fold in PDM and SXM when *ANN1* is deleted. Transcript level of *ANN3* is reduced more than 5-fold in the ectopic *ANN1* over expression strains when cultured in CDM. Expression of the transcription factor-encoding gene *ANN2* is reduced in all tested media when *ANN1* is deleted. qRT-PCR were performed with at least three biological replicates. One-sample t-test was performed to compare transcript levels of each tested strain to the WT expression levels under the same culturing condition (*, P < 0.05; **, P < 0.01; ***, P < 0.001; ****, P ≤ 0.0001; ns, not significantly different).

Western experiments were performed to study how Ann1 is involved in the nutrient-dependent regulation of *ANN3*. Presence and stability of Ann1 protein in different nutrient environments were examined. The complementation strain that expresses Gfp-fused Ann1 under the control of its native promoter and the WT strain were cultured in either CDM, PDM, or SXM for five days before mycelia were harvested for protein extraction. The full-length Gfp-Ann1 protein is only detectable when the culture was grown in the nutrient-rich PDM. In contrast, full-length Gfp-Ann1 is undetectable in the minimal medium CDM, and the free-Gfp degradation product was very prominent (Fig. 4a). Intracellular localisation of Ann1 was investigated to verify the presence of Ann1 as a transcriptional regulator in the nucleus. The Gfp-Ann1 complementation strain was transformed to ectopically express Rfp-fused histone H2b for nuclear visualisation. The strain was cultured in PDM and observed under the fluorescence microscope. Since the Gfp-Ann1 signal co-localises with the histone-Rfp signal, we confirmed the nuclear localisation of Gfp-Ann1 (Fig. 4b). Our results indicate that Ann1 is most stable in nutrient-rich environments, and it is present in the nucleus. This aligns with the Ann1 function as a transcription factor, and the elevated *ANN3* transcript levels in nutrient-poor environments, in which the repressor Ann1 is mostly degraded.

**Figure 4.**
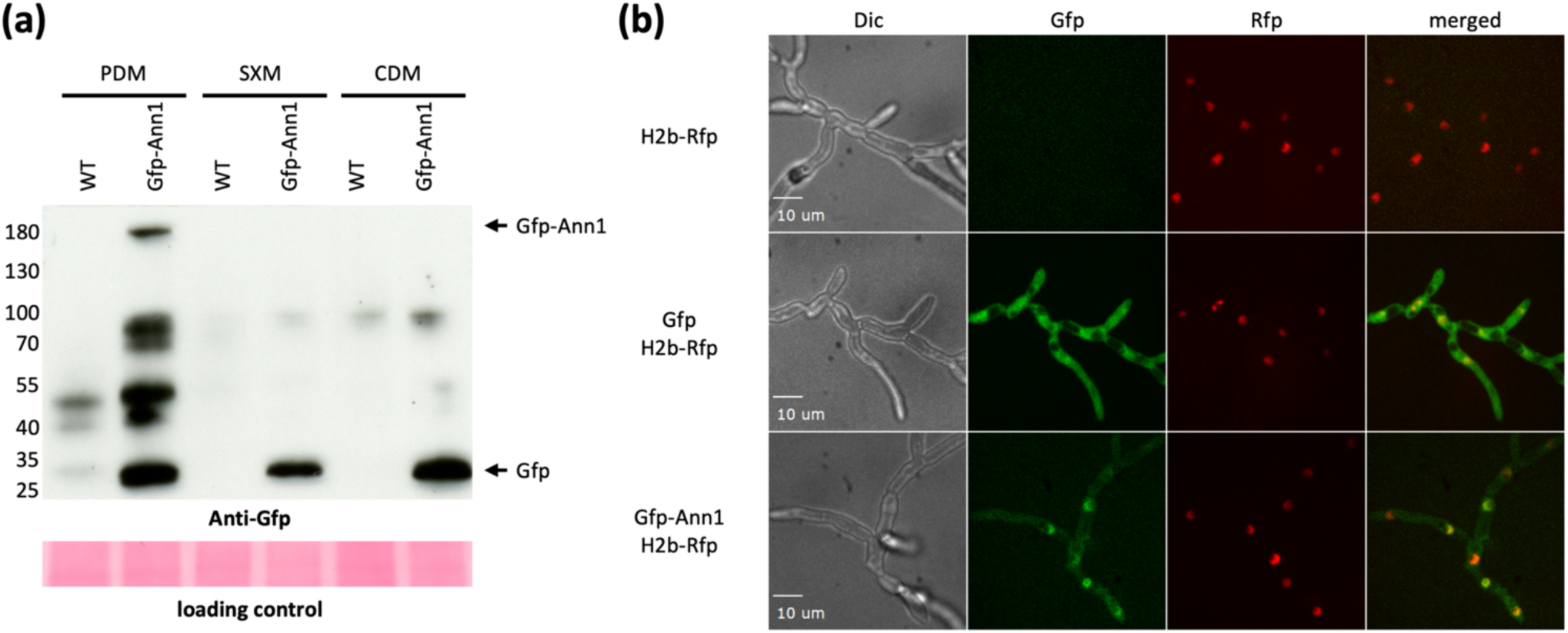
Functional Gfp-Ann1 fusion protein is less stable when *V. dahliae* is cultured in nutrient-poor conditions and it is mainly localized in the nucleus in nutrient-rich PDM. (a) The full-length Gfp-Ann1 is only present when *V. dahliae* is cultured in nutrient-rich PDM. Gfp-Ann1 is mostly degraded when *V. dahliae* is cultured in minimal medium CDM and pectin-rich medium SXM. Western experiments were performed with antibody recognising Gfp, and the loading control was done by Ponceau staining or TGX Stain-Free Protein Gels (Bio-Rad) to ensure that an equal amount of protein was loaded in each lane. The expected size of free Gfp is 28 kDa, and Gfp-Ann1 is 155 kDa, and the expected positions are labelled on the right of the image. The expected protein sizes at each position were marked on the left according to the position of the protein ladder. (b) Florescence microscopy images were taken for the WT strain, the free Gfp strain, and the strain expressing the Gfp-Ann1 fusion protein. All strains were cultured in PDM and ectopically express the histone H2b-Rfp fusion protein for nuclear visualisation. This figure presents differential interference contrast view (Dic), green fluorescent protein filter view (Gfp), red fluorescent protein filter view (Rfp), and a merged view of the Gfp and Rfp channels.

### The *ANN* secondary metabolite gene cluster is regulated by the Som1-Vta3-Vta2-Mtf1 network

Som1, Vta3, and Vta2 are transcription factors that regulate sequential steps in the early plant root infection processes of *V. dahliae*, including adhesion, root surface propagation, and colonisation of roots. The transcription factor Som1 promotes the expression of *VTA3* and *VTA2*, while *VTA2* expression is also governed by Vta3 (6, 7). In contrast, transcription factor Mtf1 controls the expression of genes involved in the later phase of the infection, and its expression is repressed by Vta3 and Vta2 (23). In addition to the transcription factors locally present within the cluster, we further investigated whether the *ANN* cluster is regulated by these global regulatory proteins. qRT-PCR experiments were performed to compare the gene expression levels of *ANN1* and *ANN3* in the WT or in the *SOM1*, *VTA3*, *VTA2*, or *MTF1* deletion strains. The tested strains were inoculated in liquid CDM, and mycelia were harvested 5 days post inoculation (dpi) for RNA extraction. The absence of *SOM1* or *VTA2* led to a 19-fold and three-fold decrease in *ANN3* transcripts respectively (Fig. 5). The changes in *ANN3* transcripts were less than two-fold in the absence of *VTA3*. In contrast, deletion of *MTF1* resulted in a 6-fold increase in *ANN3* transcript levels, indicating that the expression of *ANN3* is directly or indirectly repressed by Mtf1. By contrast, the *ANN1* transcript level remained within a two-fold range of variation in all the tested deletion strains. These results suggest that *ANN3* can be independently regulated by multiple transcriptional regulators, and its expression is favoured during the *ex planta* to *in planta* transition but suppressed in later phases of disease development.

**Figure 5.**
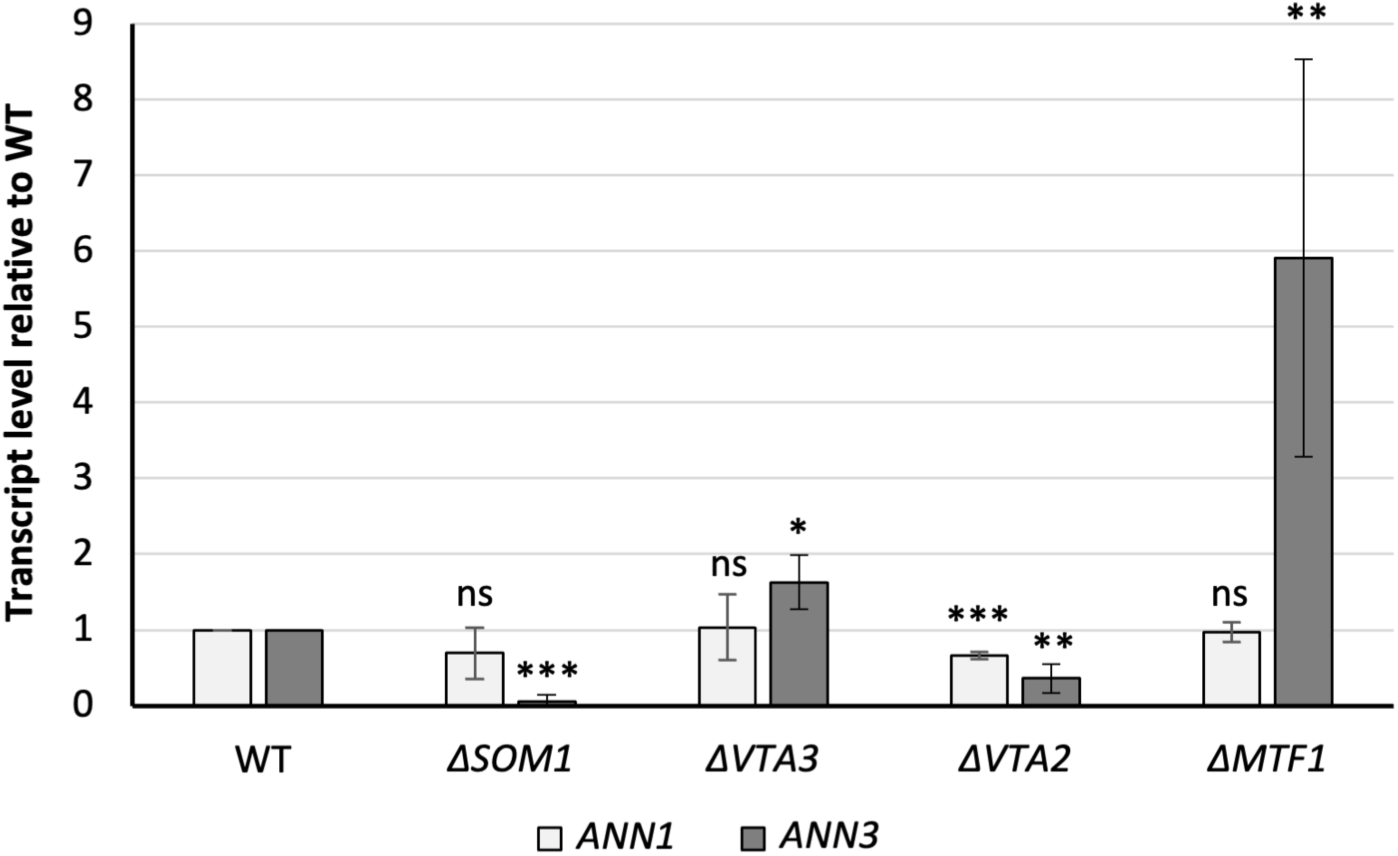
The *V. dahliae* Som1-Vta-Mtf1 regulatory network is directly or indirectly controlling the expression levels of *ANN1* and *ANN3*. Expression of *ANN3* is promoted by Som1 and Vta2 but repressed by Mtf1, and *ANN1* transcript level is decreased in the Δ*VTA2* strain. Transcript levels of *ANN1* and *ANN3* in the deletion mutants of potential upstream regulators were analysed by qRT-PCR with at least four biological replicates performed. All tested strains were cultured in liquid CDM for 5 days before the mycelia were harvested for RNA extraction. One-sample t-test was performed to compare transcript levels of each tested strain to the WT expression levels under the same culturing condition (*, P < 0.05; **, P < 0.01; ***, P < 0.001; ns, not significantly different).

### Ann1 promotes melanisation and osmotic stress resistance

*V. dahliae* forms microsclerotia, which are heavily melanised resting structures that allow survival in unfavourable growth conditions (1). The active phase of the *ex planta* life cycle of *V. dahliae* is short and transitory. However, it involves survival in competitive and potentially nutrient-limiting conditions, before the fungus reaches the plant host (1). We examined the *ex planta* growth phenotype by point inoculating an equal number of conidiospores of each *ANN1* or *ANN2* mutant strains and the WT strain on agar plates with different media. Colonies of the Δ*ANN1* strain on every tested medium were smaller in diameter, implying that Ann1 is required for regular vegetative growth (Fig. 6a and 6b). The diameters of Δ*ANN1* colonies cultured on CDM plates were 27% smaller than WT colonies. In comparison, Δ*ANN1* colonies cultured on CDM plates with cell wall stressors Congo red or sorbitol were 35% smaller, while the colonies on CDM plates with osmotic stressor NaCl were 70% smaller (Fig. 6a and 6b). This suggests that *ANN1* promotes osmotic stress resistance but not cell wall-stress resistance. Melanisation levels of the colonies grown on pectin-rich SXM and cellulose-containing CDM plates were quantified. The Δ*ANN1* colonies hardly melanised in either medium, while the *ANN1* over expression colonies showed an increased level of melanisation (Fig. 6a and 6c). Similar growth assays were also performed with the *ANN2* mutant strains. All colonies of the *ANN2* mutant strains were indistinguishable from WT colonies in size (S3 Fig.). The *ANN2* over expression strain was less melanised at 3 dpi when cultured on SXM plates. However, no differences in melanisation can be observed between the different *ANN2* mutant strains and the WT strain at 10 dpi (S3 Fig.). These results imply that Ann1 is required for vegetative growth and facilitates resistance to unfavourable growth conditions by resisting osmotic stress and promoting microsclerotia formation.

**Figure 6.**
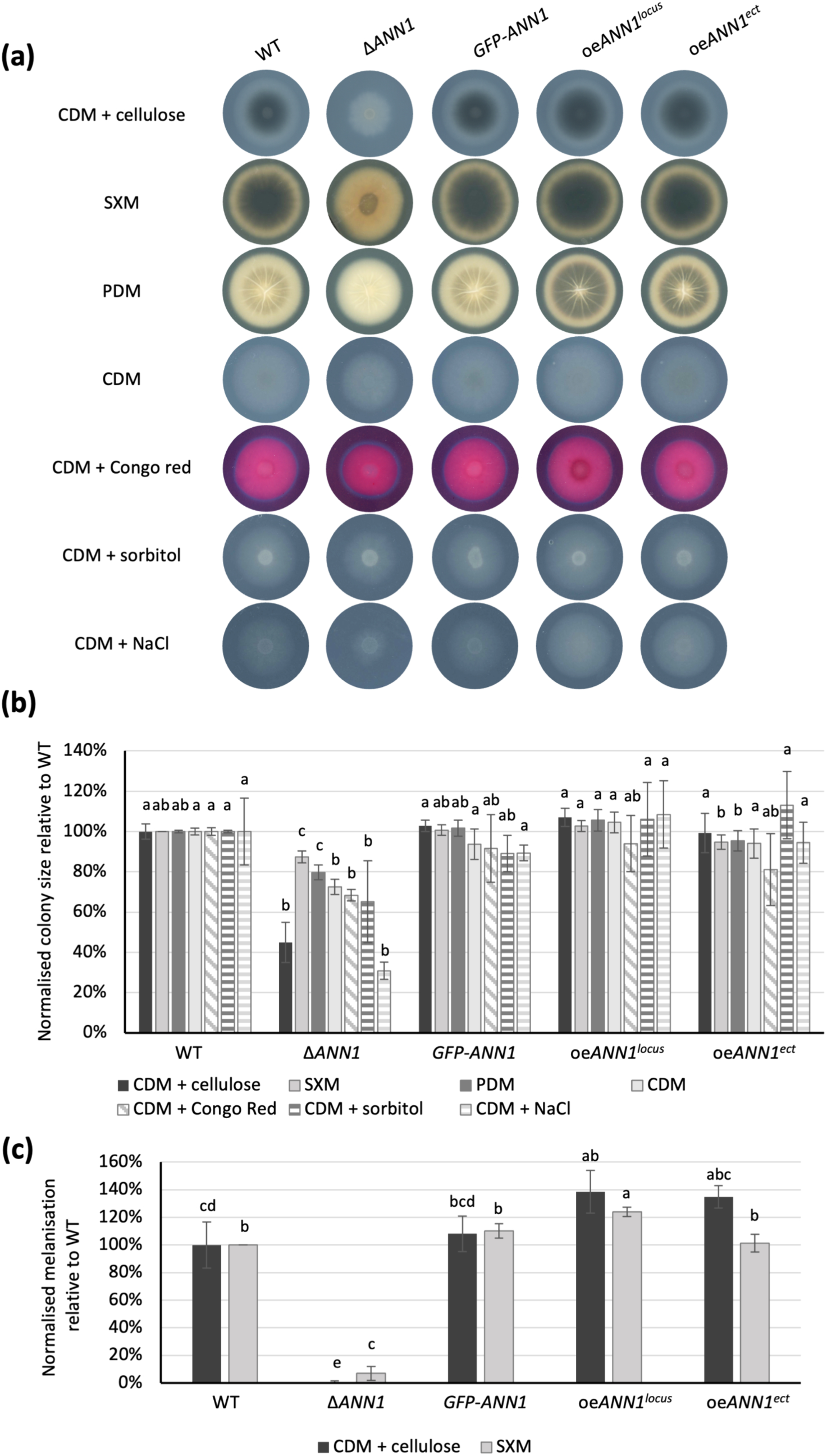
Ann1 is involved in *V. dahliae* filamentous growth and microsclerotia formation. (a) The *ex planta* growth phenotype was compared between WT and the *ANN1* mutant strains at 10 dpi on minimal medium supplemented with cellulose (CDM + cellulose), pectin-rich medium (SXM), potato dextrose medium (PDM), minimal medium (CDM), minimal medium supplemented with Congo red (CDM + Congo red), minimal medium supplemented with sorbitol (CDM + sorbitol), and minimal medium supplemented with NaCl (CDM + NaCl). 50,000 spores were point-inoculated on each agar plate and the plates were scanned 10 dpi. (b) Colony sizes were measured and normalised to the size of the WT colony in the corresponding culture medium. The Δ*ANN1* strain had smaller colony size when cultured on CDM + NaCl, which suggests that it is less resistant to osmotic stress conditions. (c) Colony sizes and melanisation were measured by Fiji ImageJ and normalised by the colony size or degree of melanisation of WT in the respective culture medium. Melanisation is increased in *ANN1* over expression strains and reduced in the Δ*ANN1* strain. One-way ANOVA with post-hoc Tukey HSD test was performed to compare the colony size and the degree of melanisation of each strain on the same culture medium. A difference of the lower-case letter on top of each bar indicates significant difference (P < 0.05).

### Ann1 regulates antibacterial activity of *V. dahliae*

The *ex planta* life cycle of *V. dahliae* consists of a potentially long period of dormancy in the form of microsclerotia, and a relatively short period of time when it germinates upon encountering suitable growth conditions (1). Complex interactions in the soil between *Verticillium* species, its plant host, and soil bacteria shapes this short *ex planta* phase in the form of competition and communications (1, 5). *V. dahliae* is well known to evade competition by growing away (8). Here we investigated whether the fungus is also able to chemically defend itself from bacterial competitors. As Ann1 is shown to be involved in the transition from the *in planta* to *ex planta* life cycle by promoting microsclerotia formation, and its protein stability is compromised in nutrient-poor growth conditions, we further studied whether Ann1 is also involved in other phases of the *ex planta* life cycle. Bacterial-fungal interaction assays were performed by co-culturing the *ANN1* mutant strains as well as the WT strain with the Gram-positive *B. subtilis* 168 or the Gram-negative *E. coli* DH5⍺. The interaction interphase between *B. subtilis* 168 and the *V. dahliae* colonies showed that they grow close or touch each other when either the *V. dahliae* WT or the *GFP-ANN1* complementation strain was used (Fig. 7a). However, a sharp edge of the *B. subtilis* 168 colony that is positioned apart from the *V. dahliae* colony can be observed when the Δ*ANN1* strain was grown on the same plate (Fig. 7a), indicating the presence of an antibacterial compound secreted by the Δ*ANN1* strain. On the contrary, the *B. subtilis* 168 and the *V. dahliae* colonies touch or even grow into each other when the *ANN1* over expression strains were inoculated (Fig. 7a). The distances between the bacterial and the fungal colonies were determined as a measurement of the antibacterial activity, and the absence of *ANN1* significantly increases the ability of *V. dahliae* to cope with antagonistic interactions against *B. subtilis* 168 (Fig. 7b). Statistical analyses were also performed on the results from the *E. coli* DH5⍺ interaction assays. However, no statistical significance can be observed possibly due to the limited growth of *E. coli* DH5⍺ in the unfavourable culturing conditions and a large variation between experimental results (S4 Fig.). Since previous results indicated that the Gfp-Ann1 protein is less stable in nutrient-poor conditions, we also examined if Gfp-Ann1 protein stability is affected by the co-cultivation of *B. subtilis* 168 and *V. dahliae*. Compared to the *V. dahliae GFP-ANN1* pure culture, signal intensity of the full-length Gfp-Ann1 has reduced whereas more free-Gfp degradation product could be detected in the co-cultivated samples (Fig. 7c). This indicates that Ann1 is less stable upon encountering antagonists, and the complete deletion of *ANN1* may have resulted in an increased presence of a secreted antibacterial compound.

**Figure 7.**
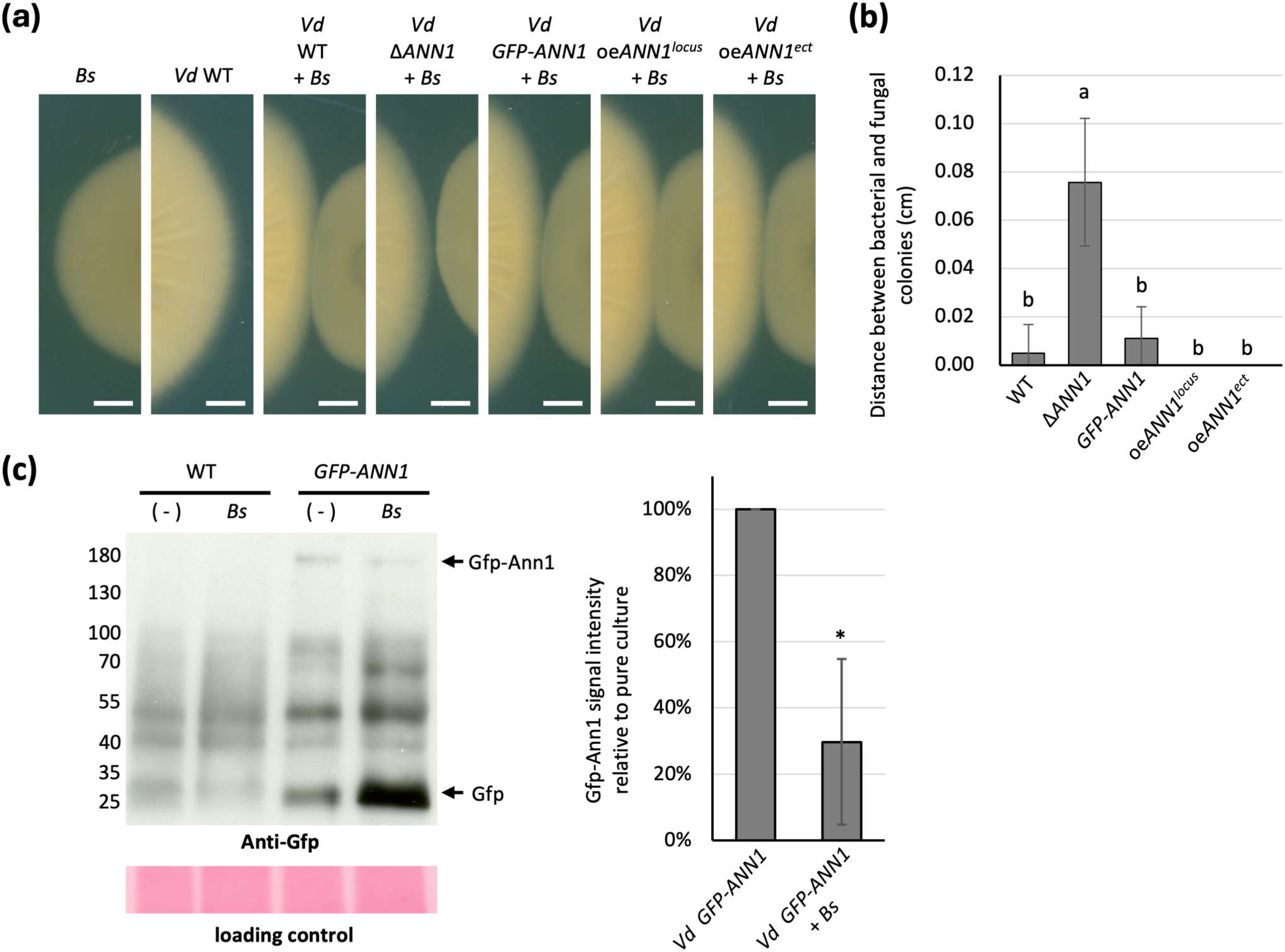
Ann1 negatively regulates antibacterial activity of *V. dahliae*. (a) Distance between the *V. dahliae* and *B. subtilis* 168 colonies was increased when *ANN1* is deleted, whereas the two co-cultured colonies grow into each other when *ANN1* is over expressed. The WT *V. dahliae* and the *ANN1* mutant strains were spotted on 100 mL PDM plates, and 10 µL of bacterial suspension at OD_600_ = 0.01 was spotted 2.5 cm apart from the centre of the *V. dahliae* colony at 3 dpi. The *V. dahliae* and bacterial colonies were co-cultured for an additional 7 days before the results were evaluated (white bar = 1 cm). (b) Antibacterial activity of *V. dahliae* against *B. subtilis* 168 was quantified by the distance between the two colonies. One-way ANOVA with post-hoc Tukey HSD test was performed to compare the antibacterial activity of each strain towards *B. subtilis* 168. A difference of the lower-case letter on top of each bar indicates significant difference (P < 0.05). (c) The abundance of intact Gfp-Ann1 is reduced when *V. dahliae* is cultured with *B. subtilis* 168. The Gfp-Ann1 and WT strains were cultured in liquid PDM for 5 days, *B. subtilis* 168 were co-cultured with *V. dahliae* for 24 hr in designated samples. Western experiments were performed with antibody recognising GFP, and Ponceau staining or TGX Stain-Free Protein Gels (Bio-Rad) served as loading control to ensure that an equal amount of protein was loaded in each lane. The expected size of free Gfp is 28 kDa, and Gfp-Ann1 is 155 kDa, and the expected positions are labelled on the right of the image. The expected protein sizes at each position were marked on the left according to the position of the protein ladder. The intensity of the Gfp-Ann1 signals were quantified and normalised by *V. dahliae GFP-ANN1* pure culture signals. One-sample t-test was performed to compare signal intensity of the co-cultivated samples to the pure culture samples (*, P < 0.05).

### Ann1 inhibits conidiation but is dispensable for pathogenicity

Upon entry into the plant vasculature system, *V. dahliae* spreads rapidly by forming asexual conidiospores that cause systemic infection (1). Conidiation is a process that takes place during the *in planta* life cycle of *V. dahliae*. Equal numbers of conidiospores of WT and each *ANN1* mutant strains were inoculated in liquid SXM and the conidiospores were harvested and counted 5 dpi to study if *ANN1* is involved in the process of conidiation. The Δ*ANN1* and the *GFP-ANN1* strains showed no significant difference in their ability to form conidiospores compared to the WT strain, whereas the *ANN1* over expression strains showed a conidiospore reduction of approximately 50% compared to WT counts (Fig. 8a). Although the *ANN1* mutant strains differ in their conidiation ability, tomato plant infection experiments revealed no significant differences in the disease scores between seedlings inoculated with different *ANN1* mutant strains at 21 dpi (Fig.8b). Therefore, *ANN1* is likely dispensable for pathogenicity.

**Figure 8.**
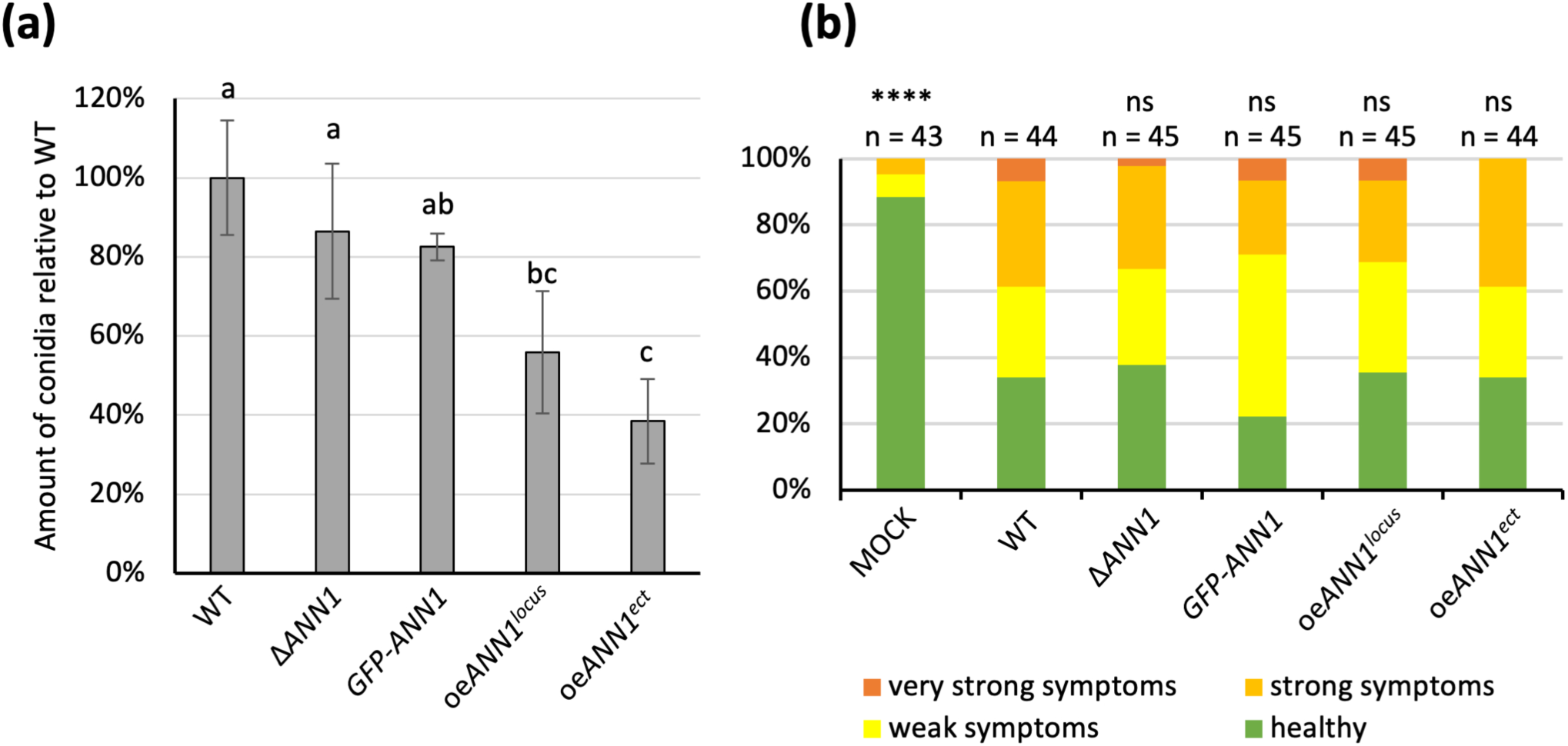
Ann1 inhibits conidiation but is not involved in virulence towards tomato plants. (a) The *ANN1* over expression strains produce less conidia than WT and other *ANN1* mutant strains. The *ANN1* mutant strains and the WT strain were cultured in SXM for 5 days before the conidiospores were harvested and counted. Four biological replicates with four technical replicates each were performed, and all results of the same biological replicate were normalised by the WT conidiation level. One-way ANOVA with post-hoc Tukey HSD test was performed to compare conidiation of each strain. A difference of the lower-case letter on top of each bar indicates significant difference (P < 0.05). (b) WT and the *ANN1* mutant strains were tested for *in planta* phenotype by infecting 10-days old tomato seedlings, and the disease scores were assigned 21 dpi. The disease scores of seedlings inoculated by water (MOCK) and each *ANN1* mutant strain were compared to the disease scores of the WT-inoculated seedlings by two-tailed Mann-Whitney U test (****, P < 0.0001; ns, no significant difference).

### Four identified metabolites represent potential *ANN* cluster products

We aimed to identify the products of the *ANN* cluster, because manipulation of transcript levels of the *ANN3*-controlling *ANN1* led to an altered antibacterial activity. Metabolites from the *ANN1* and the *SOM1* mutant strains were analysed as Ann1 represses, but Som1 activates the transcription of *ANN3*. CDM agar plates were inoculated with WT or the *ANN1*/*SOM1* mutant strains and incubated for 14 days before metabolites were extracted for LC-MS analysis. Empty CDM agar plates served as control. By analysing the total ion chromatogram of the LC-MS results, 11 metabolites were discovered to have altered abundance in either the Δ*ANN1* strain or the *ANN1* over expression strains. The Δ*ANN1* strain had an increased abundance of substances (I), (II), (IX), and (X) compared to the WT or over expression strains. (Fig. 9). The production of 15 metabolites was affected by the deletion or the over expression of *SOM1* (S5 Fig.). The deletion of *SOM1* reduced the production of 13 metabolites. Within which, substances (I) – (IV) and (VI) – (X) also had altered abundance in the *ANN1* mutant strains, and the production of substances (XII) – (XIV) is known to be controlled by the velvet proteins (28). The velvet proteins are also global transcriptional regulators, and the expression of *VEL1* is known to be governed by Som1 (6, 28). Since the expression of the core biosynthetic gene *ANN3* is repressed by Ann1 and promoted by Som1, possible candidates for the metabolite produced by the *ANN* cluster or the derivatives of the product can be limited to compounds (I), (II), (IX), and (X).

**Figure 9.**
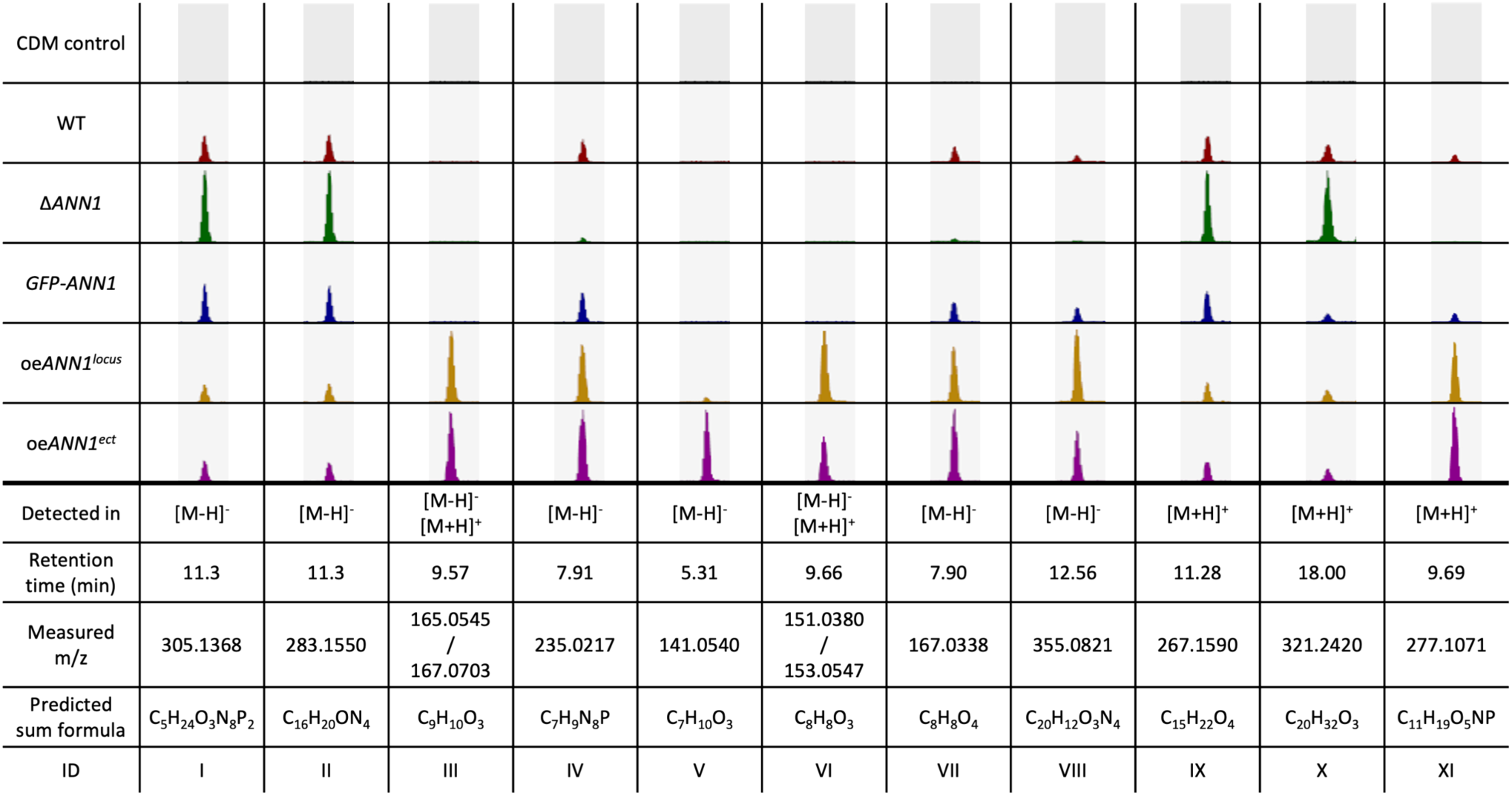
Eleven metabolites had altered abundance in the *V. dahliae ANN1* mutant strains. Four of the listed metabolites are more abundant in the Δ*ANN1* strain (I), (II), (IX), (X), and seven are more abundant in the over expression *ANN1* strains (III) – (VIII), (XI). All tested strains were grown on CDM plates for two weeks before the metabolites were extracted by 50% ethyl acetate from the homogenised agar. Extracted ion chromatograms of masses that had differed abundance in the tested strains are shown, and 5 ppm of mass deviation was tolerated. The height of each peak corresponds to the relative abundance of a certain mass within the tested strains. The predicted sum formula of each compound is calculated by the calculated exact mass.

Metabolites were also extracted from the interaction interface between the *B. subtilis* 168 colony and the *V. dahliae* WT or *ANN1* mutant strain colony on PDM plates (Fig. 7a) to analyse if the metabolite produced by the *ANN* cluster or the derivatives are possibly contributing to the antibacterial activity. Out of the 18 fungal metabolites with altered abundance in the *ANN* mutant strains detected in the co-culture interface on PDM, substances (I), (II), (IV), (VII), (IX), and (X) overlapped with previously detected metabolite samples from pure cultures of *ANN1* mutant strains on CDM (S6 Fig.). Combined with the metabolites discovered from pure cultures of *ANN1* or *SOM1* mutant strain, compounds (I), (II), (IX), and (X) are the most probable candidates as the product or its derivatives of the *ANN* cluster, and it is possible that they are involved in antibacterial activity.

## Discussion

Phytopathogenic Verticillia perceive different growth conditions during their life cycle and react by adjusting their transcriptome or secretome (8, 24, 29). By comparing the *V. dahliae* transcriptome in nutrient-rich and nutrient-poor environments, we revealed that eight biosynthetic gene clusters (BGCs) in the *V. dahliae* genome had their core biosynthetic enzyme-encoding genes differentially expressed. Among which, *ANN3* of the *ANN* cluster and *PKS2* (also published as *PKS1* (11, 19)) in the melanin biosynthesis cluster were the most highly transcribed core genes in nutrient-poor environments. Melanin ensures the dormancy phase of the life cycle by protecting microsclerotia (18, 20), whereas the biological role of the *ANN* cluster was so far unstudied. The regulation of fungal secondary metabolism often involves complex signalling pathways. Specific environmental cues activate certain pathways, and transcription factors act in the end to activate or repress genes involved in secondary metabolite biosynthesis (12). In this study, we discovered that the expression of *ANN3* is governed by the in-cluster transcriptional repressor Ann1 (Fig. 3), as well as global transcriptional regulators Som1, Vta2, and Mtf1 (Fig. 5). It is estimated that 60% of fungal biosynthetic gene clusters encode in-cluster transcription factors that act as pathway-specific regulators (12). Although most of these clusters only have one pathway-specific transcription factor, in rare cases, they can also include more regulatory genes. For example, both *VTA1* and *CMR1* are located within the melanin biosynthesis cluster in the *V. dahliae* genome (18, 20), and *AflR* and *AflS* in the aflatoxin biosynthesis cluster of *A. flavus* (30). In the case of *VTA1* and *CMR1*, the presence of two transcription factors enables the activation of melanin biosynthesis by two independent pathways, which can be triggered by different environmental cues. An additional in-cluster transcription factor also gives regulatory flexibility, as each member gene within the melanin biosynthesis cluster can also be controlled differently by *VTA1* and *CMR1* (18, 20). The transcription factor Ann1 mainly controls the nutrient-based expression of *ANN3* (Fig. 3 & 4), but the regulatory role of Ann2 towards *ANN3* remains yet unknown in our tested conditions (S2 Fig.). The presence of *ANN2* suggests a potential of the *ANN* cluster to be activated or repressed by environmental stimuli other than nutrient availability, or the possibility of a different modification or transportation of the produced metabolite.

Our results revealed that Ann1 negatively regulates the core biosynthetic enzyme-encoding *ANN3*, and Ann1 is more stable in nutrient-rich environment and mostly degraded in nutrient-poor culturing conditions, respectively (Fig. 3b & 4a). To date, two orthologs of Ann1 have reported functions (Fig. 2). Adr1 in *S. cerevisiae* regulates the utilisation of non-glucose carbon sources (26, 27), whereas *A. nidulans* AmdX allows the fungus to utilise simple amides as the sole nitrogen source by activating the expression of acetamidase-encoding AmdS (25). Genes regulated by *Sc*Adr1 and *An*AmdX are repressed when the preferred nutrient source is available (26, 31). In the case of *Sc*Adr1, the presence of glucose results in the phosphorylation of *Sc*Adr1, which leads to the subsequent inactivation of downstream target genes (32). The similar involvement of Ann1, *Sc*Adr1, and *An*AmdX in adapting to limited or unfavourable nutrient sources suggests an evolutionarily conserved function within the Ascomycota phylum.

This study further investigated the regulatory network that governs *ANN1* and *ANN3* to understand the role of the *ANN* cluster in the *V. dahliae* life cycle. Som1 and Vta2 are known to be regulating the early root-colonisation process. In contrast, Mtf1 is involved in later phases of the infection (6, 7, 23). The fact that *ANN3* expression is promoted by Som1 and Vta2 and repressed by Mtf1 (Fig. 5) suggests that the *ANN* cluster is less needed in the later phases of disease development and is more important during the *ex planta* to *in planta* lifestyle transition. The fact that altered *ANN1* expression levels did not result in differences in disease symptoms further suggests that the *ANN* cluster is dispensable for later disease development and virulence (Fig. 8b). As *V. dahliae* is mostly dormant in resistant microsclerotia during its *ex planta* phase, and there are no effective control methods once the fungus enters the plant vasculature system (33), the short biotrophic *ex planta* period presents a window of opportunity to prevent Verticillium wilt. Earlier studies on different *Fusarium oxysporum* strains suggested that pathogenic isolates are less well-adapted to competitions in the soil compared to non-pathogenic isolates (34). Supporting this model, the phytopathogen *V. longisporum* decreases the expression of genes involved in metabolic processes and cell wall organisation upon encountering antagonists, and the hyphal tip also grows away from co-cultivated bacteria to avoid contact (8). The *ANN* cluster supports that *V. dahliae* can also act on bacterial antagonists as alternative to taking the usual “wait or escape” strategy. The *ANN* cluster acts when the fungus must cope with unfavourable growth conditions and has to counteract against bacterial competitors. Competition from antagonists can lead to nutrient limitation, thus triggering the degradation of Ann1 (Fig. 7c). The reduced Ann1 de-represses the expression of the core biosynthetic enzyme-encoding *ANN3* and possibly results in the increased production of metabolites (I), (II), (IX), or (IX) that may potentially be involved in inhibiting bacterial growth (Fig. 7 & 9). Since the expression of genes involved in metabolic processes and cell wall organisation decrease during competition (8), the reduced vegetative growth and resistance of the *V. dahliae* Δ*ANN1* strain towards osmotic stress may be a result of the growth-defence trade-off. As *V. dahliae* spends most of its biotrophic phase in the plant host, *in planta* developmental processes such as conidiation and microsclerotia formation are subjected to complex regulations (1, 6, 28, 35). Upon the initial root infection, rapid spread and growth is needed to secure a successful host colonisation. The *ANN* cluster is less involved in the *in planta* life cycle presumably because it is a mechanism for defence. Instead of supporting processes needed for plant colonisation, Ann1 is shown to withhold conidiation and promote dormancy (Fig. 6 & 8a).

In summary, our study reports the *ANN* cluster as a novel defence mechanism for the phytopathogenic *V. dahliae* to cope with antagonists and unfavourable nutrient environments. The biological functions and the regulatory relationship between members of the *ANN* cluster are summarised in Fig. 10. The in-cluster transcription factor Ann1 influences growth, developmental processes, and interspecies competition. Involvement of the *ANN* cluster in the short yet exposed *ex planta* growth phase makes it an interesting target for the future development of effective Verticillium wilt controlling methods.

**Figure 10.**
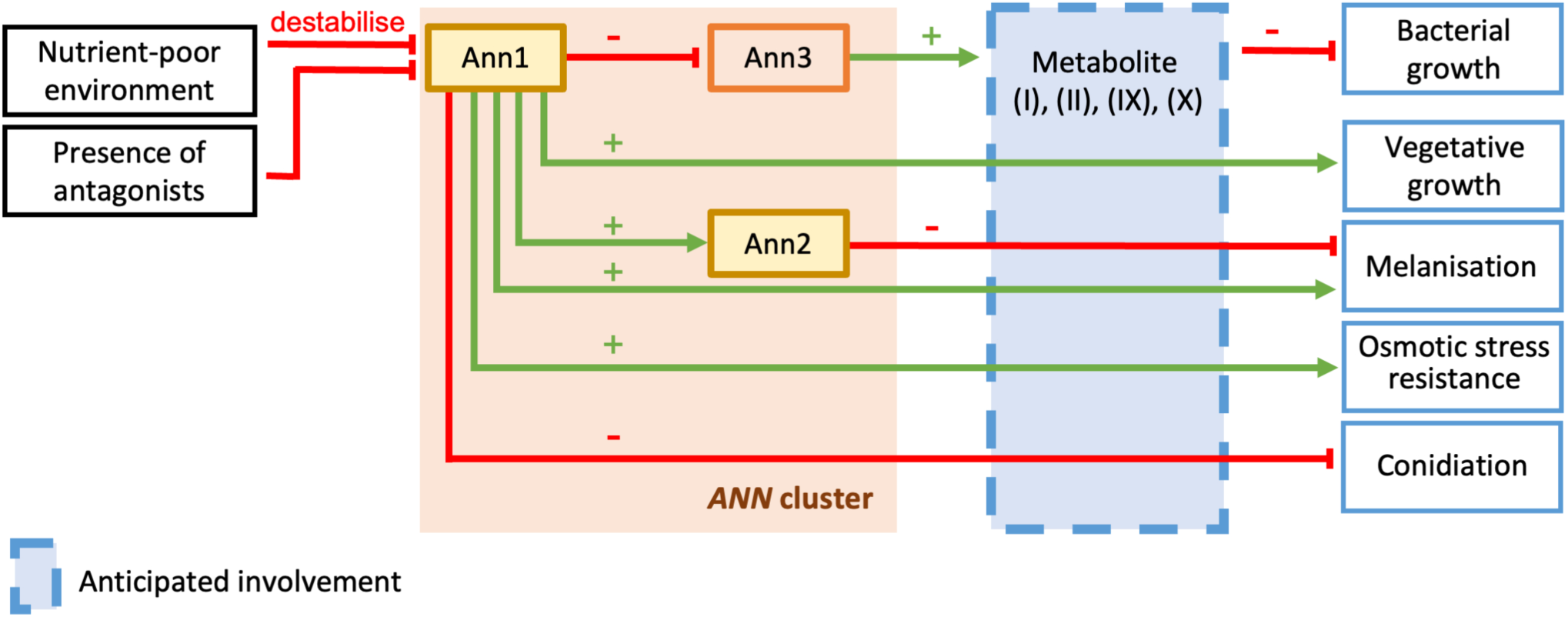
Regulatory relationships between the members of the *V. dahliae ANN* cluster and their functions in biological processes. The in-cluster transcription factor Ann1 represses the expression of the core-biosynthetic enzyme-encoding *ANN3* but activates the transcription factor-encoding *ANN2*. Nutrient-poor environments and the presence of antagonists destabilise the Ann1 protein. Ann1 withholds antibacterial activity possibly by inhibiting the production of metabolite synthesised by Ann3. Ann1 promotes melanisation, while Ann2 plays a minor role in inhibiting it. Ann1 additionally promotes vegetative growth and osmotic stress resistance but inhibits conidiation. Transcription factors are presented in yellow boxes, the biosynthetic enzyme in orange, and the anticipated involvement of metabolites is depicted in a blue box with dashed boarder.

## Materials and Methods

### Strains and growth conditions

*Verticillium dahliae*, *Escherichia coli*, *Agrobacterium tumefaciens*, and *Bacillus subtilis* strains used in this study are listed in S1 Table. Bacterial strains were cultured in lysogeny broth medium (36) and supplemented with kanamycin (AppliChem) with a final concentration of 100 µg/mL when required. *E. coli*, *A. tumefaciens*, and *B. subtilis* were incubated at 37°C, 25°C, and 30°C respectively when cultured alone. *E. coli* and *B. subtilis* were incubated at 25 °C when co-cultivated with *V. dahliae*. *V. dahliae* strains were grown at 25°C in simulated xylem medium (SXM), potato dextrose medium (PDM), or Czapek Dox medium (CDM) as previously described (37). SXM was supplemented with 300 µg/mL cefotaxime (Fujifilm Wako Chemicals), and PDM was supplemented with 50 μg/mL hygromycin B (Invivogen) or 72 μg/mL nourseothricin (Werner BioAgents) when required. Conidiospores were harvested and quantified as described in earlier studies (23).

### Bioinformatic tools

Information on gene annotations was obtained from the Ensembl Fungi database (38). Protein sequence of *V. dahliae* JR2 was obtained from the Ensembl Fungi database, whereas protein sequences of all other fungi mentioned in this study were obtained from the UniProt database (38, 39). Protein sequences were compared and phylograms were generated using the Clustal Omega multiple sequence alignment website (40).

### *Verticillium* mutant strain construction

Plasmids containing the deletion, complementation, or over expression cassette were transformed into *E. coli* DH5α by heat shock as established in prior studies (41), and the sequence-confirmed plasmids were transformed to *A. tumefaciens* AGL1 as mentioned previously (42). Plasmids were designed and constructed according to the principles of earlier studies, and the plasmids and primers are listed in S2 and S3 Table. *Agrobacterium*-mediated transformation was performed to manipulate the *V. dahliae* genome (6). The *ANN1* and *ANN2* deletion strains were constructed by replacing the open reading frame with a resistance cassette via homologous recombination. The complementation strains were generated by reintroducing the *ANN1* gene with *GFP* fused to the 5’ end, and the *ANN2* gene with *GFP* fused to the 3’ to the genomic locus of the respective deletion strains. The in locus over expression strains of *ANN1* and *ANN2* were generated by adding a strong *A. nidulans gpdA* promoter in the 5’ end of the respective genes in the respective genomic locus. The *ANN1* ectopic over expression strain was generated by ectopically integrating an extra copy of *ANN1* gene with *gpdA* promoter in the 5’ end in the WT background, whereas the *SOM1* over expression strain was generated by ectopically reintroducing the *SOM1* gene with *gpdA* promoter in the 5’ end in the *SOM1* deletion strain. Histone *H2B* with *RFP* fused to the 3’ end was ectopically integrated into the genome of the WT strain, a strain over expressing free Gfp, and the *ANN1* complementation strain for localisation studies. Genome of the mutant strains were verified by Southern hybridisation (S7 & S8 Fig.)

### RNA extraction and quantitative real-time PCR (qRT-PCR)

5 x 10^7^ conidiospores of each tested *V. dahliae* strain were inoculated in 50 mL of CDM, PDM, or SXM and incubated at 25°C. Mycelia of each tested strains were harvested 5 days post inoculation (5 dpi) and ground into powder. RNA was extracted from each mycelial sample with TRIzol (Invitrogen) (28). DNase treatments were performed on each crude RNA extract with TURBO DNA-free kit (Invitrogen), and cDNA was synthesised from DNase-treated RNA samples by RevertAid RT kit (Thermo Scientific) according to the manufacturer’s protocol. qRT-PCRs were performed to analyse transcript levels of *ANN1*, *ANN2*, and *ANN3* according to previously described methods (23), and the primers used are listed in S2 Table. Transcript levels of each tested mutant strain were normalised by the WT levels of the corresponding culture condition in the same biological replicate. One-sample t-test was performed for statistical analysis.

### Protein extraction and Western experiments

5 x 10^7^ conidiospores of each tested *V. dahliae* strain were inoculated into 50 mL of CDM, PDM, or SXM and incubated at 25°C for 5 days. For samples involving bacterial-fungal co-cultivations, *B. subtilis* 168 cultures were inoculated into 4-days old *V. dahliae* culture to reach a final *B. subtilis* 168 concentration of OD_600_ = 0.01, and fungal mycelia were harvested a day after the inoculation of bacterial culture. Mycelia of each sample were ground into powder and weighed. 500 µL of modified lysis buffer (10 mM Tris pH 7.5, 150 mM NaCl, 0.5 mM EDTA pH 8, 1 mM PMSF, 2 mM DTT, 0.5 % Nonidet-P40, 2x c0mplete™ EDTA-free Proteinase Inhibitor Cocktail, 4 % SDS) were added per 1 g of mycelial powder (43). Mycelial-lysis buffer mixtures were incubated at 65°C for 5 min, and centrifuged at 10,000 rpm for 20 min to separate protein extracts from cell debris. Protein concentration was determined by Bradford assay (28), and Western experiments and signal detection were done as previously described (37). Loading controls were either done by Ponceau staining, or by 4–20% Mini-PROTEAN TGX Stain-Free Protein Gels (Bio-Rad) with ChemiDoc Touch Imaging System (Bio-Rad) to ensure equal loading of proteins in each well.

### Confocal microscopy

The localisation of Gfp-fused Ann1 was visualised by fluorescence confocal microscopy with Gfp filters (35). Nuclei were visualised by ectopic expression of Rfp-fused histone H2b, and observed under the fluorescence confocal microscopy with Rfp filters (35).

### Phenotypic analysis

5 x 10^4^ of freshly harvested conidiospores were point-inoculated onto 30 mL agar plates. In addition to SXM, PDM, and CDM plates, CDM plates with 3% cellulose instead of sucrose as carbon source were used as a microsclerotia-inducing condition. CDM plates supplemented with 0.5 M NaCl or 0.8 M sorbitol were used for osmotic stress tests, and CDM plates supplemented with 20 µg/mL Congo red were used for cell wall stress tests. Plates were incubated at 25°C for 10 days and the bottom view of the plates was documented. Sizes of each documented colony were measured by the Fiji (ImageJ) software (44). Melanisation was measured as a representation of microsclerotia formation. Melanisation levels were quantified using Fiji (ImageJ) by converting images to 8-bit greyscale, adjusting the threshold to isolate melanised regions, and measuring the area using the ROI Manager tool (44). Colony size and melanisation level of each colony were normalised to the corresponding value of the WT colony within the same biological replicate. One-way ANOVA with post-hoc Tukey HSD test was performed for statistical analysis.

### Quantification of conidiation

Experiments performed to compare conidiation between different mutant strains were done as reported in earlier studies (23, 28). The amount of conidiospores counted from each strain was normalised by the WT levels in the same biological replicate. One-way ANOVA with post-hoc Tukey HSD test was performed for statistical analysis.

### Bacterial-fungal interaction assay

Bacterial-fungal interaction assays were performed by point-inoculating 5 x 10^4^ of freshly harvested conidiospores on 100 mL PDM agar plates. 10 µL of bacterial suspension at OD_600_ = 0.01 2.5 cm apart from the point of the fungal inoculation 3 days after inoculating the fungus. *E. coli* DH5⍺ or *B. subtilis* 168 were used to interact with the *V. dahliae* WT or *ANN1* mutant strains. Distances between the bacterial and the fungal colonies were measured 10 days after fungal inoculation. One-way ANOVA with post-hoc Tukey HSD test was performed for statistical analysis.

### Tomato plant infection

Pathogenicity of the *V. dahliae* WT and *ANN1* mutant strains was investigated by tomato plant infection experiments as described reported in earlier studies (18). Two-tailed Mann-Whitney U test was performed to compare the disease scores of seedlings inoculated WT with those inoculated with deionised water or with other *V. dahliae* strains.

### Secondary metabolite extraction and analysis

Two 30 mL CDM agar plates were each plated with 1 x 10^6^ of freshly harvested conidiospores and incubated for two weeks at 25°C. A total of 30 mL of agar were homogenised per tested strain, and metabolites were extracted from homogenised agar (28). Metabolite samples were measured and analysed as previously described (28, 45), and extracted ion chromatograms were generated by the FreeStyle 1.6 software (ThermoFisher Scientific). Each tested condition was at least performed in three biological replicates. MS2 spectra of the metabolites were compared between experiments to confirm the reproducibility of a certain compound in different experiments. MS2 spectra of all metabolites described in this study are listed in S4 Table.

## Acknowledgments

We thank K. Heimel, J. W. Kronstad, A. Nagel, A. Strohdiek for the fruitful discussions, N. Scheiter, K. Reimann, X. Jin, M. Bromm, and J. Reißmann for the support and technical assistance.

## Supporting information

**S1 Fig. The genomic structure of the core biosynthetic enzyme-encoding gene *ANN3*.**

**S2 Fig. *V. dahliae* Ann2 is not regulating *ANN3* in the tested conditions.**

**S3 Fig. *V. dahliae* Ann2 is involved in microsclerotia formation.**

**S4 Fig. The *V. dahliae* Ann1-regulated antibacterial activity has no significant impact on *E. coli* DH5⍺.**

**S5 Fig. Production of 15 metabolites are altered in the *V. dahliae SOM1* mutant strains.**

**S6 Fig. Production of 18 fungal metabolites are altered in the *V. dahliae ANN1* mutant strains during the co-cultivation with *B. subtilis* 168.**

**S7 Fig. Verification of the *V. dahliae ANN1* mutant strains.**

**S8 Fig. Verification of the *V. dahliae ANN2* mutant strains.**

**S1 Table. Bacterial and fungal strains used in this study.**

**S2 Table. Plasmids used in this study.**

**S3 Table. Primers used in this study.**

**S4 Table. MS2 spectra of all metabolites described in this study.**

## References

1. Fradin EF, Thomma BP. Physiology and molecular aspects of Verticillium wilt diseases caused by *V. dahliae* and *V. albo-atrum*. Mol Plant Pathol. 2006;7(2):71–86.

2. Wilhelm S. Longevity of the Verticillium Wilt Fungus in the Laboratory and Field. Phytopathology. 1955;45(3):180–1.

3. Hu DF, Wang CS, Tao F, Cui Q, Xu XM, Shang WJ, et al. Whole Genome Wide Expression Profiles on Germination of *Verticillium dahliae* Microsclerotia. Plos One. 2014;9(6).

4. Sarenqimuge S, Rahman S, Wang Y, von Tiedemann A. Dormancy and germination of microsclerotia of *Verticillium longisporum* are regulated by soil bacteria and soil moisture levels but not by nutrients. Front Microbiol. 2022;13:979218.

5. Sarenqimuge S, Wang Y, Alhussein M, Koopmann B, von Tiedemann A. The interplay of suppressive soil bacteria and plant root exudates determines germination of microsclerotia of *Verticillium longisporum*. Appl Environ Microbiol. 2024;90(6):e0058924.

6. Bui TT, Harting R, Braus-Stromeyer SA, Tran VT, Leonard M, Hofer A, et al. *Verticillium dahliae* transcription factors Som1 and Vta3 control microsclerotia formation and sequential steps of plant root penetration and colonisation to induce disease. New Phytol. 2019;221(4):2138–59.

7. Tran VT, Braus-Stromeyer SA, Kusch H, Reusche M, Kaever A, Kuhn A, et al. Verticillium transcription activator of adhesion Vta2 suppresses microsclerotia formation and is required for systemic infection of plant roots. New Phytol. 2014;202(2):565–81.

8. Harting R, Nagel A, Nesemann K, Hofer AM, Bastakis E, Kusch H, et al. *Pseudomonas* Strains Induce Transcriptional and Morphological Changes and Reduce Root Colonization of *Verticillium* spp. Front Microbiol. 2021;12:652468.

9. Künzler M. How fungi defend themselves against microbial competitors and animal predators. Plos Pathogens. 2018;14(9).

10. Stockli M, Morinaka BI, Lackner G, Kombrink A, Sieber R, Margot C, et al. Bacteria-induced production of the antibacterial sesquiterpene lagopodin B in *Coprinopsis cinerea*. Mol Microbiol. 2019;112(2):605–19.

11. Zhang T, Zhang B, Hua C, Meng P, Wang S, Chen Z, et al. *VdPKS1* is required for melanin formation and virulence in a cotton wilt pathogen *Verticillium dahliae*. Sci China Life Sci. 2017;60(8):868–79.

12. Brakhage AA. Regulation of fungal secondary metabolism. Nat Rev Microbiol. 2013;11(1):21–32.

13. Medema MH, Blin K, Cimermancic P, de Jager V, Zakrzewski P, Fischbach MA, et al. antiSMASH: rapid identification, annotation and analysis of secondary metabolite biosynthesis gene clusters in bacterial and fungal genome sequences. Nucleic Acids Res. 2011;39(Web Server issue):W339–46.

14. Keller NP. Fungal secondary metabolism: regulation, function and drug discovery. Nat Rev Microbiol. 2019;17(3):167–80.

15. Hollensteiner J, Wemheuer F, Harting R, Kolarzyk AM, Diaz Valerio SM, Poehlein A, et al. *Bacillus thuringiensis* and *Bacillus weihenstephanensis* Inhibit the Growth of Phytopathogenic *Verticillium* Species. Front Microbiol. 2016;7:2171.

16. Nesemann K, Braus-Stromeyer SA, Harting R, Hofer A, Kusch H, Ambrosio AB, et al. Fluorescent pseudomonads pursue media-dependent strategies to inhibit growth of pathogenic *Verticillium* fungi. Appl Microbiol Biotechnol. 2018;102(2):817–31.

17. Snelders NC, Rovenich H, Petti GC, Rocafort M, van den Berg GCM, Vorholt JA, et al. Microbiome manipulation by a soil-borne fungal plant pathogen using effector proteins. Nat Plants. 2020;6(11):1365–74.

18. Harting R, Hofer A, Tran VT, Weinhold LM, Barghahn S, Schluter R, et al. The Vta1 transcriptional regulator is required for microsclerotia melanization in *Verticillium dahliae*. Fungal Biol. 2020;124(5):490–500.

19. Shi-Kunne X, Jove RP, Depotter JRL, Ebert MK, Seidl MF, Thomma B. *In silico* prediction and characterisation of secondary metabolite clusters in the plant pathogenic fungus *Verticillium dahliae*. FEMS Microbiol Lett. 2019;366(7).

20. Wang Y, Hu X, Fang Y, Anchieta A, Goldman PH, Hernandez G, et al. Transcription factor VdCmr1 is required for pigment production, protection from UV irradiation, and regulates expression of melanin biosynthetic genes in *Verticillium dahliae*. Microbiology (Reading). 2018;164(4):685–96.

21. Xiong DG, Wang YL, Tian CM. A novel gene from a secondary metabolism gene cluster is required for microsclerotia formation and virulence in *Verticillium dahliae*. Phytopathology Research. 2019;1(1).

22. Luo X, Tian T, Tan X, Zheng Y, Xie C, Xu Y, et al. *VdNPS*, a nonribosomal peptide synthetase, is involved in regulating virulence in *Verticillium dahliae*. Phytopathology. 2020;110(8):1398–409.

23. Maurus I, Harting R, Herrfurth C, Starke J, Nagel A, Mohnike L, et al. *Verticillium dahliae* Vta3 promotes *ELV1* virulence factor gene expression in xylem sap, but tames Mtf1-mediated late stages of fungus-plant interactions and microsclerotia formation. PLoS Pathog. 2023;19(1):e1011100.

24. Maurus I, Leonard M, Nagel A, Starke J, Kronstad JW, Harting R, et al. Tomato Xylem Sap Hydrophobins Vdh4 and Vdh5 Are Important for Late Stages of *Verticillium dahliae* Plant Infection. J Fungi (Basel). 2022;8(12).

25. Murphy RL, Andrianopoulos A, Davis MA, Hynes MJ. Identification of *amdX*, a new Cys-2-His-2 (C2H2) zinc-finger gene involved in the regulation of the *amdS* gene of *Aspergillus nidulans*. Molecular Microbiology. 1997;23(3):591–602.

26. Simon M, Adam G, Rapatz W, Spevak W, Ruis H. The *Saccharomyces cerevisiae ADR1* gene is a positive regulator of transcription of genes encoding peroxisomal proteins. Mol Cell Biol. 1991;11(2):699–704.

27. Turcotte B, Liang XB, Robert F, Soontorngun N. Transcriptional regulation of nonfermentable carbon utilization in budding yeast. FEMS Yeast Res. 2010;10(1):2–13.

28. Hofer AM, Harting R, Assmann NF, Gerke J, Schmitt K, Starke J, et al. The velvet protein Vel1 controls initial plant root colonization and conidia formation for xylem distribution in Verticillium wilt. PLoS Genet. 2021;17(3):e1009434.

29. Leonard M, Kuhn A, Harting R, Maurus I, Nagel A, Starke J, et al. *Verticillium longisporum* Elicits Media-Dependent Secretome Responses With Capacity to Distinguish Between Plant-Related Environments. Front Microbiol. 2020;11:1876.

30. Georgianna DR, Payne GA. Genetic regulation of aflatoxin biosynthesis: from gene to genome. Fungal Genet Biol. 2009;46(2):113–25.

31. Davis MA, Kelly JM, Hynes MJ. Fungal catabolic gene regulation: molecular genetic analysis of the *amdS* gene of *Aspergillus nidulans*. Genetica. 1993;90(2-3):133–45.

32. Bemis LT, Denis CL. Identification of functional regions in the yeast transcriptional activator *ADR1*. Mol Cell Biol. 1988;8(5):2125–31.

33. Carroll CL, Carter CA, Goodhue RE, Lawell CCL, Subbarao KV. A Review of Control Options and Externalities for Verticillium Wilts. Phytopathology. 2018;108(2):160–71.

34. Olivain C, Humbert C, Nahalkova J, Fatehi J, L’Haridon F, Alabouvette C. Colonization of tomato root by pathogenic and nonpathogenic *Fusarium oxysporum* strains inoculated together and separately into the soil. Appl Environ Microb. 2006;72(2):1523–31.

35. Nagel A, Leonard M, Maurus I, Starke J, Schmitt K, Valerius O, et al. The Frq-Frh Complex Light-Dependently Delays Sfl1-Induced Microsclerotia Formation in *Verticillium dahliae*. J Fungi (Basel). 2023;9(7).

36. Bertani G. Studies on lysogenesis. I. The mode of phage liberation by lysogenic *Escherichia coli*. J Bacteriol. 1951;62(3):293–300.

37. Starke J, Harting R, Maurus I, Leonard M, Bremenkamp R, Heimel K, et al. Unfolded Protein Response and Scaffold Independent Pheromone MAP Kinase Signaling Control *Verticillium dahliae* Growth, Development, and Plant Pathogenesis. J Fungi (Basel). 2021;7(4).

38. Yates AD, Allen J, Amode RM, Azov AG, Barba M, Becerra A, et al. Ensembl Genomes 2022: an expanding genome resource for non-vertebrates. Nucleic Acids Res. 2022;50(D1):D996–D1003.

39. UniProt C. UniProt: the Universal Protein Knowledgebase in 2025. Nucleic Acids Res. 2025;53(D1):D609–D17.

40. Madeira F, Madhusoodanan N, Lee J, Eusebi A, Niewielska A, Tivey ARN, et al. The EMBL-EBI Job Dispatcher sequence analysis tools framework in 2024. Nucleic Acids Res. 2024;52(W1):W521–W5.

41. Hanahan D, Jessee J, Bloom FR. Plasmid transformation of *Escherichia coli* and other bacteria. Methods Enzymol. 1991;204:63–113.

42. Jyothishwaran G, Kotresha D, Selvaraj T, Srideshikan SM, Rajvanshi PK, Jayabaskaran C. A modified freeze-thaw method for efficient transformation of *Agrobacterium tumefaciens*. Curr Sci India. 2007;93(6):770–2.

43. Hollstein LS, Groth S, Schmitt K, Valerius O, Pöggeler S. *In vivo* proximity-labeling with miniTurboID to screen for protein-protein interactions in the filamentous ascomycete *Sordaria macrospora*. MethodsX. 2025;14:103351.

44. Schindelin J, Arganda-Carreras I, Frise E, Kaynig V, Longair M, Pietzsch T, et al. Fiji: an open-source platform for biological-image analysis. Nat Methods. 2012;9(7):676–82.

45. Kohler AM, Harting R, Langeneckert AE, Valerius O, Gerke J, Meister C, et al. Integration of Fungus-Specific CandA-C1 into a Trimeric CandA Complex Allowed Splitting of the Gene for the Conserved Receptor Exchange Factor of CullinA E3 Ubiquitin Ligases in *Aspergilli*. mBio. 2019;10(3).

