## Supporting figure 1 for "The adapt-to-nutrient NRPS-like secondary metabolite gene cluster facilitates *Verticillium dahliae* adaptation to different nutrient environments"

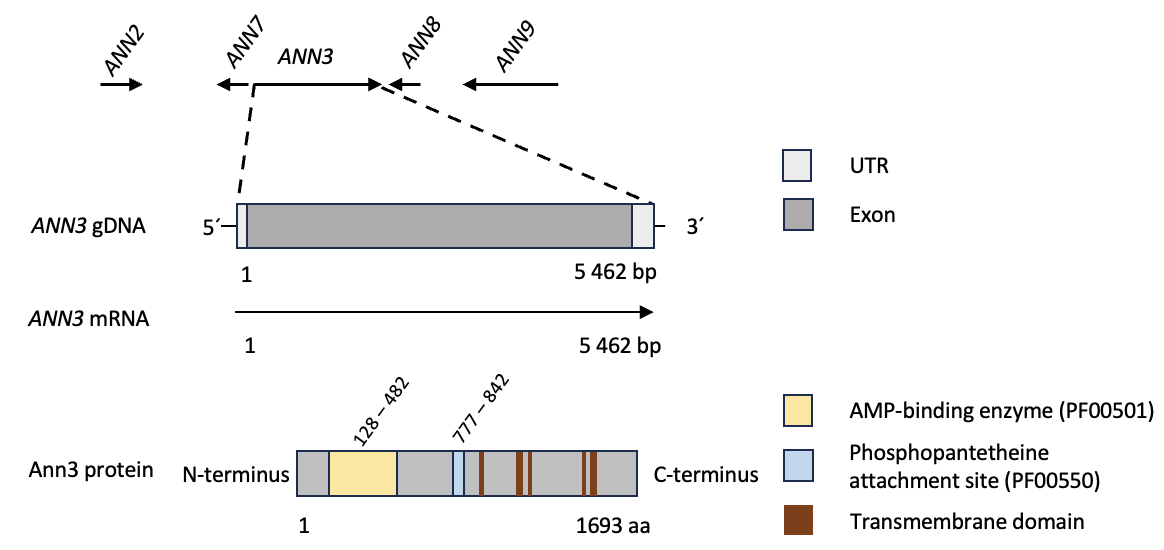

**S1 Fig. The genomic structure of the core biosynthetic enzyme-encoding gene *ANN3*.** The 5462 bp *ANN3* gene contains 1 exon (dark grey), and 5’- and 3’ UTRs (light grey). The 1693 aa Ann3 protein contains an AMP-binding enzyme domain (PF00501; 128 – 482 aa; yellow), a Phosphopantetheine attachment site (PF00550; 777 – 842 aa; light blue), and five Transmembrane domains (919 – 940, 1102 – 1129, 1154 – 1177, 1418 – 1439, 1468 – 1496; brown).
