## Supporting figure 2 for "The adapt-to-nutrient NRPS-like secondary metabolite gene cluster facilitates *Verticillium dahliae* adaptation to different nutrient environments"

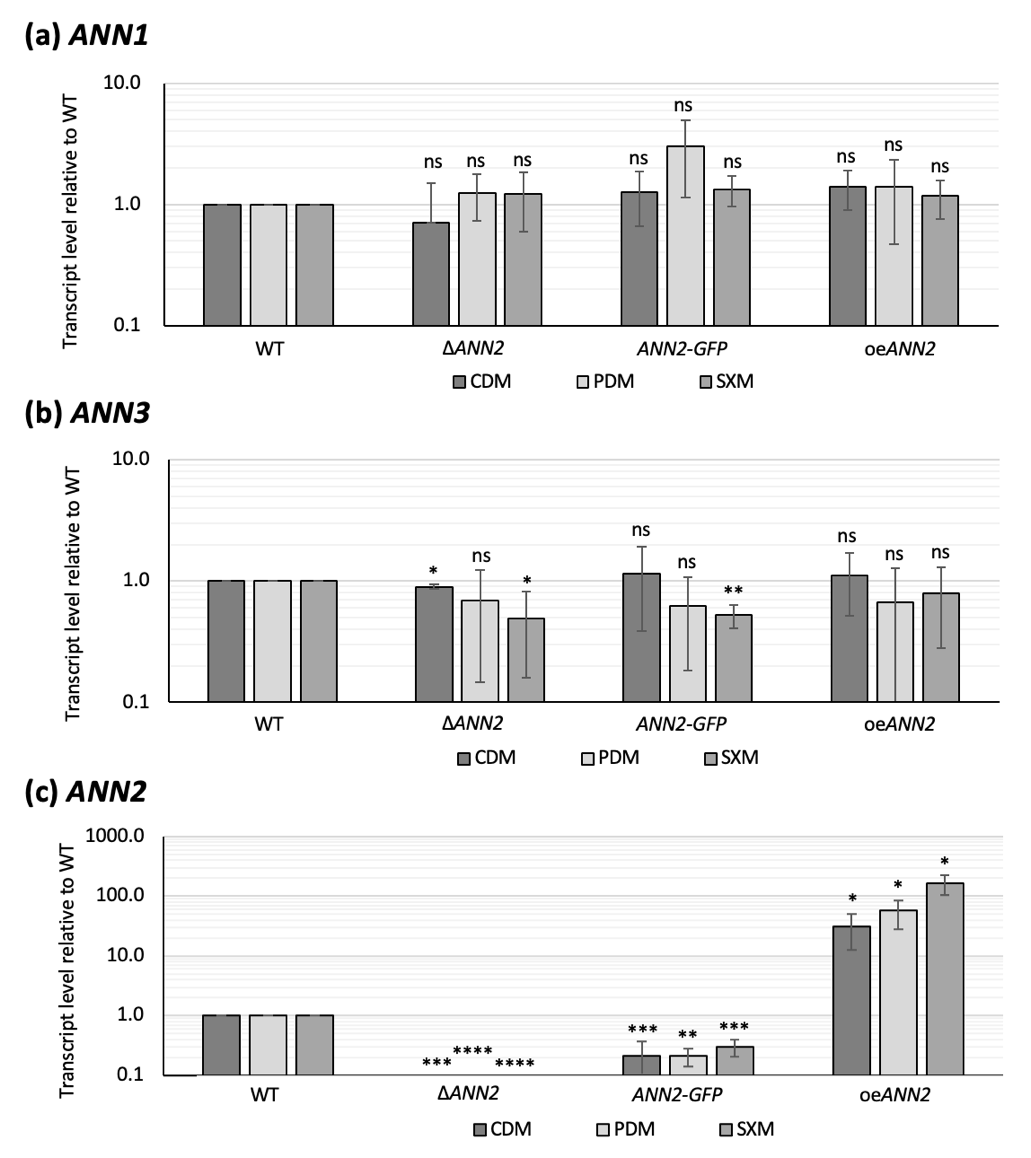

**S2 Fig. *V. dahliae* Ann2 is not regulating *ANN3* in the tested conditions.** Transcript levels of (a) *ANN1*, (b) *ANN3*, and (c) *ANN2* in the *ANN2* mutant strains were analysed by qRT-PCR. *ANN2* transcript levels are elevated in the over expression strains. Compared to the WT expression levels, transcript levels of *ANN1* and *ANN3* did not change more than 2-fold in any tested media in any *ANN2* mutant strain. One-sample t-test was performed to compare transcript levels of each tested strain to the WT expression levels under the same culturing condition (*, P < 0.05; **, P < 0.01; ***, P < 0.001; ****, P ≤ 0.0001; ns, not significantly different).
