## Supporting figure 3 for "The adapt-to-nutrient NRPS-like secondary metabolite gene cluster facilitates *Verticillium dahliae* adaptation to different nutrient environments"

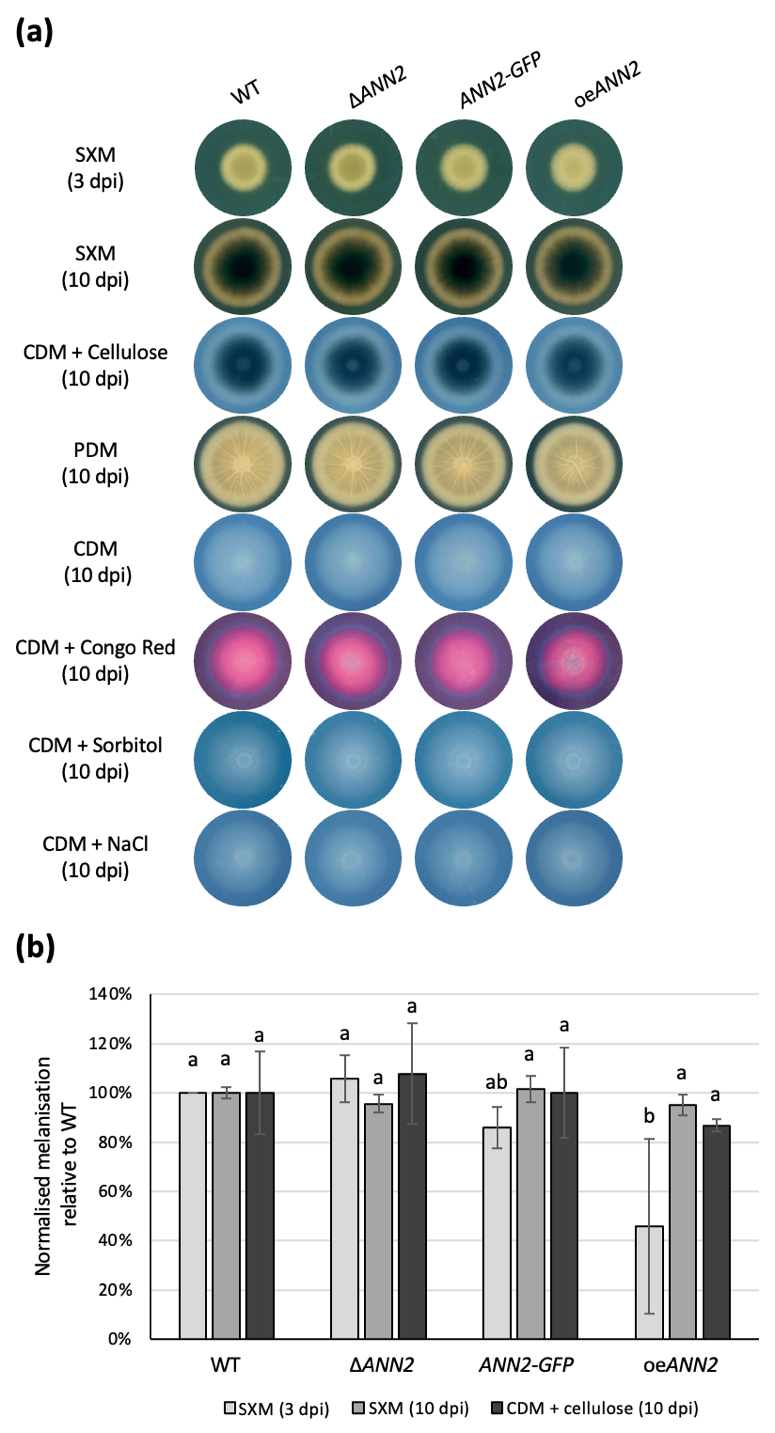

**S3 Fig. *V. dahliae* Ann2 plays a minor role in microsclerotia formation.** (a) The *ex planta* growth phenotype was compared between WT and the *ANN2* mutant strains on pectin-rich medium (SXM), minimal medium supplemented with cellulose (CDM + cellulose), potato dextrose medium (PDM), minimal medium (CDM), minimal medium supplemented with Congo red (CDM + Congo red), minimal medium supplemented with sorbitol (CDM + sorbitol), and minimal medium supplemented with NaCl (CDM + NaCl). 50,000 spores were dropped on each agar plate. SXM plates were scanned 3 dpi or 10 dpi, and all the other plates were scanned 10 dpi. (b) Melanisation was measured by Fiji ImageJ and normalised by the degree of melanisation of WT in the respective culture medium. Melanisation is delayed in the oe*ANN2* strain. The oe*ANN2* strain is less melanised than all other tested strains on SXM plates at 3 dpi, but no difference in melanisation level can be observed at 10 dpi. One-way ANOVA with post-hoc Tukey HSD test were performed to compare the colony size and the degree of melanisation of each strain under the same culture medium. A difference of the lower-case letter on top of each bar indicates significant difference (P < 0.05).
