## Supporting figure 4 for "The adapt-to-nutrient NRPS-like secondary metabolite gene cluster facilitates *Verticillium dahliae* adaptation to different nutrient environments"

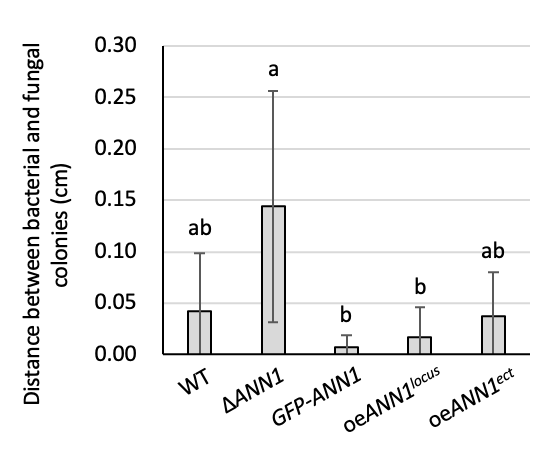

**S4 Fig. The *V. dahliae* Ann1-regulated antibacterial activity has no significant impact on *E. coli* DH5⍺.** The WT *V. dahliae* and the *ANN1* mutant strains were spotted on 100 mL PDM plates, and 10 µL of bacterial suspension at OD_600_ = 0.01 was spotted 2.5 cm apart from the centre of the *V. dahliae* colony at 3 dpi. The *V. dahliae* and bacterial colonies were co-cultured for an additional 7 days before the results were evaluated. Antibacterial activity of *V. dahliae* against *E. coli* DH5⍺ was quantified by the distance between the two colonies. One-way ANOVA with post-hoc Tukey HSD test were performed to compare the antibacterial activity of each strain towards *E. coli* DH5⍺. A difference of the lower-case letter on top of each bar indicates significant difference (P < 0.05).
