## Supporting figure 5 for "The adapt-to-nutrient NRPS-like secondary metabolite gene cluster facilitates *Verticillium dahliae* adaptation to different nutrient environments"

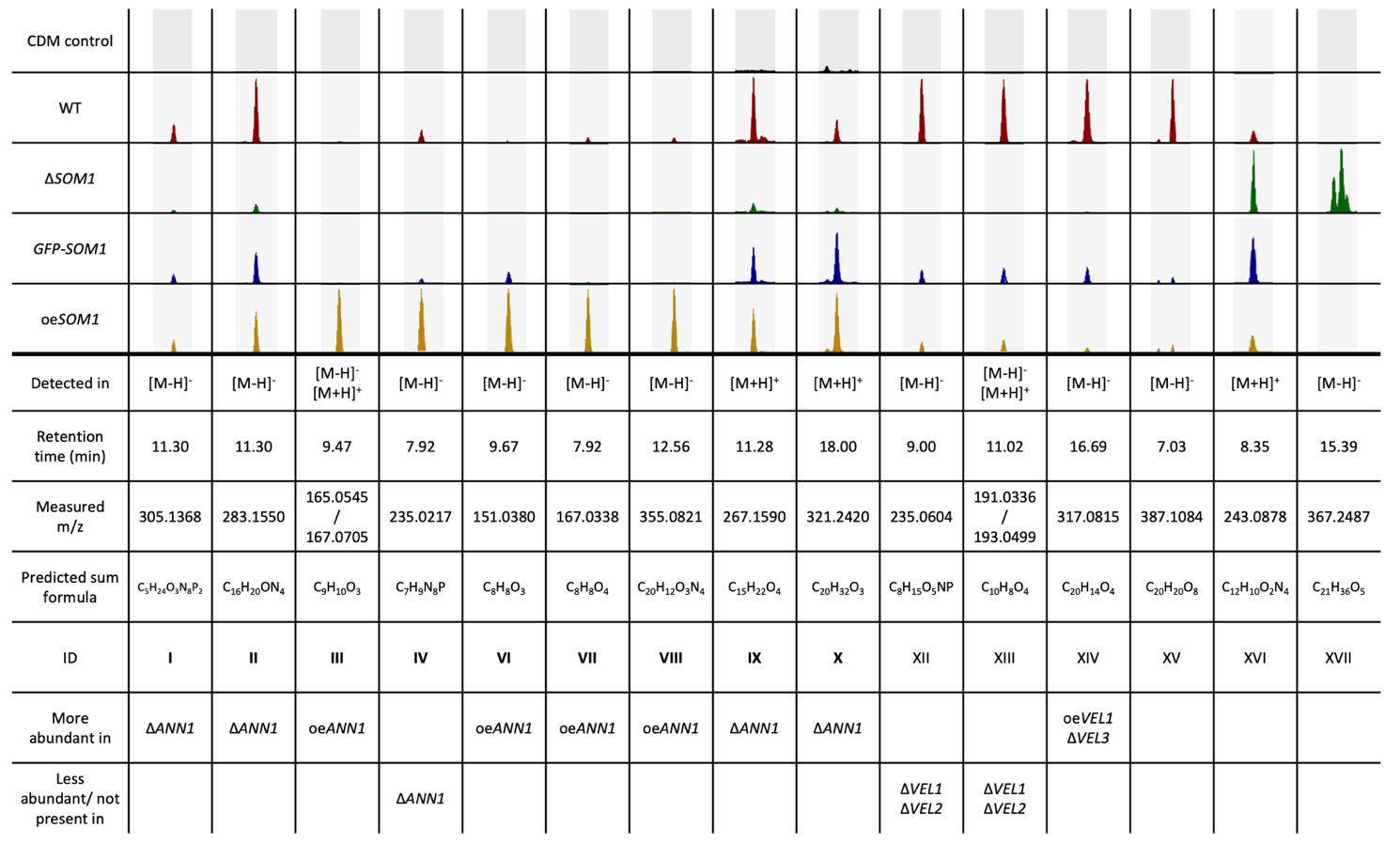

**S5 Fig. 15 metabolites had altered abundance in the *V. dahliae SOM1* mutant strains.** 13 of the listed metabolites (I) – (IV), (VI) – (X), (XII) – (XV) are either more abundant in the over expression *SOM1* strain, or less abundant in the ∆*SOM1* strain, whereas two metabolites are more abundant in the ∆*SOM1* strain (XVI) – (XVII). Extracted ion chromatogram of masses that had differed abundance in the tested strains are shown, and 5 ppm of mass deviation was tolerated. The height of each peak corresponds to the relative abundance of a certain mass in the tested strain. The predicted sum formula of each compound is calculated by the calculated exact mass. Nine of the detected masses (I) – (IV), (VI) – (X) (in **bold**) were also known to have altered abundance in *ANN1* mutant strain pure cultures, and three masses (XII) – (XIV) were known to have altered abundance in different velvet protein mutant strain pure cultures by comparing the respective MS2 spectra (1).

**References:**

1.	Hofer AM, Harting R, Assmann NF, Gerke J, Schmitt K, Starke J, et al. The velvet protein Vel1 controls initial plant root colonization and conidia formation for xylem distribution in Verticillium wilt. PLoS Genet. 2021;17(3):e1009434.
