## Supporting figure 6 for "The adapt-to-nutrient NRPS-like secondary metabolite gene cluster facilitates *Verticillium dahliae* adaptation to different nutrient environments"

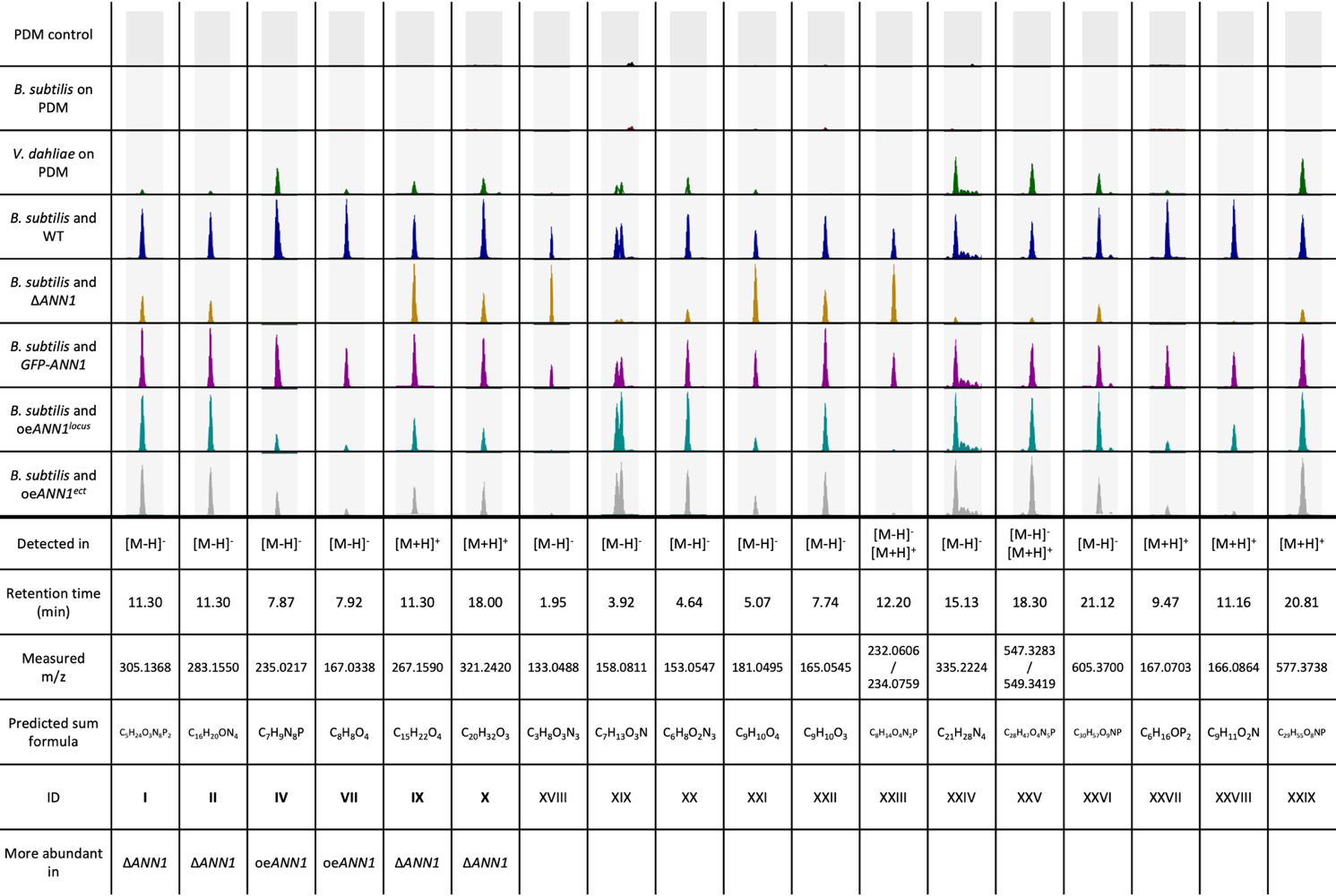

**S6 Fig. 18 fungal metabolites had altered abundance in the *V. dahliae* *ANN1* mutant strains during the co-cultivation with *B. subtilis* 168.** Extracted ion chromatogram of masses that had differed abundance in the tested strains are shown, and 5 ppm of mass deviation was tolerated. The height of each peak corresponds to the relative abundance of a certain mass in the tested strain. The predicted sum formula of each compound is calculated by the calculated exact mass. Six of the detected metabolites (I) (II), (IV), (VII), (IX), (X) (in **bold**) were also known to have altered abundance in *ANN1* mutant strain pure culture samples by comparing the respective MS2 spectra.
