## Supporting figure 7 for "The adapt-to-nutrient NRPS-like secondary metabolite gene cluster facilitates *Verticillium dahliae* adaptation to different nutrient environments"

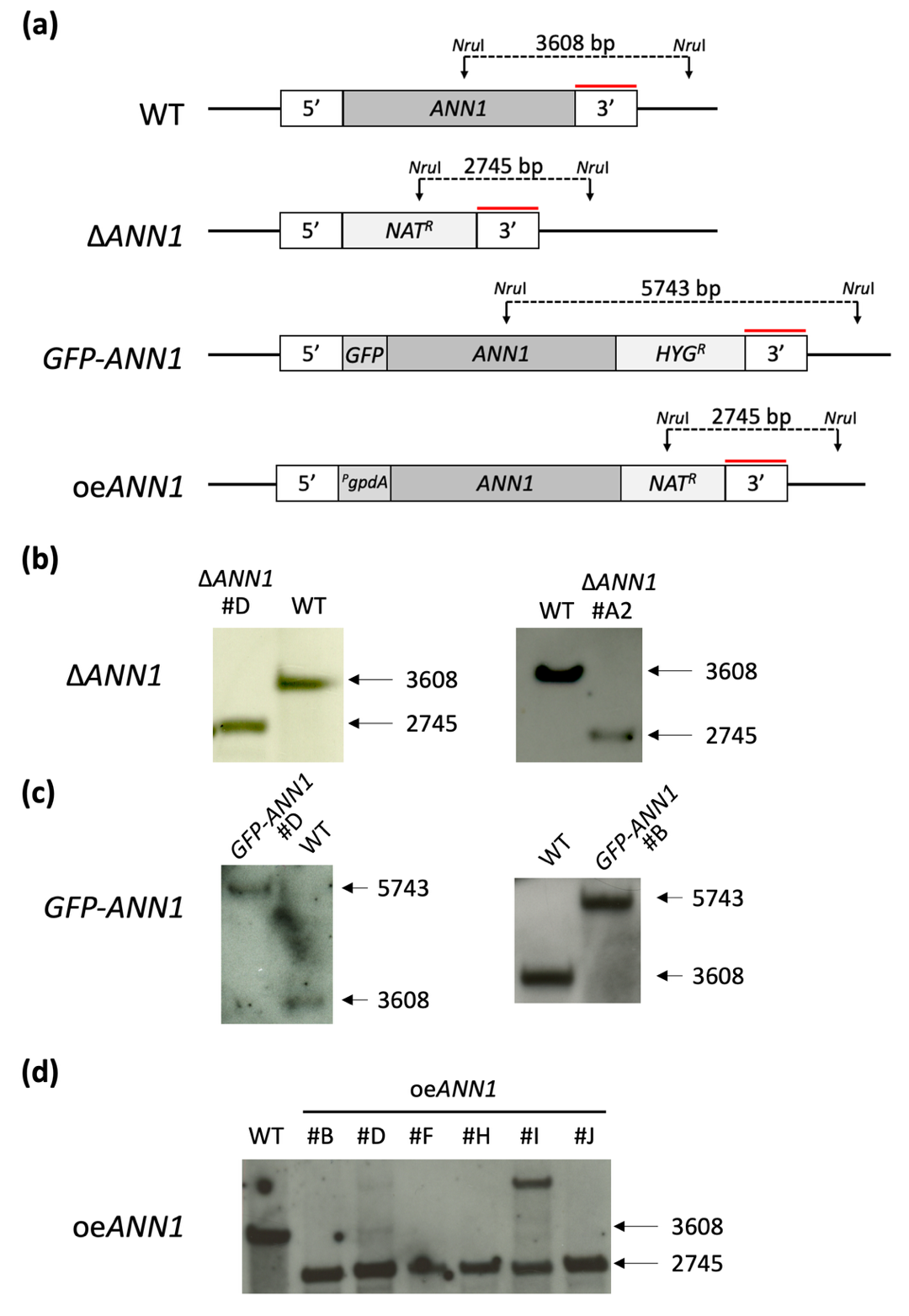

**S7 Fig**. **Verification of the *V. dahliae ANN1* mutant strains.** (a) Schemes of the genome of the WT and *ANN1* mutant strains and the mutant strains are depicted. *Nru*I ristriction sites are labelled in black arrows, and the probe that binds to the 3’ flanking region of *ANN1* are labelled in red line. The expected fragment sizes are written in the scheme. (b) The genome of ∆*ANN1* isolates D and A2 were confirmed to be correct. The WT strain served as control. (c) The genome of *GFP-ANN1* isolates D and B were confirmed. The WT served as control. (d) The genome of oe*ANN2* isolates H and J were confirmed. Isolates D and I were incorrect, and isolates B, D, F, and I were not used for further studies. The WT strain served as control.
