## Supporting figure 8 for "The adapt-to-nutrient NRPS-like secondary metabolite gene cluster facilitates *Verticillium dahliae* adaptation to different nutrient environments"

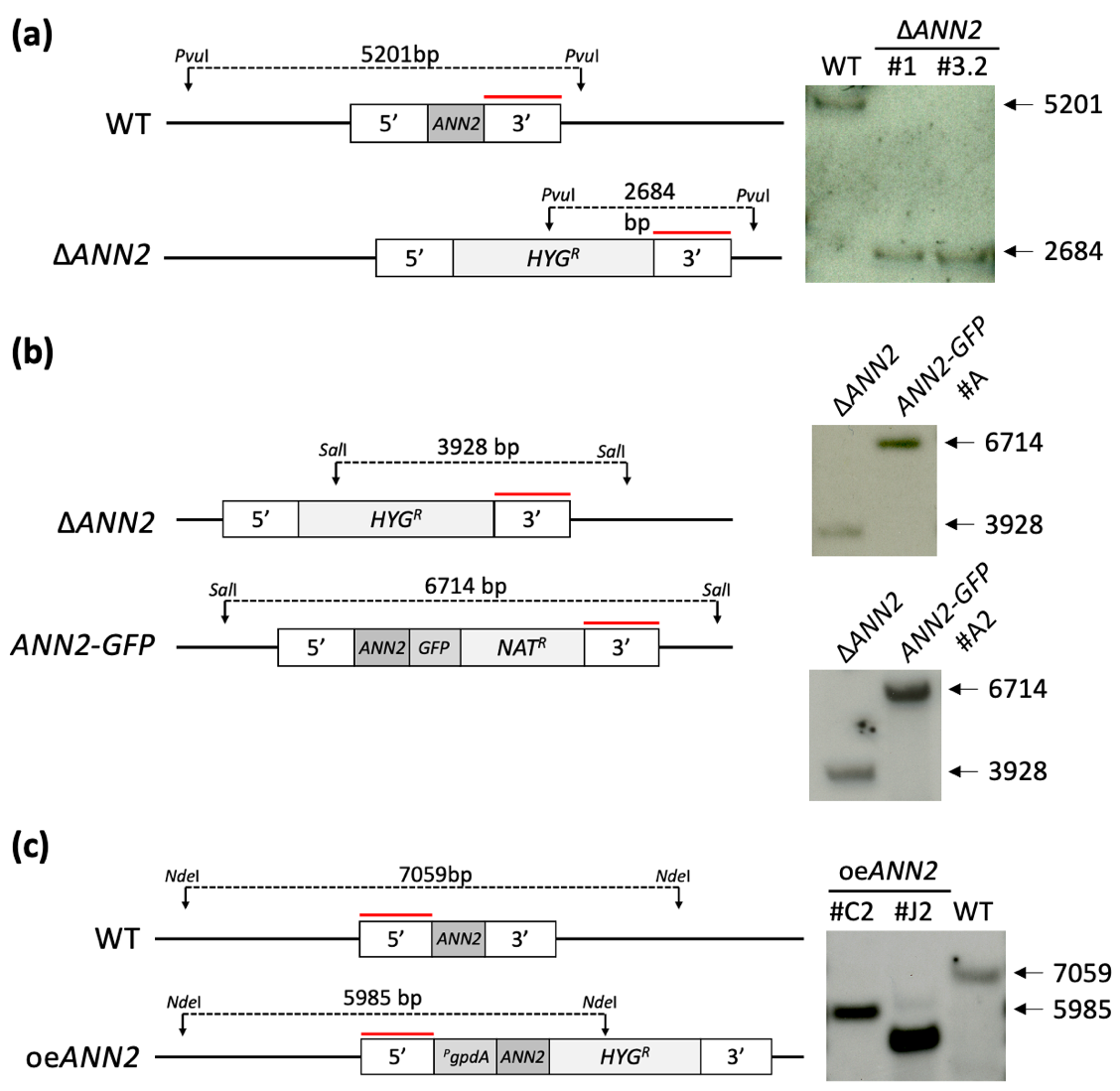

**S8 Fig**. **Verification of the *V. dahliae ANN2* mutant strains.** Schemes of the genome of the parental strains and the mutant strains are depicted in the left. Ristriction sites and the used restriction enzyme are labelled in black arrows, and the binding sites of the probes are labelled in red line. The expected fragment sizes are written in the scheme. (a) The genome of ∆*ANN2* isolates 1 and 3.2 were confirmed by restricting genomic DNA samples with *Pvu*I and hybridising with probe that binds to the 3’ flanking region of *ANN2*. The WT strain served as control. (b) The genome of *ANN2-GFP* isolates A and A2 were confirmed by restricting genomic DNA samples with *Sal*I and hybridising with probe that binds to the 3’ flanking region of *ANN2*. The parental ∆*ANN2* strain served as control. (c) The genome of oe*ANN2* isolate C2 were confirmed by restricting genomic DNA samples with *Nde*I and hybridising with probe that binds to the 5’ flanking region of *ANN2*. Isolate J2 was incorrect, and it was not used for further studies. The parental WT strain served as control.
