## Supporting table 1 for "The adapt-to-nutrient NRPS-like secondary metabolite gene cluster facilitates *Verticillium dahliae* adaptation to different nutrient environments"

**S1 Table**. **Bacterial and fungal strains used in this study.**

| **Strain name** | **Description** | **Reference** |
| --- | --- | --- |
| ***Escherichia coli*** | | |
| DH5α | Used for cloning and in bacterial-fungal interaction assays. | Invitrogen |
| ***Bacillus subtilis*** | | |
| 168 | Used in bacterial-fungal interaction assays. | (1) |
| ***Agrobacterium tumefaciens*** | | |
| AGL1 | Used for *Agrobacterium*-mediated *Verticillium* transformation. | (2) |
| ***Verticillium dahliae*** | | |
| JR2 (WT) | The WT *V. dahliae* strain isolated from *Solanum lycopersicum*. | (3) |
| VGB734/VGB740  (∆*ANN1*) | ∆*ANN1::^p^gpdA:nat^R^:trpC^t^* | This study |
| VGB742/VGB752  (*GFP-ANN1*) | ∆*ANN1::^p^ANN1:GFP:ANN1:^p^trpC:hyg^R^:trpC^t^* | This study |
| VGB735/VGB741  (oe*ANN1*^locus^) | *^p^gpdA:ANN1:^p^trpC:nat^R^:trpC^t^* | This study |
| VGB753  (oe*ANN1*^ect^) | *^p^gpdA:ANN1:^p^trpC:nat^R^:trpC^t^* | This study |
| VGB22  (*H2B-RFP*) | *^p^gpdA:histoneH2B:RFP:trpc^t^:^p^gpdA:nat^R^* | (4) |
| VGB23  (*H2B-RFP*) | *^p^gpdA:histoneH2B:RFP:trpc^t^:^p^gpdA:hyg^R^:trpC^t^* | This study |
| VGB768/VGB769  (*GFP*/*H2B-RFP*) | *^p^gpdA:GFP:trpC^t^:^p^gpdA:hyg^R^:trpC^t^*,  *^p^gpdA:histoneH2B:RFP:trpc^t^:^p^gpdA:nat^R^:trpC^t^* | This study |
| VGB770/VGB771  (*GFP-ANN1*/*H2B-RFP*) | ∆*ANN1::^p^ANN1:GFP:ANN1:^p^trpC:hyg^R^:trpC^t^*,  *^p^gpdA:histoneH2B:RFP:trpc^t^:^p^gpdA:nat^R^:trpC^t^* | This study |
| VGB763/VGB764  (∆*ANN2*) | ∆*ANN2::^p^gpdA:hyg^R^:trpC^t^* | This study |
| VGB765/VGB766  (*ANN2-GFP*) | ∆*ANN2::^p^ANN2:ANN2:GFP:^p^trpC:nat^R^:trpC^t^* | This study |
| VGB767  (oe*ANN2*) | *^p^gpdA:ANN2:^p^trpC:hyg^R^:trpC^t^* | This study |
| VGB001/VGB002  (∆*SOM1*) | ∆*SOM1::nat^R^* | (5) |
| VGB084/VGB085  (*SOM1-GFP*) | ∆*SOM1::nat^R^, ^p^SOM1:SOM1:GFP:trpC^t^*, *^p^gpdA:HPH:trpC^t^* | (5) |
| VGB176/VGB177  (oe*SOM1*) | ∆*SOM1::nat^R^*,  *^p^gpdA:SOM1:GFP:TrpC^t^:hyg^R^* | This study |

*^p^*: promoter**,** *^t^*: terminator, *hyg^R^*: hygromycin B resistance marker, *nat^R^*: nourseothricin resistance marker

**References:**

1.	Kunst F, Ogasawara N, Moszer I, Albertini AM, Alloni G, Azevedo V, et al. The complete genome sequence of the gram-positive bacterium *Bacillus subtilis*. Nature. 1997;390(6657):249-56.

2.	Lazo GR, Stein PA, Ludwig RA. A DNA transformation-competent *Arabidopsis* genomic library in *Agrobacterium*. Biotechnology (N Y). 1991;9(10):963-7.

3.	Fradin EF, Zhang Z, Juarez Ayala JC, Castroverde CD, Nazar RN, Robb J, et al. Genetic dissection of Verticillium wilt resistance mediated by tomato Ve1. Plant Physiol. 2009;150(1):320-32.

4.	Hofer AM, Harting R, Assmann NF, Gerke J, Schmitt K, Starke J, et al. The velvet protein Vel1 controls initial plant root colonization and conidia formation for xylem distribution in Verticillium wilt. PLoS Genet. 2021;17(3):e1009434.

5.	Bui TT, Harting R, Braus-Stromeyer SA, Tran VT, Leonard M, Hofer A, et al. *Verticillium dahliae* transcription factors Som1 and Vta3 control microsclerotia formation and sequential steps of plant root penetration and colonisation to induce disease. New Phytol. 2019;221(4):2138-59.
