## Supporting table 2 for "The adapt-to-nutrient NRPS-like secondary metabolite gene cluster facilitates *Verticillium dahliae* adaptation to different nutrient environments"

**S2 Table**. **Plasmids used in this study**

| **Plasmid** | **Description** | **Reference** |
| --- | --- | --- |
| pME4548 | *^p^trpC:nat^R^; kan^R^*  Cloning vector | (1) |
| pME4557 | *^p^gpdA:SOM1:GFP:trpC^t^*  Used for the generation of strain VGB176 and 177 | (1) |
| pME4975 | *^p^gpdA:H2B:RFP:trpC^t^:^P^gpdA:hyg^R^:trpC^t^*  Used for the generation of strain VGB23 | (2) |
| pYYC07 | *^p^ANN1: ^p^gpdA:nat^R^:trpC^t^:ANN1^t^*  Used for the generation of strain VGB734 and 740 | This study |
| pYYC09 | *^p^ANN1: ^p^gpdA:ANN1:^p^trpC:nat^R^:trpC^t^:ANN1^t^*  Used for the generation of strain VGB735 and 741 | This study |
| pYYC13 | *^p^ANN1:GFP:ANN1:^p^trpC:hyg^R^:trpC^t^:ANN1^t^*  Used for the generation of strain VGB742 and 752 | This study |
| pYYC15 | *^p^gpdA:ANN1: ^p^trpC:nat^R^:trpC^t^*  Used for the generation of strain VGB753 | This study |
| pYYC17 | *^p^ANN2:^p^gpdA:hyg^R^:trpC^t^:ANN2^t^*  Used for the generation of strain VGB763 and 764 | This study |
| pYYC18 | *^p^ANN2:ANN2:GFP:^p^trpC:nat^R^:trpC^t^:ANN2^t^*  Used for the generation of strain VGB765 and 766 | This study |
| pYYC19 | *^p^ANN2: ^p^gpdA:ANN2: ^p^trpC:hyg^R^:trpC^t^:ANN2^t^*  Used for the generation of strain VGB767 | This study |

**References:**

1.	Bui TT, Harting R, Braus-Stromeyer SA, Tran VT, Leonard M, Hofer A, et al. *Verticillium dahliae* transcription factors Som1 and Vta3 control microsclerotia formation and sequential steps of plant root penetration and colonisation to induce disease. New Phytol. 2019;221(4):2138-59.

2.	Starke J, Harting R, Maurus I, Leonard M, Bremenkamp R, Heimel K, et al. Unfolded Protein Response and Scaffold Independent Pheromone MAP Kinase Signaling Control *Verticillium dahliae* Growth, Development, and Plant Pathogenesis. J Fungi (Basel). 2021;7(4).
