## Supporting table 3 for "The adapt-to-nutrient NRPS-like secondary metabolite gene cluster facilitates *Verticillium dahliae* adaptation to different nutrient environments"

**S3 Table**. **Primers used in this study**

| **Name** | **Sequence** | **Target gene** |
| --- | --- | --- |
| **qRT-PCR primers** | | |
| KR97 | CGAAAATGCCTCGTCAATCACT | *ANN1* (VDAG_JR2_Chr5g11420a) |
| KR96 | TCTCGTGACGTTTCAGATGCT | *ANN1* (VDAG_JR2_Chr5g11420a) |
| YYC98 | TCAATTTGCAGCCTGTTGAGGTC | *ANN3* (VDAG_JR2_Chr5g11480a) |
| YYC99 | AGTCTCATGTTCACCGAGCC | *ANN3* (VDAG_JR2_Chr5g11480a) |
| YYC118 | AGGCGACATTCGACCACTG | *ANN2* (VDAG_JR2_Chr5g11460a) |
| YYC119 | TACGCCATGATCTGCTAGGG | *ANN2* (VDAG_JR2_Chr5g11460a) |
| SZ9 | AACACCCAGAACAAGATGCGC | *H2A* (VDAG_JR2_Chr4g01430a) |
| SZ10 | GCTTGACCTTGAGATCCTTG | *H2A* (VDAG_JR2_Chr4g01430a) |
| SZ11 | TGCATTCTTGGCAAGAGATGTGTG | *EIF2B* (VDAG_JR2_Chr4g00410a) |
| SZ12 | AGCTTGTTATCCTTGTCCTCGGT | *EIF2B* (VDAG_JR2_Chr4g00410a) |
| **Cloning primers** | | |
| KR05 | ATTCTTAATTAAGATTTTACATGTACAGCACGCTG | 5' flanking region of *ANN1*, overhang of pME4548 vector backbone |
| KR06 | ACCGGTCACTGTACATATTTTCGAAAGGTCGGGTC | 5' flanking region of *ANN1*, overhang of *Nat* resistant cassette |
| KR07 | AGGTAATCCTTCTTTTGGAGCAAAAAGCAAGGAGG | 3' flanking region of *ANN1*, overhang of *TrpC* terminator |
| KR08 | AGGACTTCTAGAAGGACGTGGCGCTCGAAAGTA | 3' flanking region of *ANN1*, overhang of pME4548 vector backbone |
| ML8 | AAAGAAGGATTACCTCTAAACAA | 3' end of *TrpC* terminator |
| KR11 | TGAGCAGACATCACCATGACCATGAACGAAGCCGC | *ANN1*, overhang of *gpdA* promoter |
| KR12 | CAGTTAACGTCGCGGTTACTCGCAAGCCATGACCG | *ANN1*, overhang of *trpC* promoter |
| YYC35 | CCGCGACGTTAACTGATATTGA | *trpC* promoter |
| YYC74 | CAGTTAACGTCGCGGTTACTTGTACAGCTCGTCCATGC | *GFP*, overhang of STOP and *TrpC* promoter |
| YYC76 | GCCCTTGCTCACCATTATTTTCGAAAGGTCGGGTCG | 5' flanking region of *ANN1*, overhang of *GFP* |
| YYC77 | GGTGGTAGCGGTGGTACCATGAACGAAGCCGCAAT | *ANN1*, overhang of *GFP* linker |
| YYC78 | GACCTTTCGAAAATAATGACCATGAACGAAGCCGC | *ANN1*, overhang of 5' flanking region |
| YYC79 | ACCACCGCTACCACCCTCGCAAGCCATGACCGAC | *ANN1*, overhang of *GFP* linker |
| YYC80 | GGTGGTAGCGGTGGTGTGAGCAAGGGCGAGGAG | *GFP*, overhang of linker |
| YYC86 | ATTCTTAATTAAGATTGTACAGTGACCGGTGAC | *gpdA* promotor, overhang pME4548 |
| YYC87 | AGGACTTCTAGAAGGAAAGAAGGATTACCTCTAAACAAG | *trpC* treminator, overhang of pME4548 |
| YYC83 | TTACTCGCAAGCCATGACCG | *ANN1* |
| YYC110 | GTATACATTTGGCATTCCAGGTTCTCGCAAGGC | 5' flanking region of *ANN2*, overhang *ANN2* |
| YYC111 | ATGCCAAATGTATACCCTCA | *ANN2* |
| YYC112 | ACCACCGCTACCACCGCTATAAGTCATCGGATTATCTA | *ANN2*, overhang linker |
| YYC113 | TGAGCAGACATCACCATGCCAAATGTATACCCTCA | *ANN2*, overhang *gpdA* promoter |
| YYC114 | CAGTTAACGTCGCGGTCAGCTATAAGTCATCGGATTATC | *ANN2*, overhang linker *trpC* promoter |
| So1 | ATTCTTAATTAAGATGTCTGGCTCAAACGGTGACG | 5' flanking region of *ANN2*, overhang of pME4548 vector backbone |
| So2 | ACCGGTCACTGTACATCCAGGTTCTCGCAAGGC | 5' flanking region of *ANN2*, overhang of *gpdA* promoter |
| So3 | AGGTAATCCTTCTTTACATGGAGAATGACATCGCG | 3' flanking region of *ANN2*, overhang of t*rpC* terminator |
| So4 | AGGACTTCTAGAAGGTTTCTCAAGGTACTTTTCGATTTGC | 3' flanking region of *ANN2*, overhang of pME4548 vector backbone |
| ML9 | TGTACAGTGACCGGTGAC | 5' end of *gpdA* promoter |
| ML31 | GGTGATGTCTGCTCAAGCGG | *gpdA* promoter |
| RH518 | ATGGTGAGCAAGGGCGAGG | *GFP* |
| RH519 | ACCACCGCTACCACCCTTGTACAGCTCGTCCATGC | *GFP*, overhang linker |
| AN45 | GTCGAGGGTGGCCATATCGATGCTTGGGTAGAAT | *trpC* promoter, overhang *Nat* resistant gene |
| AN46 | ATGGCCACCCTCGACG | *Nat* resistant gene |
| AN47 | AAAGAAGGATTACCTCTAAACAAGTGTA | *trpC* terminator |
| **Primers for sequencing** | | |
| YYC47 | CACTTCATCGCAGCTTGACT | *gpdA* promoter |
| YYC51 | CGGACACGCTGAACTTGT | *GFP* |
| YYC52 | CACTACCTGAGCACCCAGT | *GFP* |
| YYC57 | GCCAATTTGCCCATGCAT | *ANN1* |
| YYC58 | GGTGTCAACCATTACCGAGG | *ANN1* |
| YYC59 | CTGGTGACAGCGTACAACC | *ANN1* |
| SZ6 | GCGATTAAGTTGGGTAACGC | pME4548 plasmid backbone 3'- end |
| IM75 | GCTGATCTGACCAGTTGCCT | *trpC* promoter |
| RH463 | TCAACATGCTACCCTCCGC | pME4548 plasmid backbone 5'- end |
| RH615 | CATCCACTGCACCTCAGAG | *gpdA* promoter |
| RH658 | CAGGACACACATTCATCGTAG | *TrpC* terminator |
