## Supporting table 4 for "The adapt-to-nutrient NRPS-like secondary metabolite gene cluster facilitates *Verticillium dahliae* adaptation to different nutrient environments"

**S4 Table**. **MS2 spectra of all metabolites described in this study**

| MS2 305.1368, negative  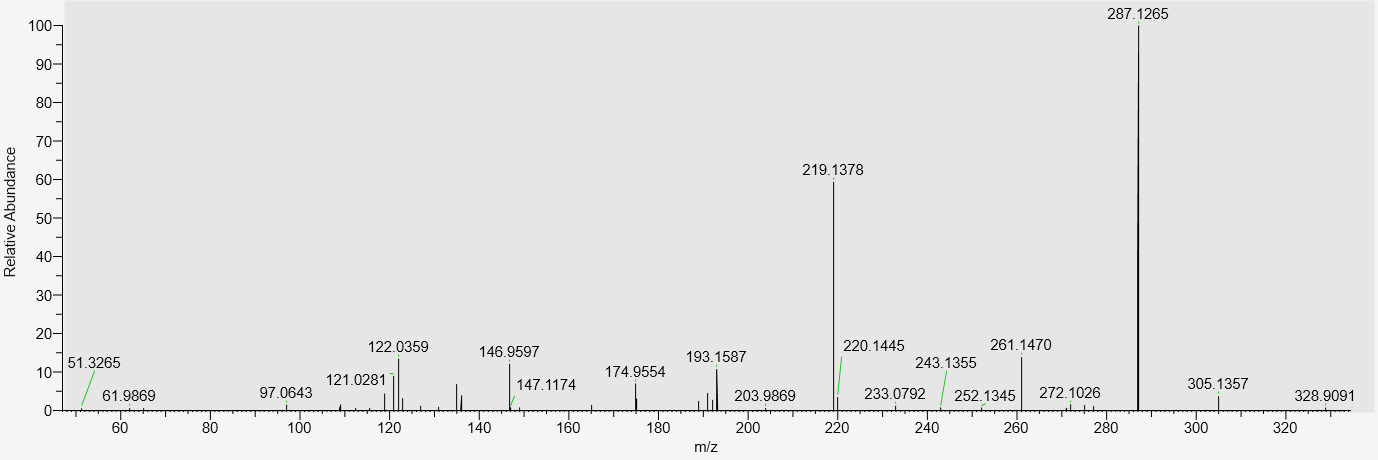  Substance I | MS2 283.1550, negative  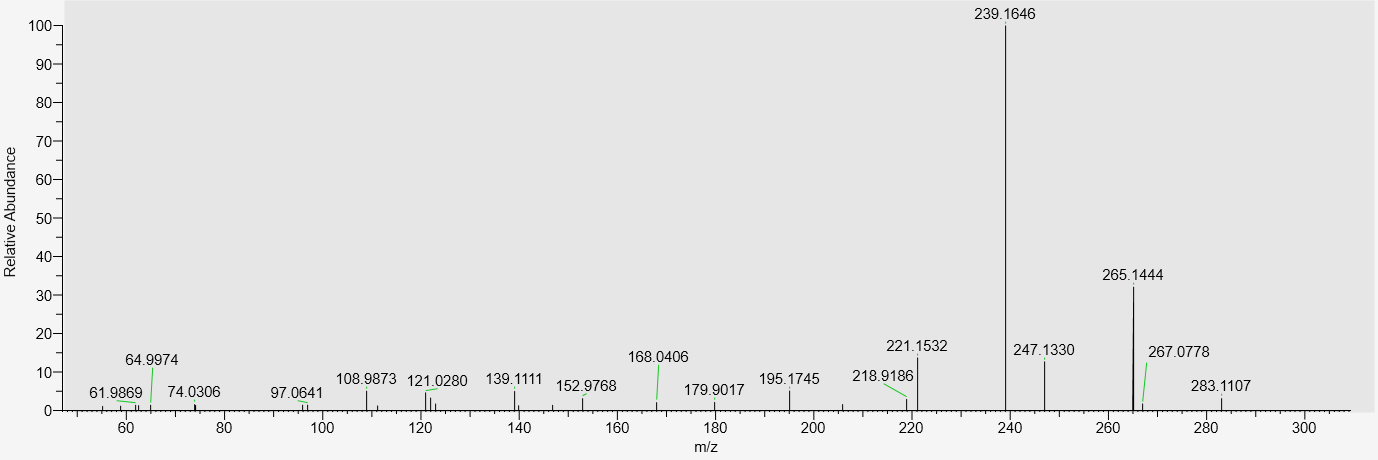  Substance II |
| --- | --- |
| MS2 165.0545, negative  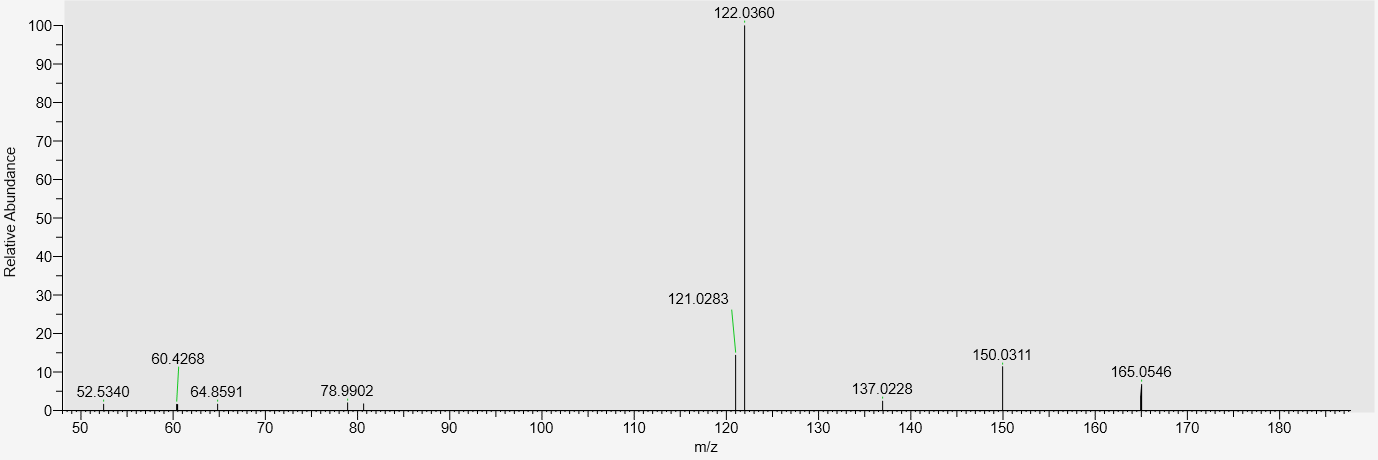  Substance III | MS2 167.0703, positive  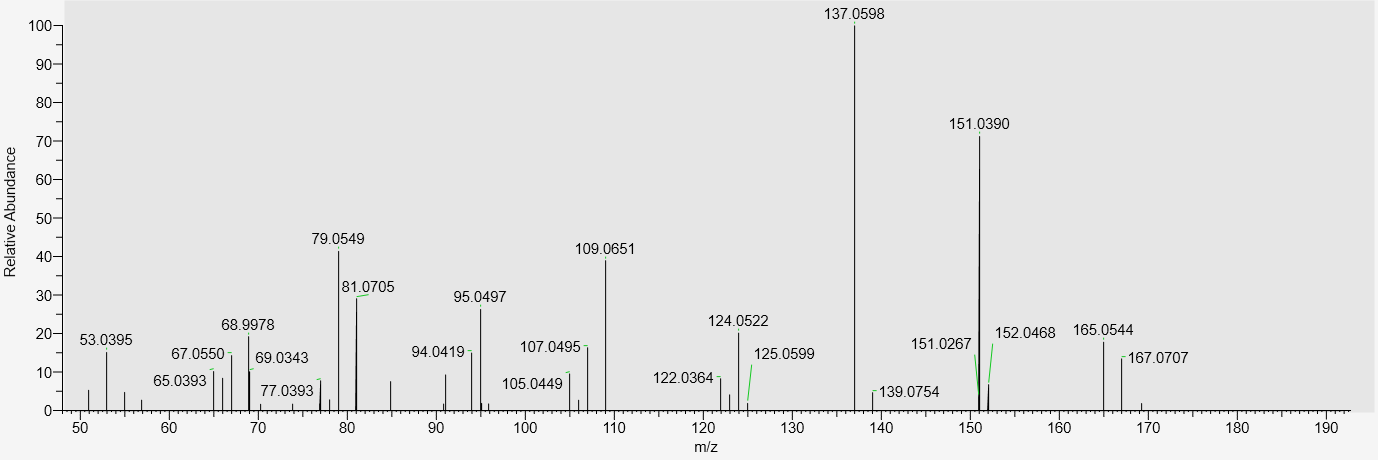 |
| MS2 235.0217, negative  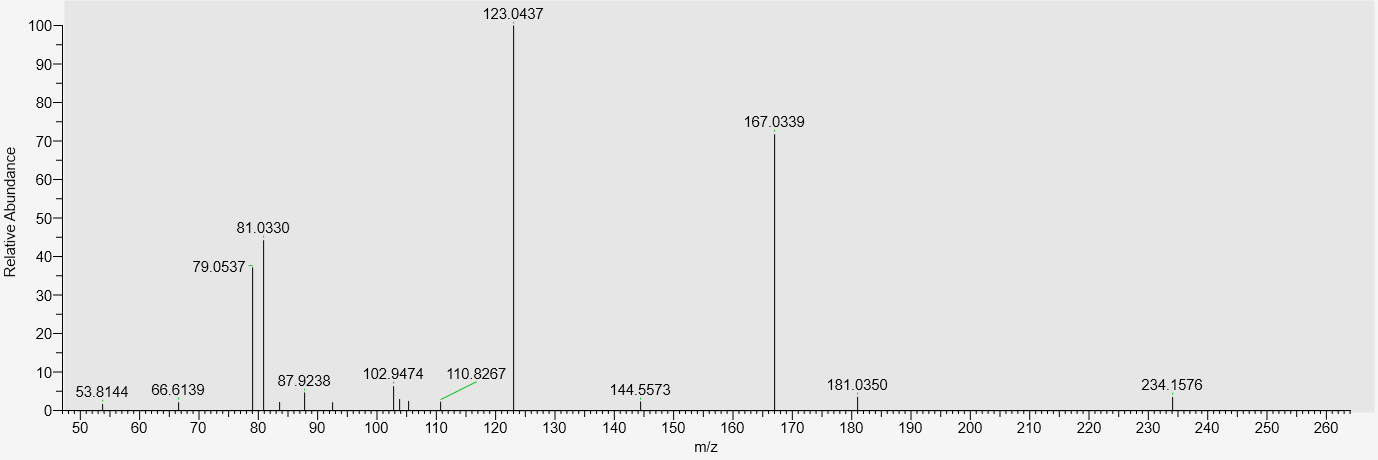  Substance IV | MS2 141.0540, negative  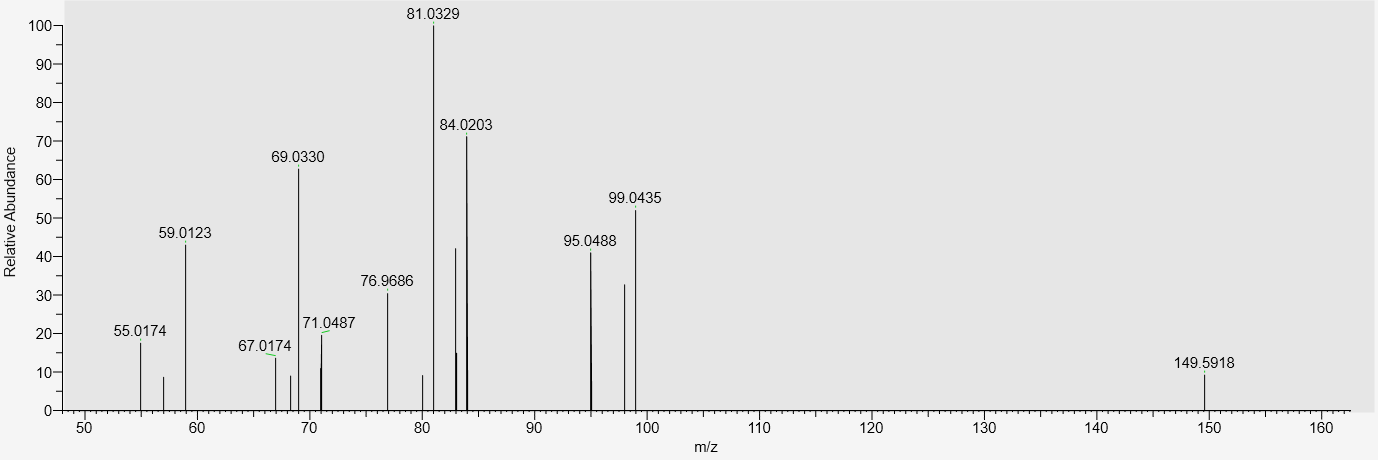  Substance V |

| MS2 151.0380, negative  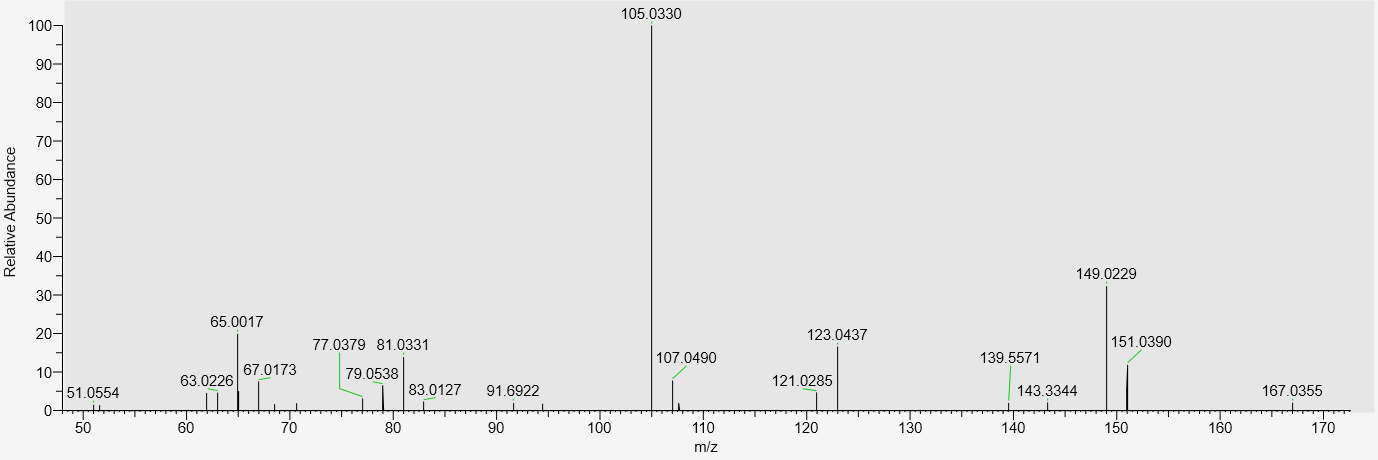  Substance VI | MS2 153.0547, positive  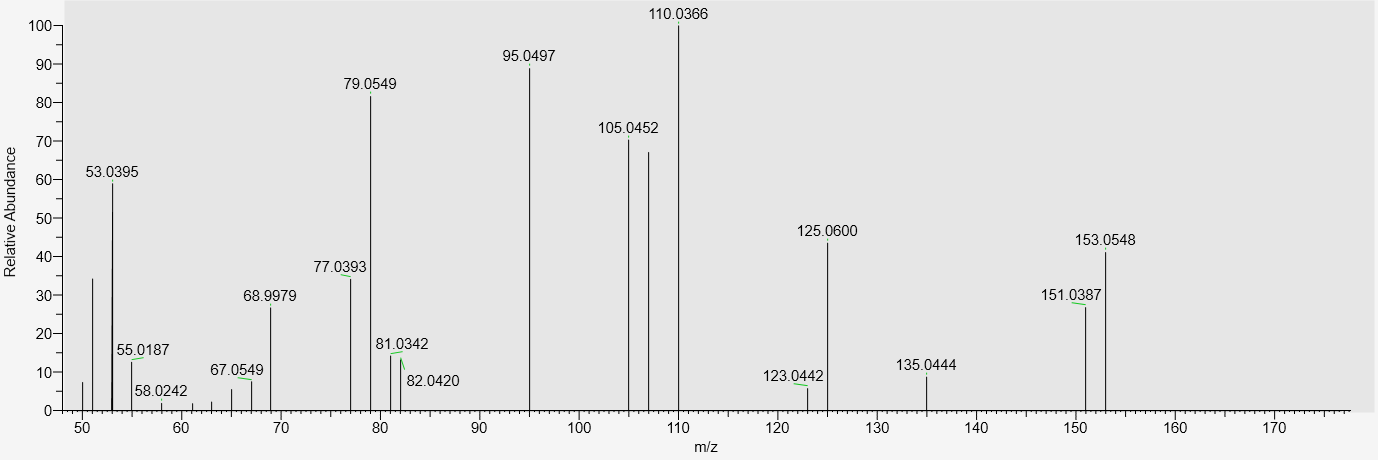 |
| --- | --- |
| MS2 167.0338, negative  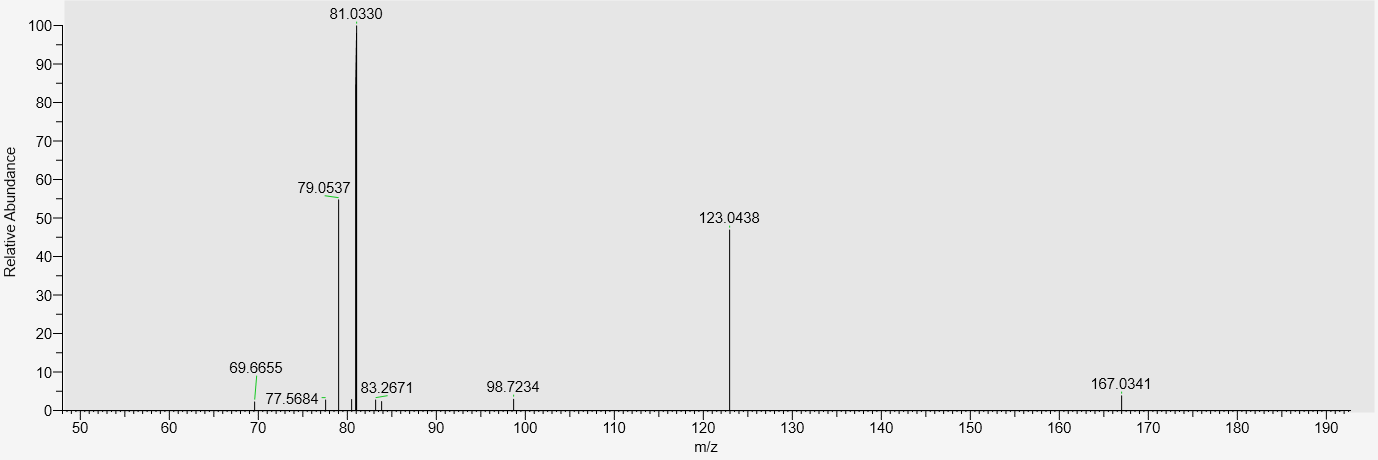  Substance VII | MS2 355.0821, negative  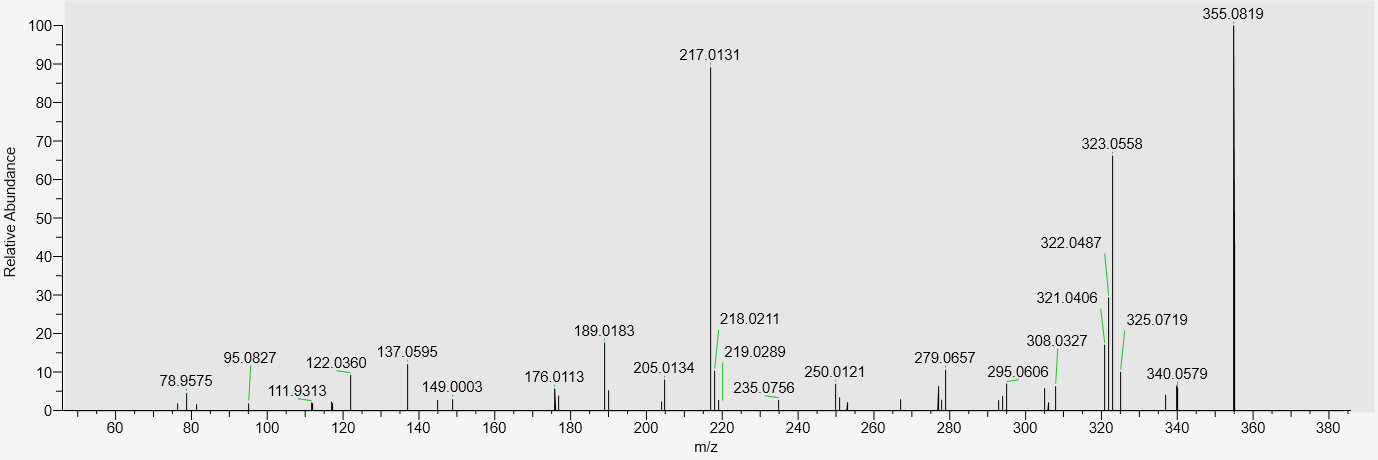  Substance VIII |
| MS2 267.1590, positive  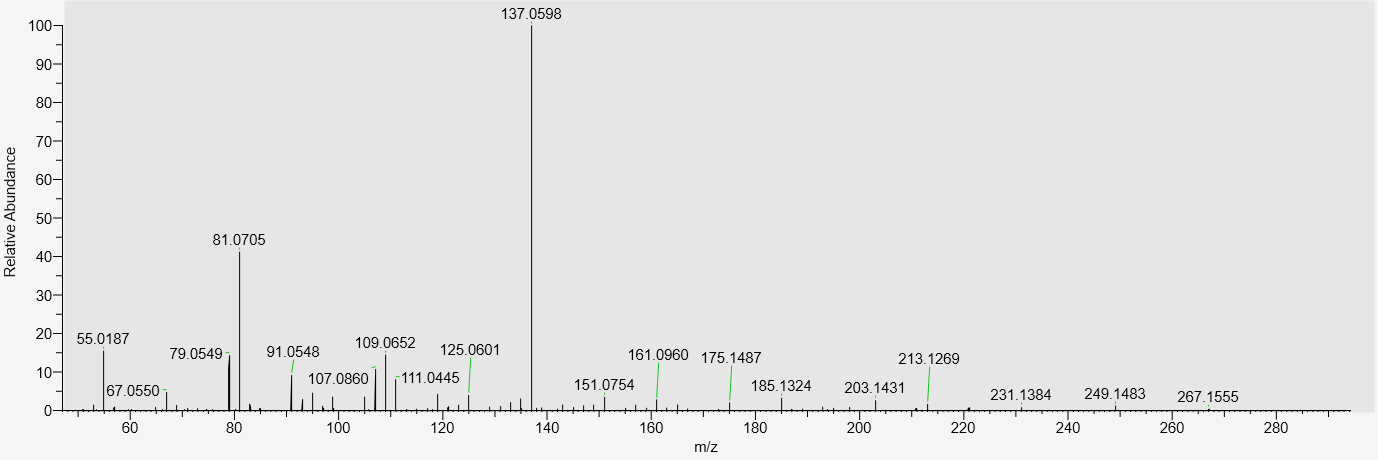  Substance IX | MS2 321.2420, positive  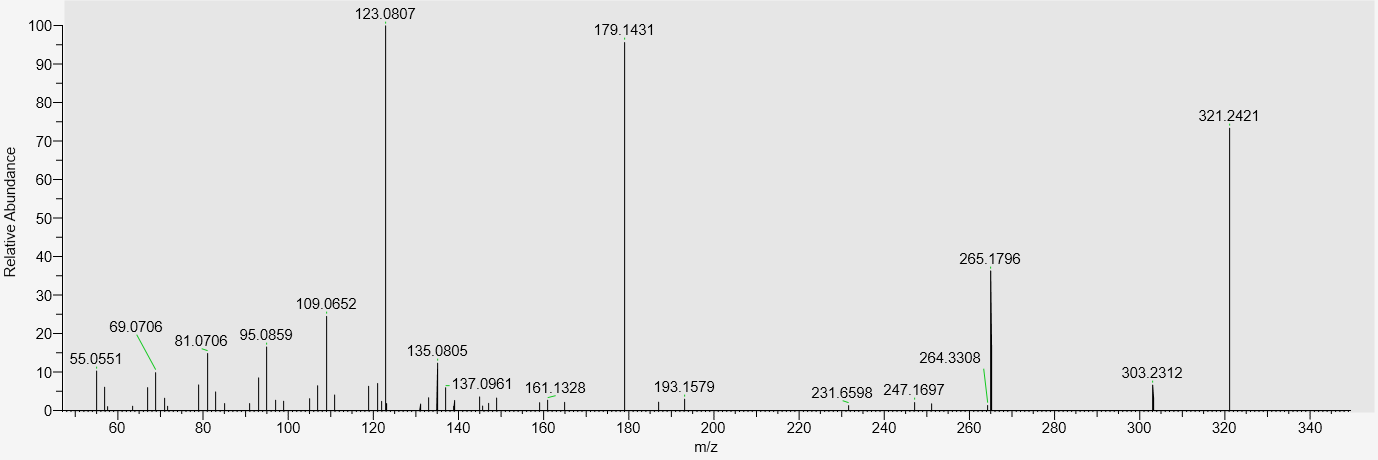  Substance X |

| MS2 277.1071, positive  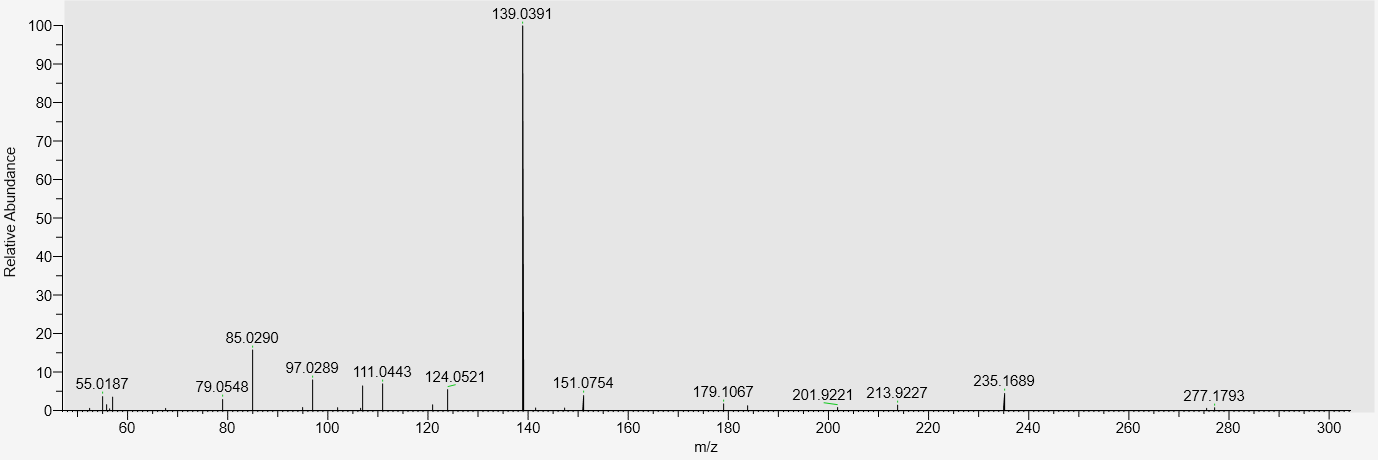  Substance XI | MS2 235.0604, negative  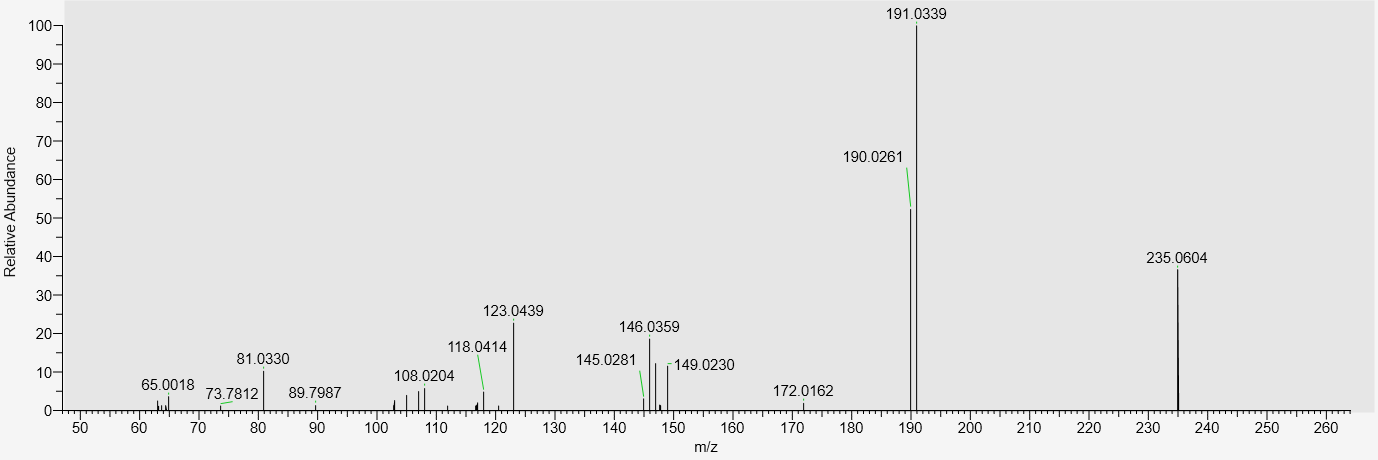  Substance XII |
| --- | --- |
| MS2 191.0336, negative  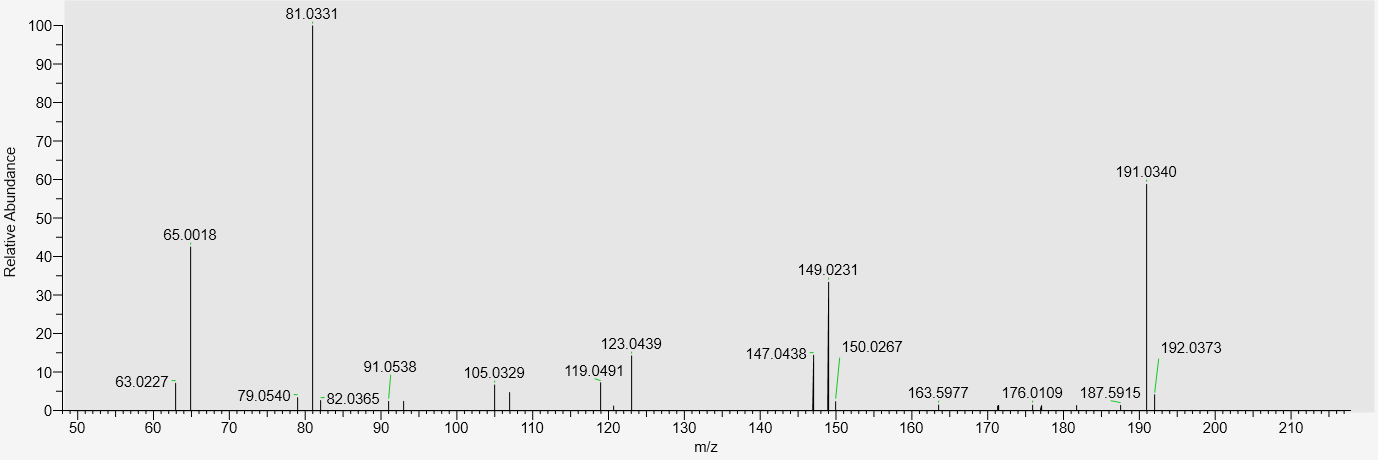  Substance XIII | MS2 193.0499, positive  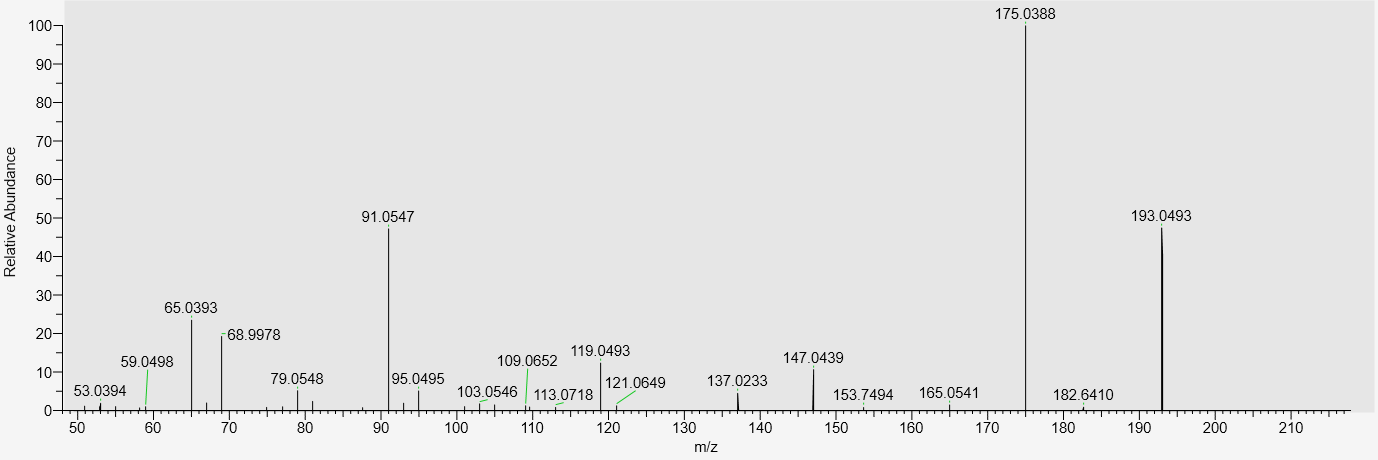 |
| MS2 317.0815, negative  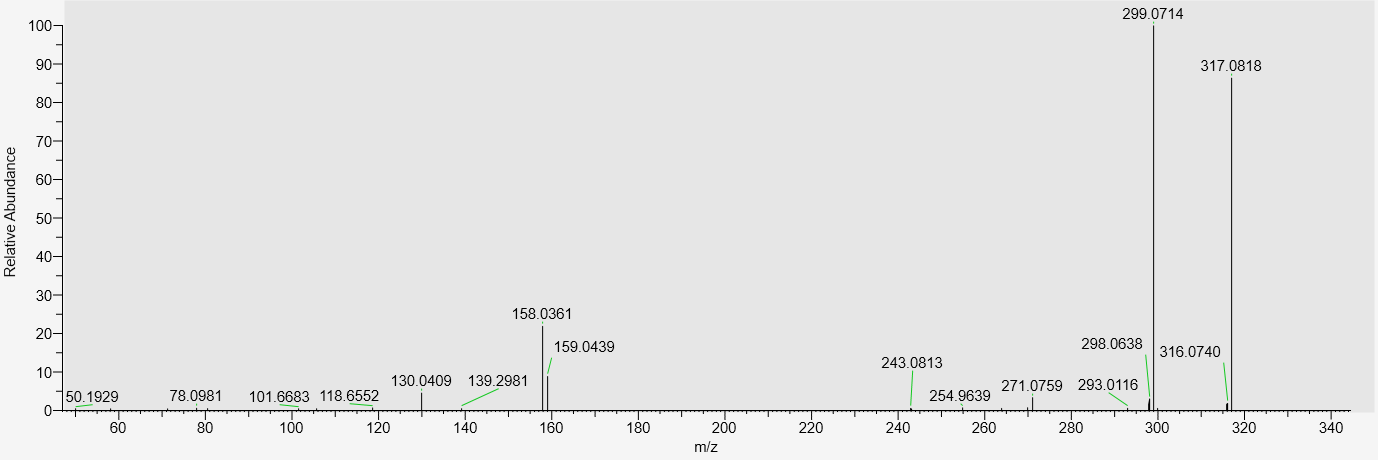  Substance XIV | MS2 387.1084, negative  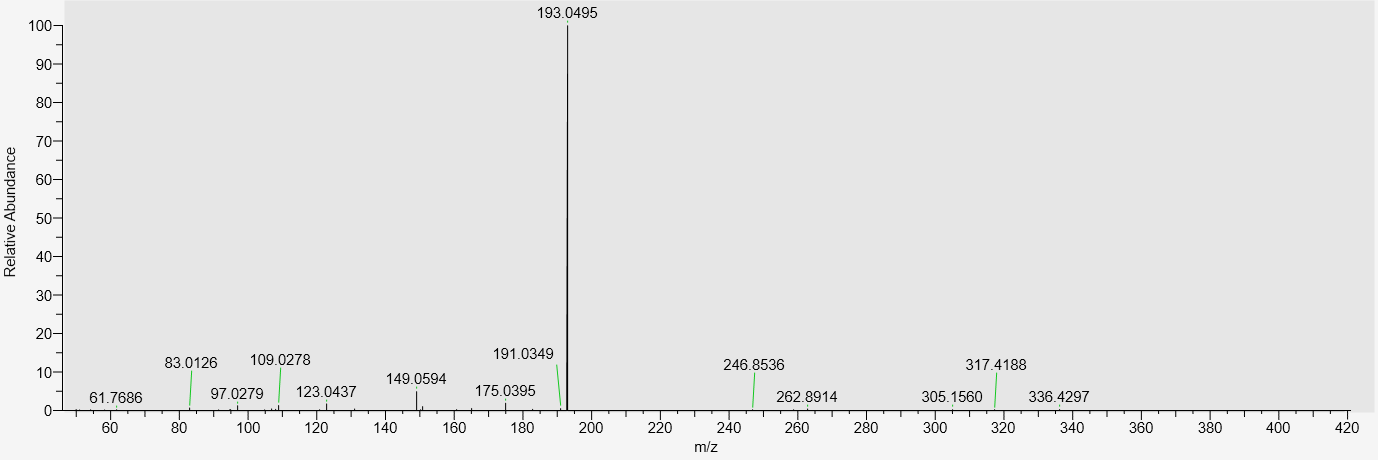  Substance XV |

| MS2 243.0878, positive  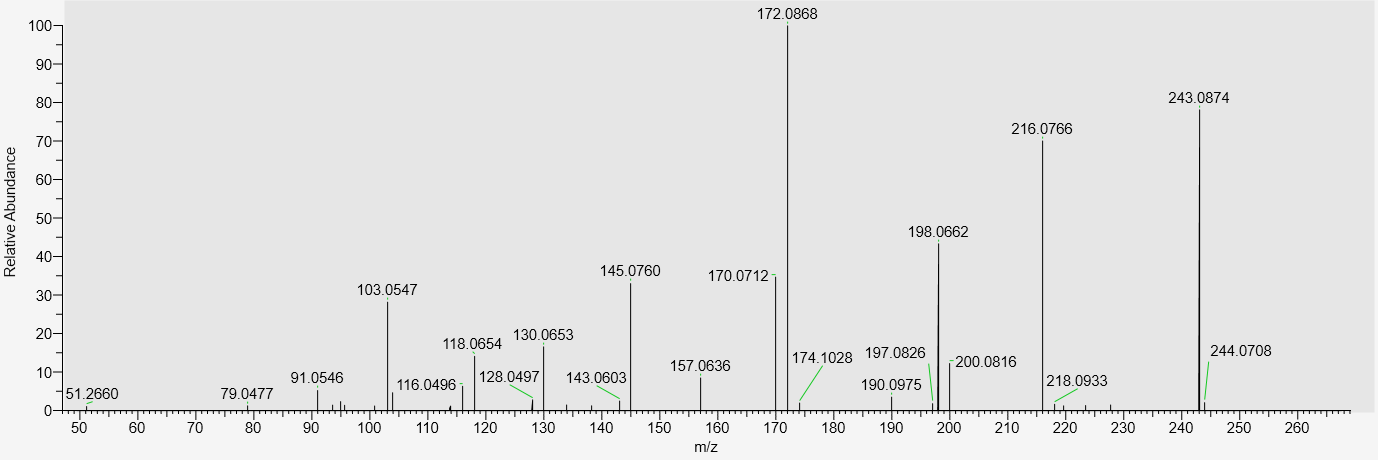  Substance XVI | MS2 367.2487, negative  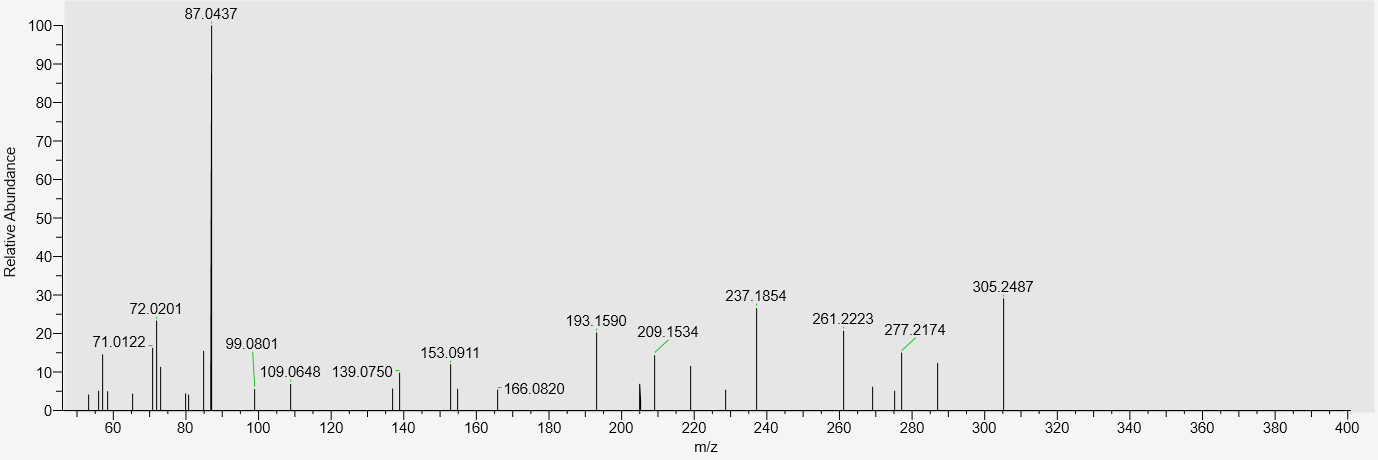  Substance XVII |
| --- | --- |
| MS2 133.0488, negative  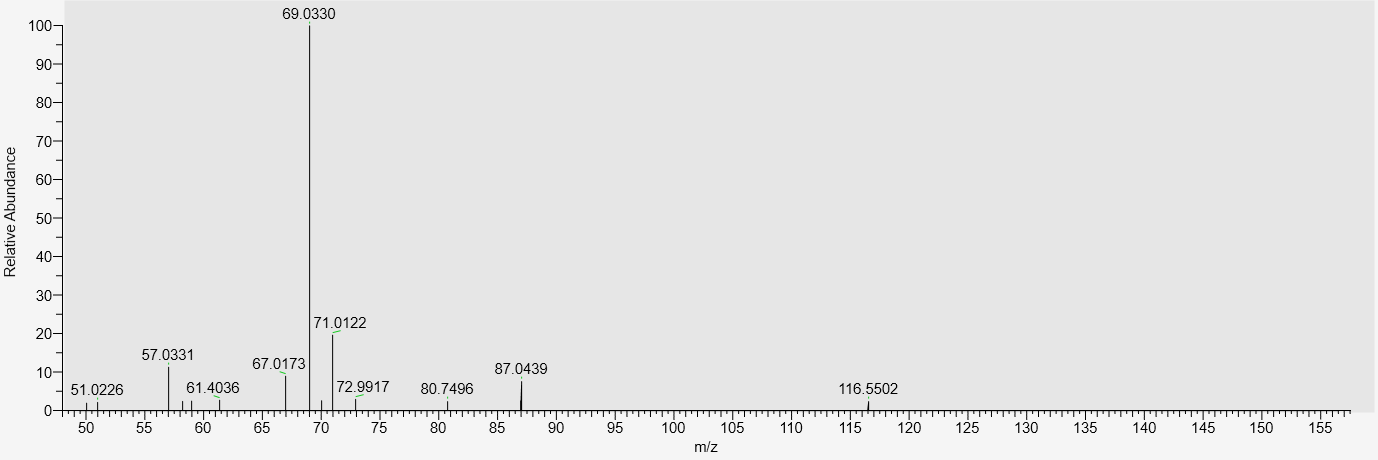  Substance XVIII | MS2 158.0811, negative  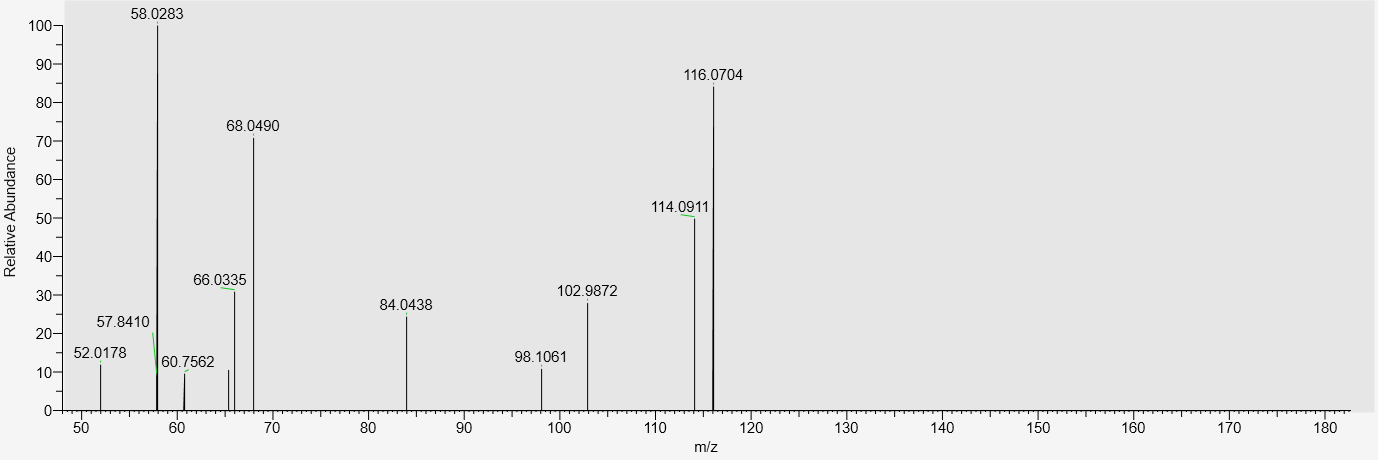  Substance XIX |
| MS2 153.0547, negative  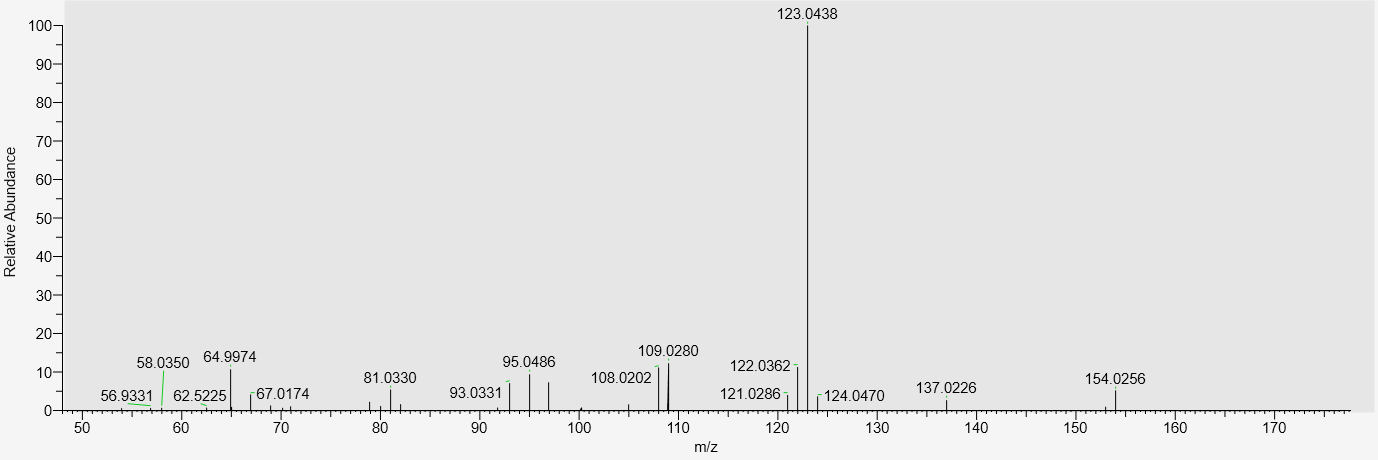  Substance XX | MS2 181.0495, negative  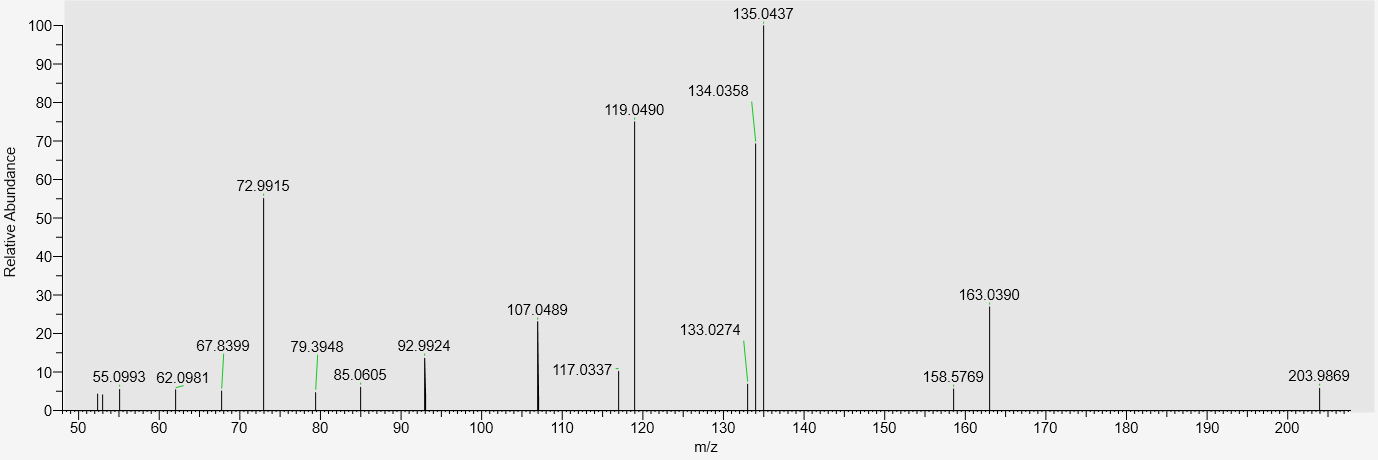  Substance XXI |

| MS2 165.0545, negative  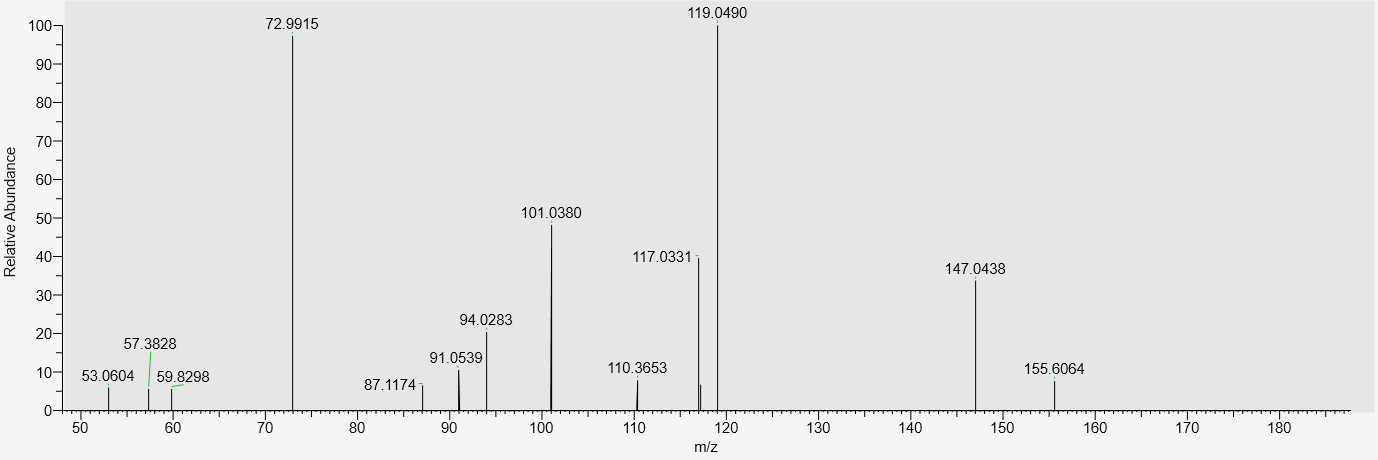  Substance XXII | MS2 335.2224, negative  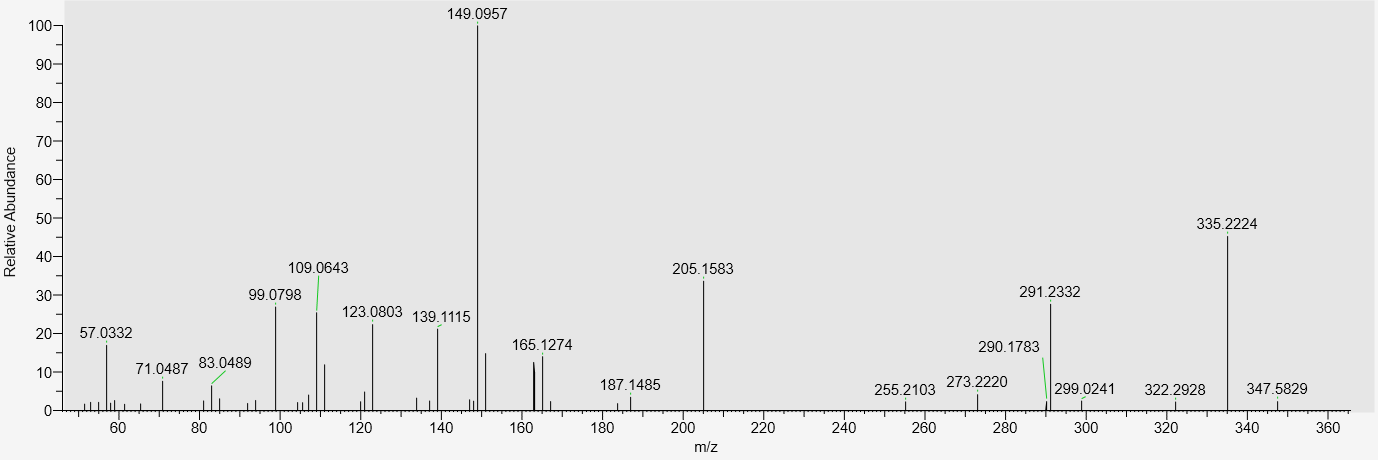  Substance XXIV |
| --- | --- |
| MS2 232.0606, negative  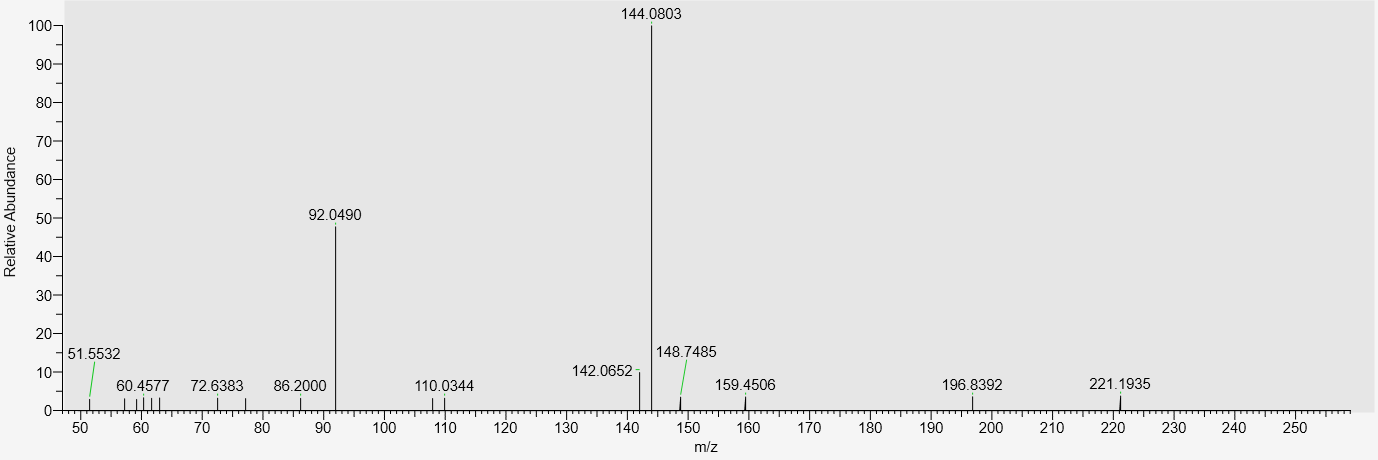  Substance XXIII | MS2 234.0759, positive  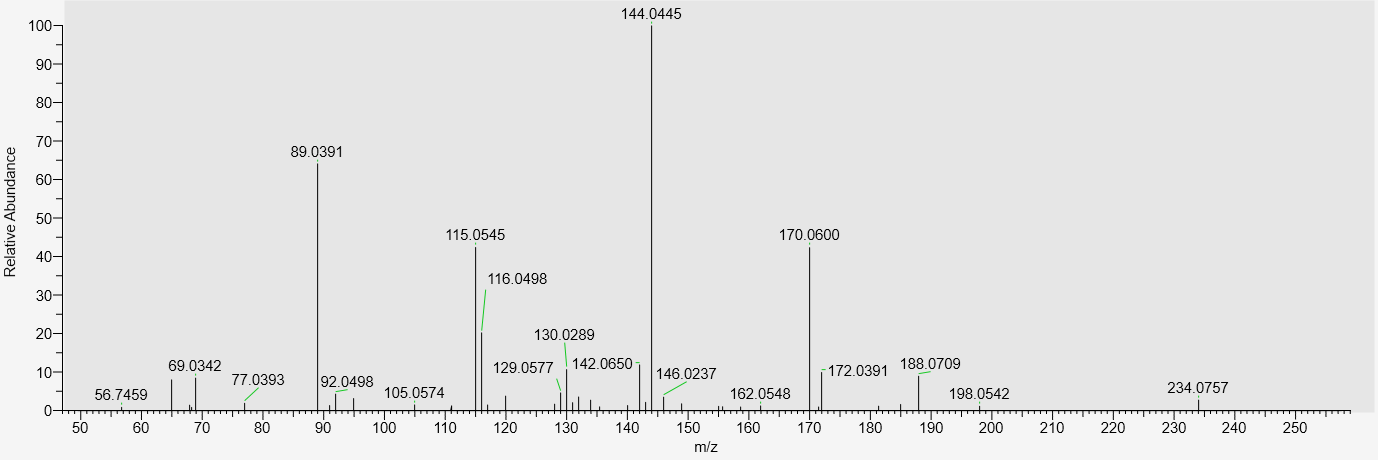 |
| MS2 547.3283, negative  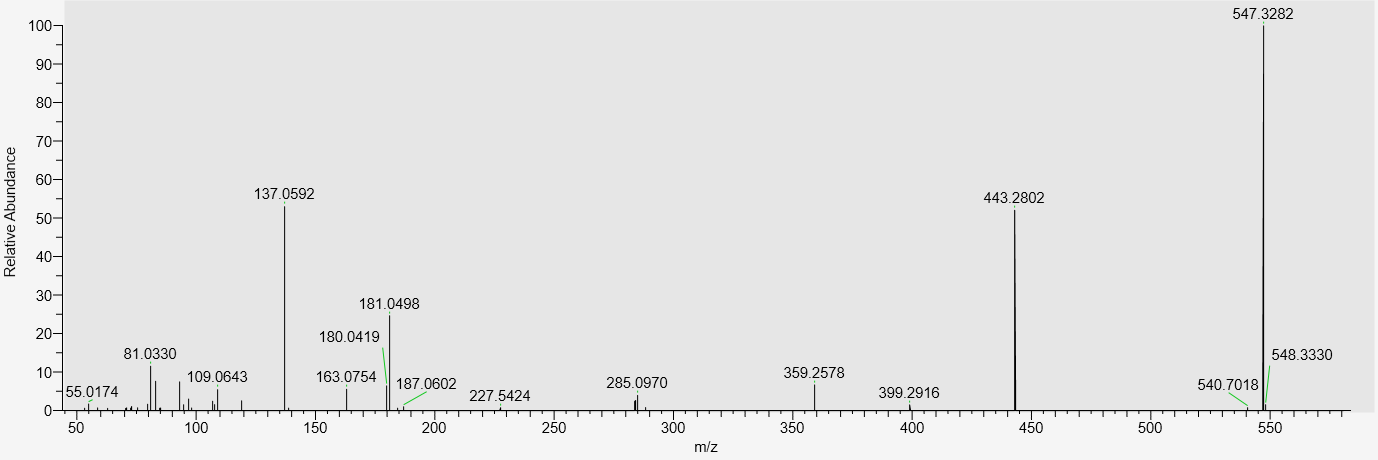  Substance XXV | MS2 549.3419, positive  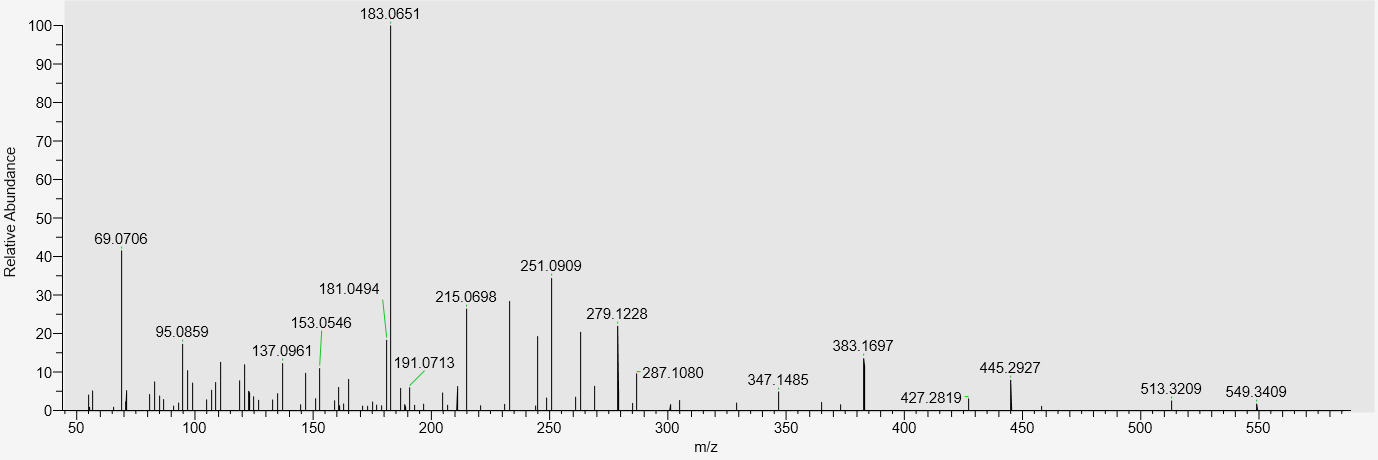 |

| MS2 605.3700, negative  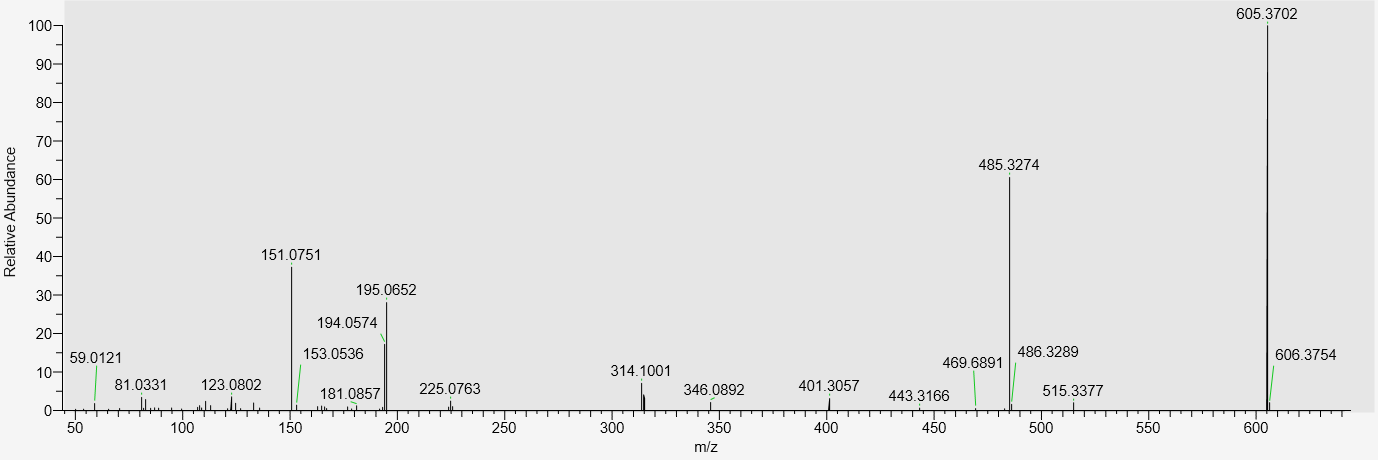  Substance XXVI | MS2 167.0703, positive  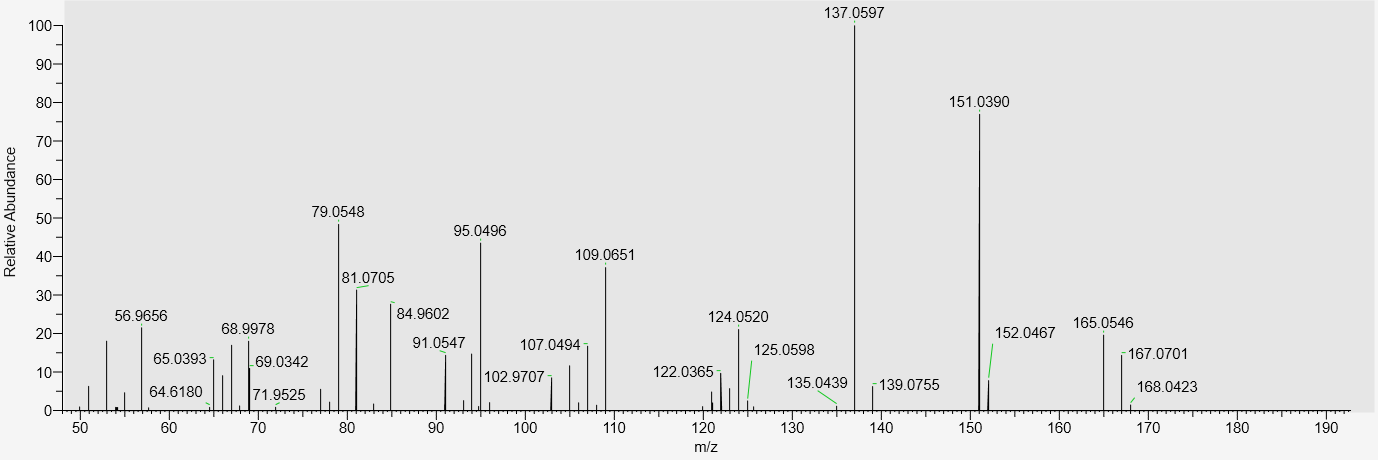  Substance XXVII |
| --- | --- |
| MS2 166.0864, positive  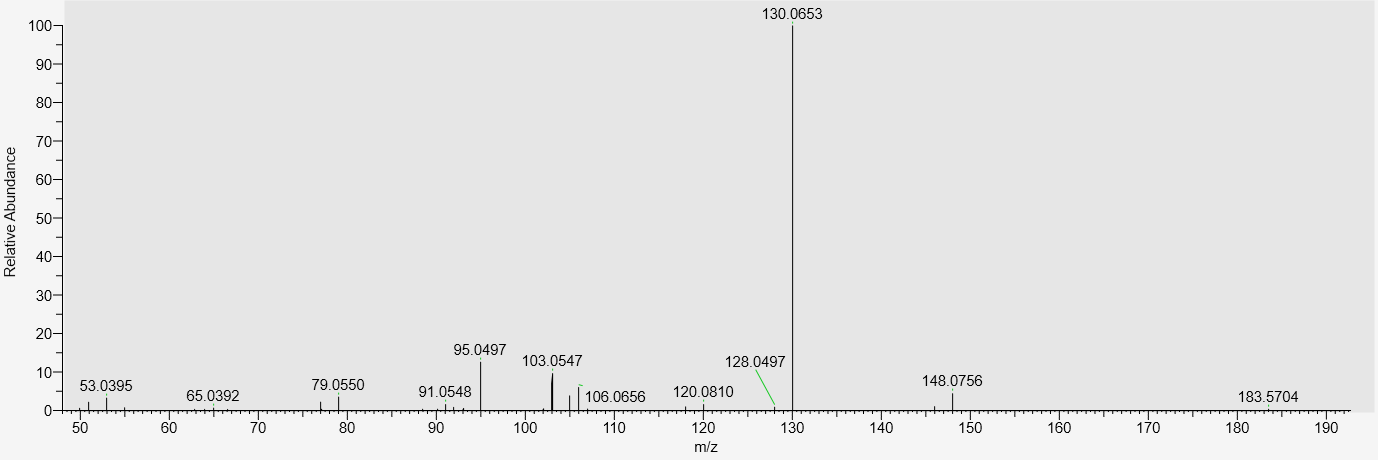  Substance XXVIII | MS2 577.3738, positive  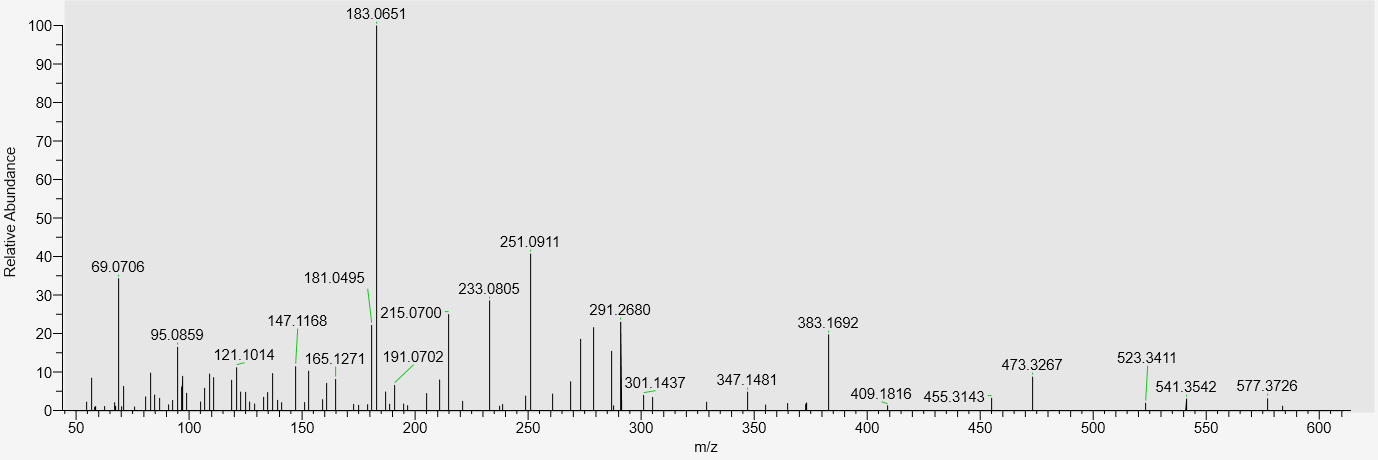  Substance XXIX |
